# Nucleation-dependent propagation of Polycomb modifications emerges during the *Drosophila* maternal to zygotic transition

**DOI:** 10.1101/2025.07.02.662854

**Authors:** Natalie Gonzaga-Saavedra, Eleanor A. Degen, Isabella V. Soluri, Corinne Croslyn, Shelby A. Blythe

**Affiliations:** Interdisciplinary Biological Sciences Graduate Program, Northwestern University, Evanston, IL, United States; Department of Molecular Biosciences, Northwestern University, Evanston, IL, United States; National Institute for Theory and Mathematics in Biology, Northwestern University and the University of Chicago, Chicago, IL, United States

## Abstract

During zygotic genome activation (ZGA) in *Drosophila*, broad domains of Polycomb-modified chromatin are rapidly established across the genome. Here, we investigate the spatial and temporal dynamics by which Polycomb group (PcG) histone modifications, H3K27me3 and H2Aub, emerge during early embryogenesis. Using ChIP-seq and live imaging of CRISPR-engineered GFP-tagged PcG components, we show that PRC2-dependent H3K27me3 accumulates adjacent to a subset of E(z)-bound prospective Polycomb Response Elements (PREs) beginning in nuclear cycle 14 (NC14), with patterns indicative of nucleation followed by spreading. Surprisingly, PRE-binding factors Pho, Combgap, and GAGA-factor are excluded from interphase nuclei prior to NC10 despite nuclear localization of E(z) throughout early interphases. Loss-of-function studies further demonstrate that GAGA-factor is largely dispensable for PcG domain establishment, whereas the pioneer factor Zelda is required for proper deposition of H3K27me3 and H2Aub at a subset of Polycomb domains. The role of Zelda at Polycomb domains is context-dependent; a subset of targets requires Zelda not for E(z) recruitment, but instead to license an E(z)-loaded PRE to deposit H3K27me3. Our findings support a model where licensing of PcG domains is an initial step in the regulatory processes governing Polycomb-regulated developmental genes.

**Impact Statement:** Epigenomics and quantitative imaging are used to investigate the re-establishment of histone modifications associated with the Polycomb group of epigenetic regulators during the *Drosophila* maternal-to-zygotic transition.

## Introduction

During the maternal to zygotic transition, the genomes of two highly specialized cells, sperm and egg, are epigenetically reprogrammed in order to support the development of the zygote. Following an initial period of post-fertilization mitotic proliferation, embryos undergo genome activation and initiate zygotic gene regulatory programs against the background of the reprogrammed epigenetic landscape. Many of the genes involved in cell fate specification and pattern formation become associated with chromatin modifications deposited by the Polycomb group (PcG), whose function is ultimately necessary for maintaining these genes in a heritable, transcriptionally silenced state. While the mechanism for PcG function for maintenance of silenced states has been intensely studied, the rules whereby genomic loci are selected for Polycomb silencing, and the mechanisms for establishing these genomic compartments in early embryogenesis remain unclear.

PcG factors associate in multi-subunit complexes that carry out distinct steps required for the maintenance of long-term repression (Schuettengruber et al., 2017; Laugesen et al., 2019). Polycomb repressive complexes 1 and 2 (PRC1 and 2) mediate the reading and writing of histone post-translational modifications at the heart of the PcG system (Müller et al., 2002; Shao et al., 1999). PRC2 mediates methylation of Histone H3 lysine 27 (H3K27) through the histone methyltransferase subunit Enhancer of Zeste (E(z)) (Cao et al., 2002; Czermin et al., 2002). Tri-methylation of H3K27 (H3K27me3) deposited by E(z) can be ‘read’ by additional PcG factors to allosterically enhance E(z) catalysis and to promote the spreading of H3K27 methylation across broader genomic regions which can span tens to hundreds of kilobase-pairs (Margueron et al., 2009; Lee, Holder, et al., 2018; Lee, Yu, et al., 2018). The ubiquitylation of Histone H2A (H2Aub) by the Sce/dRing1 component of PRC1 can also lead to allosteric enhancement of E(z) catalysis (Kalb et al., 2014; Kasinath et al., 2021). In *Drosophila*, targeting of PRC1 and 2 to sites designated for PcG regulation is driven in part by *cis-*regulatory sequences termed Polycomb Response Elements (PREs) which are bound by several sequence-specific *trans*-acting factors including but not limited to Pleiohomeotic (Pho), Pho-like (Phol), Combgap (Cg), and GAGA-factor (GAF) (Hagstrom et al., 1997; Brown et al., 1998; Horard et al., 2000; Brown et al., 2003; L. Wang et al., 2004; Klymenko et al., 2006; Ray et al., 2016; Brown et al., 2018). In proliferating cells, functional PRE sequences are necessary for maintaining PcG states across successive cell divisions (Coleman & Struhl, 2017; Laprell et al., 2017), supported by allosteric enhancement from these read-write feedback loops.

Early embryos present several challenges for the establishment and maintenance of PcG modifications. *Drosophila* embryos initially undergo thirteen rapid, metasynchronous nuclear divisions before undergoing cell cycle pause during nuclear cycle 14 (NC14, cellular blastoderm stage) and completing zygotic genome activation (ZGA) (Foe & Alberts, 1983). Over the cleavage period, embryos gradually establish patterns of chromatin accessibility that will sustain the initial rounds of transcriptional activity that drive developmental patterning (Blythe & Wieschaus, 2016). These patterns of accessibility are determined in whole or in part by specialized transcription factors termed pioneer factors that operate at distinct timepoints to gradually install the initial chromatin accessibility state (Liang et al., 2008; Schulz et al., 2015; Sun et al., 2015; Gaskill et al., 2021; Duan et al., 2021; Harrison et al., 2023). Genome-wide assessments of H3K27me3 deposition in early *Drosophila* embryos have yielded two distinct models for the establishment of PcG modifications. In one case, H3K27me3 modifications are observed to arise *de novo* only during mid nuclear cycle 14 (NC14), which is coincident with the acquisition of substantial chromatin accessibility and cell cycle lengthening that accompanies ZGA (Chen et al., 2013; Li et al., 2014; Reinig et al., 2020). In another case, patterns of H3K27me3 deposition have been argued to be transmitted to offspring through the maternal germline, with retention of H3K27me3 at some level throughout the cleavage divisions (Zenk et al., 2017; Cardamone et al., 2025). We wished to investigate further the discrepancy between these observations and approach the problem from the mechanistic standpoint of existing models for PcG maintenance. In the following, we have addressed the timing of PcG modifications over the course of *Drosophila* ZGA, with a specific focus on the availability of PRE-binding factors, and the relationship of PcG modification deposition to pioneer factors known to drive establishment of the initial zygotic chromatin state.

## Results

### Rapid Establishment of Polycomb histone modifications at prospective Polycomb Response Elements during ZGA

To investigate the kinetics of establishment of PcG modification of the zygotic genome, we performed ChIP-seq in cellular blastoderm stage (NC14) embryos for PcG-dependent histone modifications (H3K27me3,-me1, and H2Aub, Figure 1). To identify loci where deposition of PcG modifications could be expected, we also measured the blastoderm stage enrichment of the PRC2 component E(z), as well as the core PRE component Pho, additional PRE binding factors, Combgap (Cg) and GAGA-Factor (GAF), and the pioneer factor Zelda (Figure 1A, Zelda ChIP-seq data from (Harrison et al., 2011)). To measure E(z), Cg, GAF, and Pho binding, we performed ChIP-seq for each individual factor on chromatin derived from embryos expressing a GFP chimera for the corresponding factor generated by CRISPR/Cas9 genome editing (see Materials and Methods). The engineered EGFP-E(z), EGFP-Cg, EGFP-GAF, and Pho-sfGFP chimeric alleles are all viable and fertile as homozygotes. Measurement of these factors identifies a set of 4576 E(z) peaks that represent prospective targets of PcG regulation, only a subset of which have been functionally validated as PREs. As expected, E(z) peaks also often correspond to binding sites for either Pho, GAF and/or Cg (Figure 1A). Overall, 86.6% (n = 3966) E(z) peaks overlap with peaks of Pho, Cg, or GAF in different combinations of factors. PcG-modified domains are nucleated by recruitment of PRC1/2 to PREs, resulting in local modification of neighboring nucleosomes and subsequent spreading of modifications across topologically delimited segments of the genome. To estimate the extent of these broader regions of H3K27me3, we used a Hidden Markov Model approach (STAN) to identify a set of PcG “domains” from our late NC14 H3K27me3 dataset as described previously (Zacher et al., 2017; Bonnet et al., 2019). In blastoderm stage chromatin we identify 255 broad PcG domains in total with median length of 19.2 kb (range: 4.2 to 350 kb). Of these PcG domains, 93% (n = 237) contain one or more E(z) peaks. Notably, not all E(z) peaks are found in PcG domains: 72.3% (n = 3312) of E(z) peaks do not associate with above-threshold H3K27me3 ChIP-seq signal by the end of NC14 (Figure 2). In general, E(z) peaks outside of PcG domains were found to localize to the transcriptional start sites in promoters of genes decorated with RNA Polymerase 2 phosphorylated at position 5 of the C-terminal domain heptapeptide repeat as well as promoter-proximal Histone H3 Lysine 4 mono-, di-, and tri-methylation and Histone H3K27 acetylation (Figure 2 and Figure 2-figure supplement 1). On average, E(z) and Pho binding within PcG domains yields higher ChIP-seq enrichment, and these show evidence of long-distance “spreading” (Figure 2B). Scoring occurrences of DNA binding factors at E(z) peaks within PcG domains, only Pho and Zelda demonstrate significant enrichment within PcG domains, whereas Cg and GAF are more enriched at E(z) peaks outside of PcG domains. Based on these observations, the set of E(z) peaks can be classified into at least two categories based either on occurrence within a PcG domain, or outside of the domain. We chose to focus on the characteristics of E(z) peaks within PcG domains.

**Figure 1:**
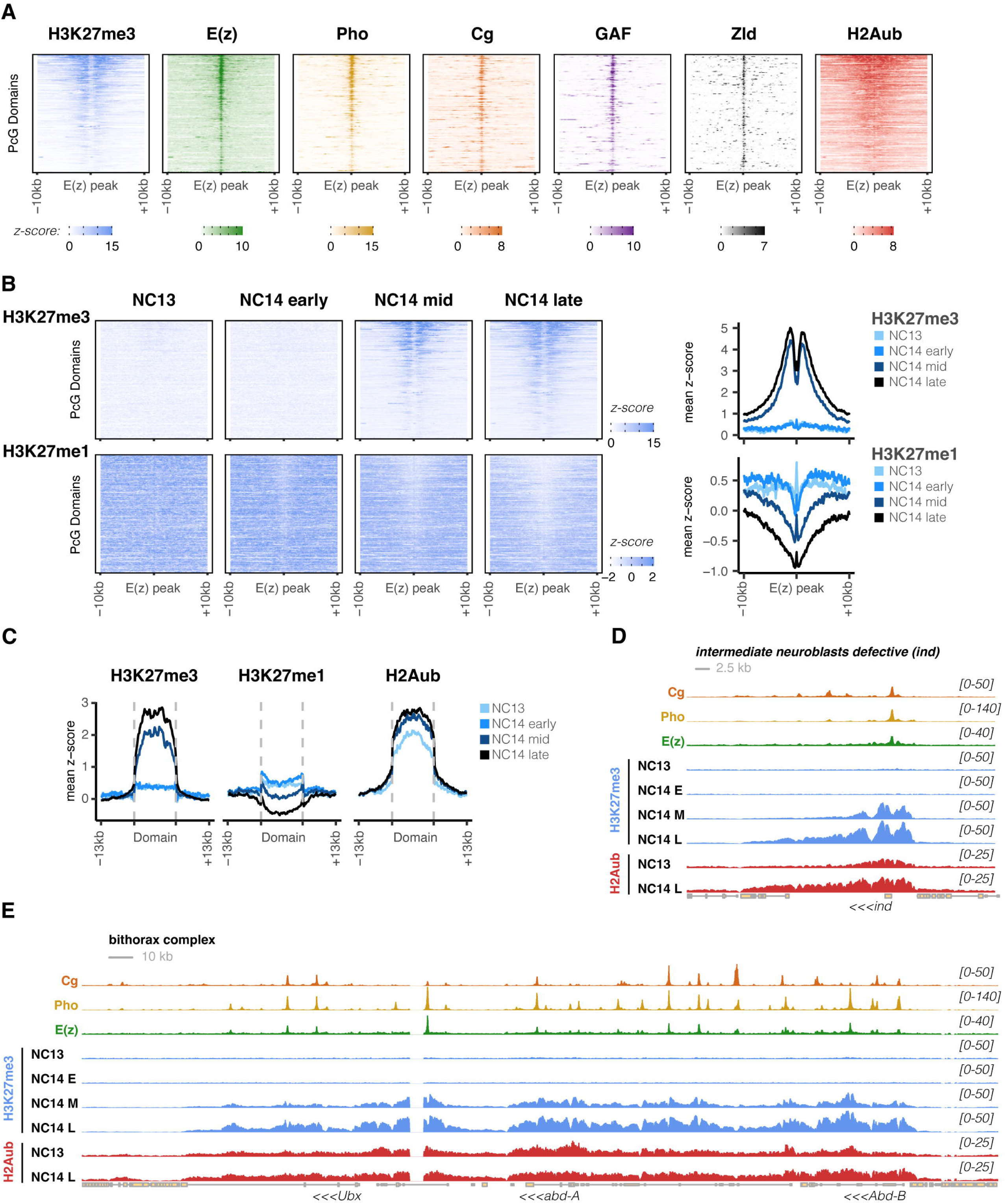
Timing of establishment of PcG modifications at ZGA. **A)** Heatmaps are plotted for ChIP-seq measurements in cellular blastoderm staged embryos for the indicated factors (top) over a set of E(z) peaks localized within PcG Domains, flanked by ± 10 kb. One representative E(z) peak per domain was selected for plotting. Heatmap rows are identically ordered from top to bottom by overall average H3K27me3 intensity per peak. E(z), Pho, Cg, and GAF ChIP-seq are all performed using an anti-GFP antibody on CRISPR-engineered embryos expressing GFP-tagged chimeric alleles of the target protein. Zld ChIP-seq was previously reported (Harrison et al., 2011). Data were standardized prior to plotting and colorbars (bottom) indicate binding intensity in units of standard deviations. **B)** Heatmaps are plotted for standardized ChIP-seq measurements in embryos collected at the indicated stages (top) for either H3K27me3 (top row), or H3K27me1, bottom row. The average standardized ChIP-seq signal per time point is plotted at right. **C)** The average standardized ChIP-seq signal for timecourse measurements of H3K27me3,-me1, and H2Aub are plotted over entire PcG domains. The domains are first divided into 100 bins and standardized ChIP-seq signal was averaged per bin prior to plotting average signal across domains ± 13 kb flanking sequence. **D)** Visualization of the PcG domain containing the *ind* locus and counts-per-million (CPM) normalized ChIP-seq measurements for Cg, Pho, E(z) at cellular blastoderm, and timecourse measurements of H3K27me3 (NC13 - NC14 late), and H2Aub (NC13, NC14 late). ChIP target is indicated at left and the display range of the CPM normalized ChIP data is indicated at right. Scale bar: 2.5 kb. **E)** Visualization of the PcG domain containing the bithorax complex for the same CPM-normalized ChIP tracks as in panel D. Scale bar: 10 kb.

**Figure 2:**
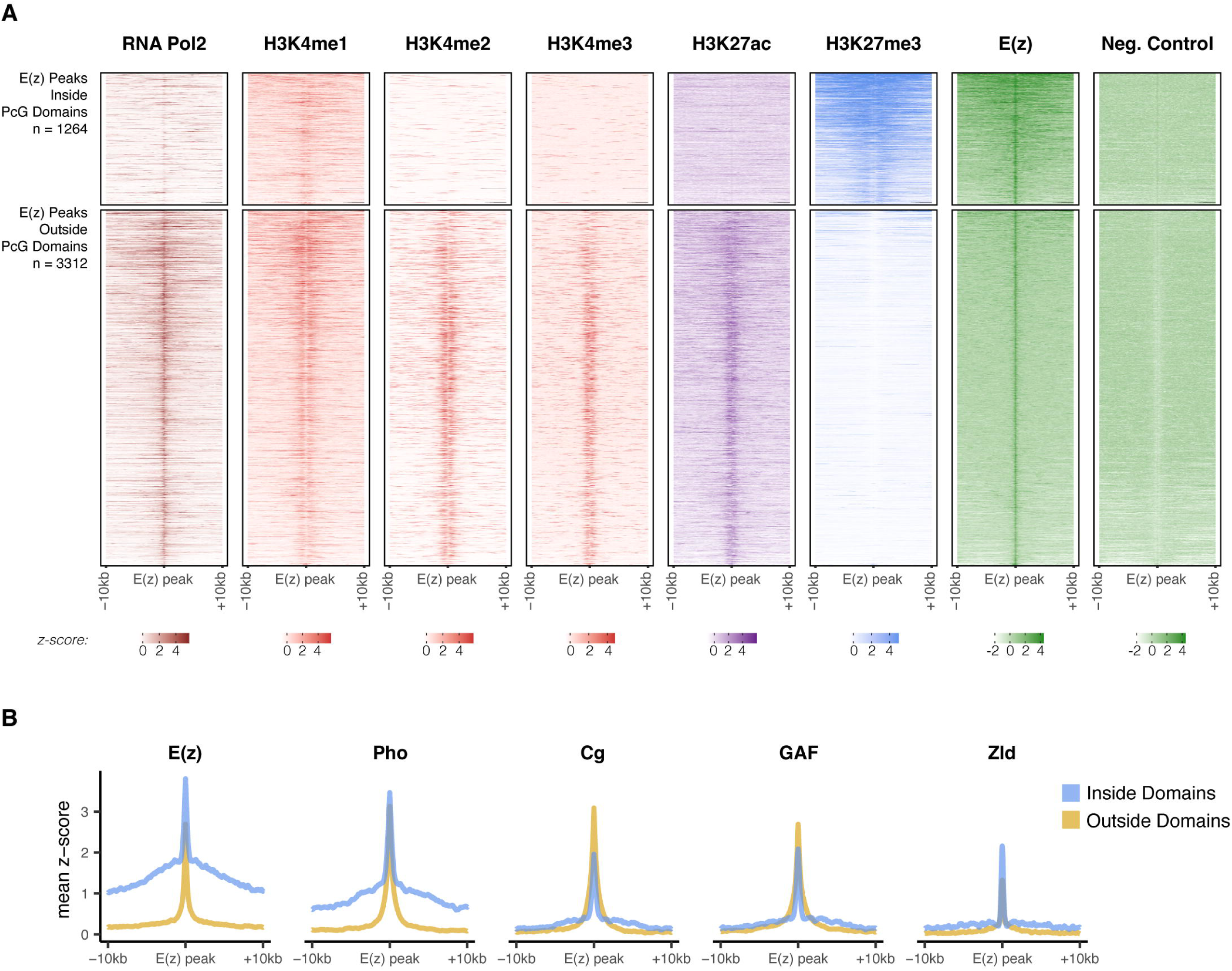
E(z) peaks outside of PcG Domains generally associate with active promoters. **A)** Heatmaps for the indicated ChIP targets were plotted centered over all 4576 E(z) peaks (± 10 kb) and subset by membership within a PcG domain. Data were z-scored prior to plotting, and the plotted intensity range is indicated by the colorbar beneath each respective heatmap. Peaks are ordered on the y-axis by the overall intensity of E(z) binding in the region shown. A negative control ChIP (IP for GFP on wild type chromatin) is shown at the far right. While E(z) peaks within domains are enriched for H3K27me3, peaks outside of domains show a high degree of correlation with marks of active transcription including RNA Pol2 (CTD pSer5), H3K4me1/2/3, and H3K27 acetylation. **B)** The mean enrichments of E(z) and DNA binding factors over E(z) peaks ± 10 kb are plotted for peaks inside (blue) and outside (yellow) of PcG domains.

Consistent with prior reports (Chen et al., 2013; Li et al., 2014; Reinig et al., 2020), we observe that H3K27me3 is strongly detected by mid-to late-NC14 adjacent to E(z) peaks and more broadly across PcG domains but is not strongly detected on NC13 and early NC14 chromatin (Figure 1A-C). This pattern of accumulation could be due either to limited access of PcG components to chromatin prior to NC14 or could reflect the expected slow kinetic maturation of sub-trimethylated histone species near sites of recruitment for PcG complexes (Sneeringer et al., 2010; Zheng et al., 2012; Reinig et al., 2020; Lundkvist et al., 2023). To distinguish these possibilities, we also performed ChIP-seq against H3K27me1 and observed a broadly distributed signal surrounding E(z) peaks at NC13 and early NC14 that is gradually depleted from sites that acquire H3K27me3 by mid-and late-NC14 (Figure 1B). The broad genome-wide distribution of H3K27me1, including in regions outside of canonical PcG domains, resembles profiles previously measured in mammalian tissue culture (Ferrari et al., 2014). At the domain scale, H3K27me1 abundance is on average higher within domains than in flanking regions during NC13 and early NC14, and this average signal is incrementally reduced at mid-and late-NC14 (Figure 1C). The anti-correlation between H3K27me1 and-me3 within domains is consistent with a mechanism where the rapid rate of cleavage stage cell cycles precludes the maturation of H3K27 residues to a fully trimethylated state until mid-to late-NC14, and that PcG factors access chromatin during cleavage divisions to catalyze sub-trimethylated states. To further evaluate this model, we tested for evidence of PRC1 interactions with PcG domains, performing ChIP-seq at NC13 and NC14 (mid-and late-) against H2Aub, catalyzed by the Sce/dRing component of PRC1. In contrast to the H3K27 modification states, H2Aub is readily detected near E(z) peaks and within PcG domains as early as NC13 (Figure 1A and C). These results suggest that both PRC2 and PRC1 access and modify *Drosophila* chromatin prior to large-scale ZGA despite rapid cell cycle progression.

Examination of the patterns with which H3K27me3 comes to be deposited at individual domains over the course of NC14 suggests that emergence of this state is produced by a “nucleation followed by spreading” mechanism that initiates at E(z) peaks within PcG domains, which we will refer to as prospective PREs (pPRE). Figure 1D shows an example of a representative moderate-sized PcG domain containing the early zygotic gene *intermediate neuroblasts defective* (*ind*). The major pPRE in this domain overlaps the *ind* transcription start site, and overlaps with peaks for E(z), Pho, and Cg. The earliest accumulation of H3K27me3 observed at mid-NC14 is always flanking the pPRE and thereafter is seen to ‘spread’ distally. H2Aub in general at equivalent timepoints demonstrates a greater degree of spreading over the PcG domain distal to sites of presumptive nucleation compared with H3K27me3, suggesting that the ubiquitylation rate of Sce outpaces the trimethylation rate of E(z). Similar kinetics are observed over the PcG domain corresponding to the bithorax complex (Figure 1E). Compared with *ind*, the bithorax complex contains numerous sites of nucleation, however, despite the increased density of nucleation sites, H3K27me3 accumulation at the bithorax complex shows a similar delay until mid-NC14.

### PRE-binding factors are excluded from nuclei during early cleavage stages

If pPREs were to serve as sites for maintenance of maternal PcG modification states, then effector proteins should localize to nuclei during the cleavage divisions. Protein expression kinetics for PcG components including E(z) and Pho have previously been measured using transgenically expressed fluorescent protein chimeras driven by heterologous alpha-tubulin regulatory sequences (Steffen et al., 2013). However, this study did not address whether nuclear concentration of these factors is consistent over the cleavage divisions. Whereas some maternally supplied proteins such as Zelda readily localize to the nucleus throughout the cleavage divisions (Harrison et al., 2011; Nien et al., 2011), other key factors, such as Polycomb, Dorsal, Bicoid, TATA-binding protein, Heat Shock Factor, and the large subunit of RNA Polymerase I are subject to regulated nuclear import and/or translation during these stages (Rushlow et al., 1989; Z. Wang & Lindquist, 1998; Falahati et al., 2016; Reinig et al., 2020; Gregor et al., 2007), demonstrating changes in the fraction of nuclear localized protein as embryos approach ZGA.

We wished to reproduce and extend these prior measurements using endogenously tagged PcG components and therefore we performed live imaging of our GFP chimeras to quantify relative GFP intensity for each chimera spanning from interphase of NC10, when nuclei first migrate to the embryonic cortex, through NC14, when embryos begin gastrulation (Figure 3). EGFP-E(z) localizes to the nucleus during interphases 10-14 and, consistent with previous reports (Steffen et al., 2013), is evicted from chromatin during mitosis (Figure 3A, Figure 3-video 1). However, E(z) concentration decreases in each subsequent interphase, reaching a minimum observed nuclear concentration at NC14 when H3K27me3 accumulates on chromatin. Such high initial concentrations could potentially facilitate rapid maturation of H3K27 through consecutive methyl-states despite an expected slow catalytic rate for the me2 to me3 modification (Sneeringer et al., 2010; Zheng et al., 2012).

**Figure 3:**
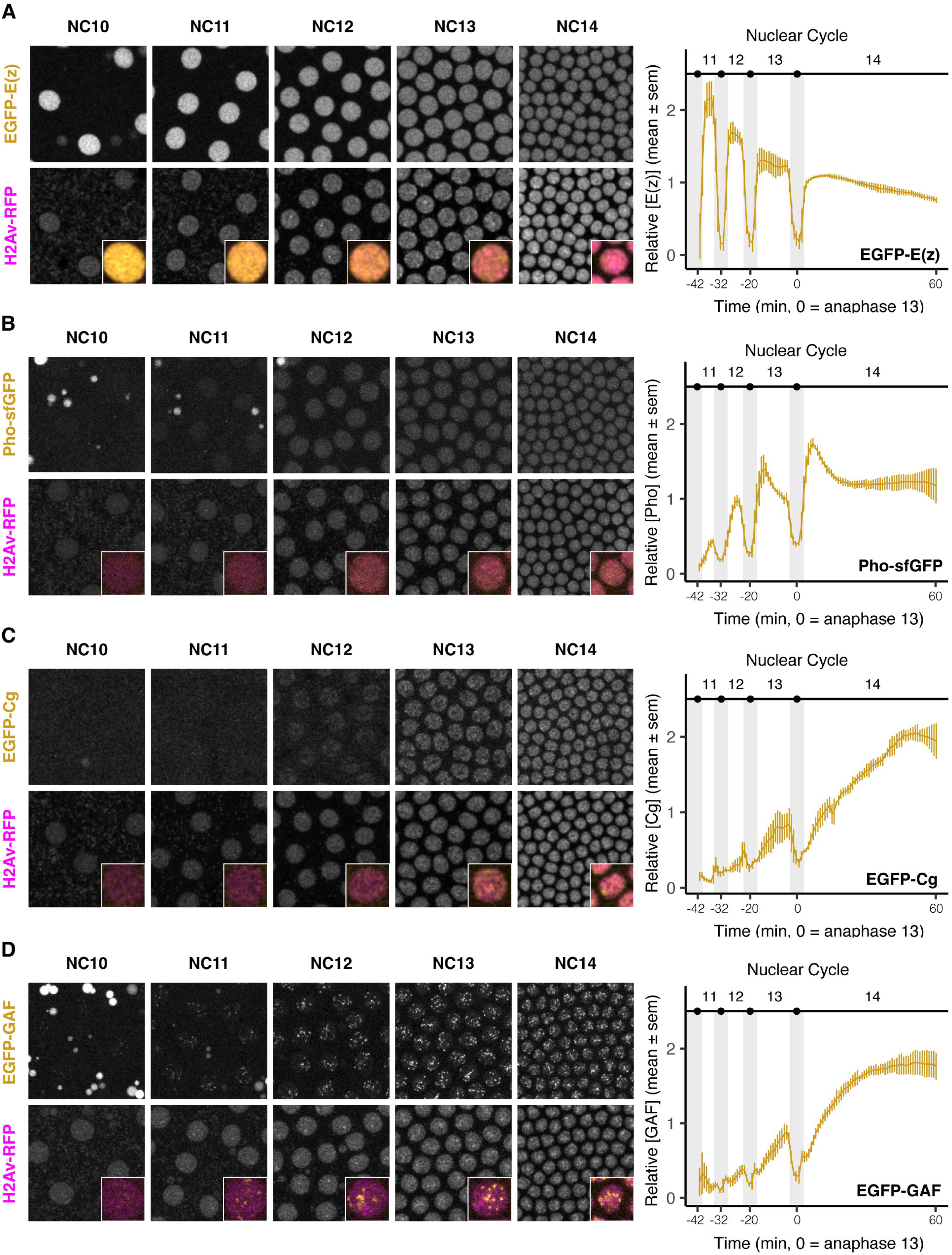
Nuclear localization patterns of E(z) and three PRE-binding factors over late cleavage divisions. EGFP-E(z) (**A**), Pho-sfGFP (**B**), EGFP-Cg (**C**), and EGFP-GAF (**D**) were subjected to time-lapse confocal microscopy from nuclear migration (NC10) through the period of cellularization (NC14 + 60 minutes). All embryos also expressed Histone H2Av-RFP to visualize nuclei. The images at left show maximum-projected still images from representative time-lapse movies for the indicated GFP chimera (top) and His2Av-RFP (bottom) at interphase of the indicated nuclear cycles. The insets show a color merge of one nucleus at each respective nuclear cycle (yellow: GFP chimera, magenta: His2Av-RFP). The quantifications at right show the relative nuclear concentration of each GFP chimera from NC11 through NC14 (top x-axis). Grey vertical bars indicate the approximate mitotic periods for each nuclear cycle. Data represent averaged intensity measurements from n ≥ 3 embryos per chimera, mean normalized ± standard error of the mean. While E(z) readily localizes to nuclei throughout the cleavage divisions, the three PRE-binding factors Pho, Cg, and GAF increase nuclear concentration over the last nuclear divisions before cell cycle lengthening at NC14.

In contrast, the three measured PRE-binding factors Pho-sfGFP, EGFP-Cg, and EGFP-GAF gradually increase nuclear concentration between NC11 and NC14 (Figure 3 B-D) reaching maxima during NC14. Pho-sfGFP is detected in nuclei during interphase, beginning at a low but detectable level at NC11 and increasing markedly at NC13 and 14 (Figure 3B). Consistent with a prior report, Pho-sfGFP is not strongly detected on mitotic chromatin (Figure 3-video 2) (Steffen et al., 2013). EGFP-Cg demonstrates a more gradual increase in nuclear concentration than Pho-sfGFP over this period, starting with lower apparent concentration at NC11, and continually increasing in nuclear concentration throughout NC14 (Figure 3C). Unlike Pho and E(z), Cg is detected on mitotic chromatin, albeit at a lower intensity than during interphase (Figure 3-video 3). Finally, consistent with previous reports, EGFP-GAF localizes from NC10-14 to bright apical puncta corresponding to simple (AAGAG)_n_ satellite repeats (Raff et al., 1994; Gaskill et al., 2021, 2023; Dima & Reeves, 2025), and nucleoplasmic concentrations of GAF gradually increase over this period, reaching a maximal intensity at NC14 (Figure 3D, Figure 3-video 4). These observations indicate that these three PRE-binding factors are effectively absent from early cleavage-stage chromatin and significantly increase their nuclear concentration over the timeframe of ZGA.

One limitation of live-imaging of GFP-tagged chimeras are potential discrepancies between protein expression and GFP maturation rates, which could lead to underestimates of protein expression levels. To validate the live-imaging measurements, we performed immunostaining on formaldehyde-fixed preparations of our endogenously GFP-tagged E(z), Pho, Cg, and GAF chimeric embryos (Figure 3-figure supplement 1). In accordance with the live imaging results, E(z) was found to localize to the nucleus at all observed nuclear cycle interphases, maintaining an apparent high concentration prior to nuclear migration and blastoderm formation at NC10 (Figure 3-figure supplement 1A). Also in accordance with the live imaging measurements is the observation that neither Pho, Cg, nor GAF (Figure 3-figure supplement 1B-D) achieve above-background nuclear concentrations until NC11 (Pho) or NC12 (Cg, GAF). Taken together, our live and fixed embryo analysis of E(z), Pho, Cg, and GAF nuclear localization indicates that while E(z) readily localizes to nuclei throughout the cleavage period, at least three PRE binding factors, Pho, Cg, and GAF show only limited nuclear localization until just prior to large-scale ZGA. Although supra-NC14 concentrations of E(z) are observed within nuclei during all syncytial interphases, key factors for mediating nucleation of PcG domains during maintenance phases are apparently absent from chromatin until just before H3K27me3 becomes detectable by ChIP at NC14.

### H3K27me3 and H2Aub are not maintained on chromatin at high levels during cleavage divisions

To determine to what degree PcG-dependent chromatin modifications are maintained on cleavage stage chromatin, we next performed immunostaining for PcG-dependent histone modifications H3K27me3 and H2Aub (Figure 4). By this approach, we confirm and reproduce the previously reported observation of heterogeneous staining of H3K27me3 in zygotic nuclei shortly after fertilization (Figure 4A, NC2) (Zenk et al., 2017). Both nuclei in Figure 4A show intense anti-H3K27me3 immunoreactivity over approximately half of each nucleus, and prior observations trace the source of this heterogeneity to the maternal pronucleus, which is marked with significant amounts of H3K27me3, in contrast to the paternal pronucleus, which carries no H3K27me3 marks (Zenk et al., 2017). However, in contrast to prior reports, the H3K27me3 signal becomes undetectable in later nuclear divisions (Figure 4A, NC8, NC13) recovering intensity by mid NC14 (Figure 4A, NC14). We also measured H2Aub in fixed embryos. Unlike H3K27me3, H2Aub is not strongly detected in nuclei immediately after fertilization (Figure 4B, NC3). We do detect, however, a low-level H2Aub signal on the condensed meiotic polar body that is barely above the cytoplasmic background signal (Figure 4B, NC10 asterisk) indicating that a small quantity of H2Aub is deposited on maternal chromatin. Even on compacted mitotic chromatin, H2Aub signal is undetectable prior to syncytial blastoderm stages (Figure 4B, NC3 & NC8). Coincident with the observed nuclear localization of PRE-binding factors just prior to ZGA, we begin to observe above-background H2Aub accumulation by NC12, and this signal increases throughout NC14 (Figure 4B, NC12 & NC14). These results indicate that whatever degree of H2Aub is inherited maternally, it is not strongly maintained during the cleavage divisions. We conclude that the H2Aub modification is established wholly *de novo* shortly before ZGA. Taken together, these observations are consistent with a model where the initial, maternally determined H3K27 state is diluted by the rapid syncytial divisions, and re-establishment of this modification state is restricted to late cleavage divisions by limiting nuclear localization of nucleating factors such as Pho, Cg and GAF.

**Figure 4:**
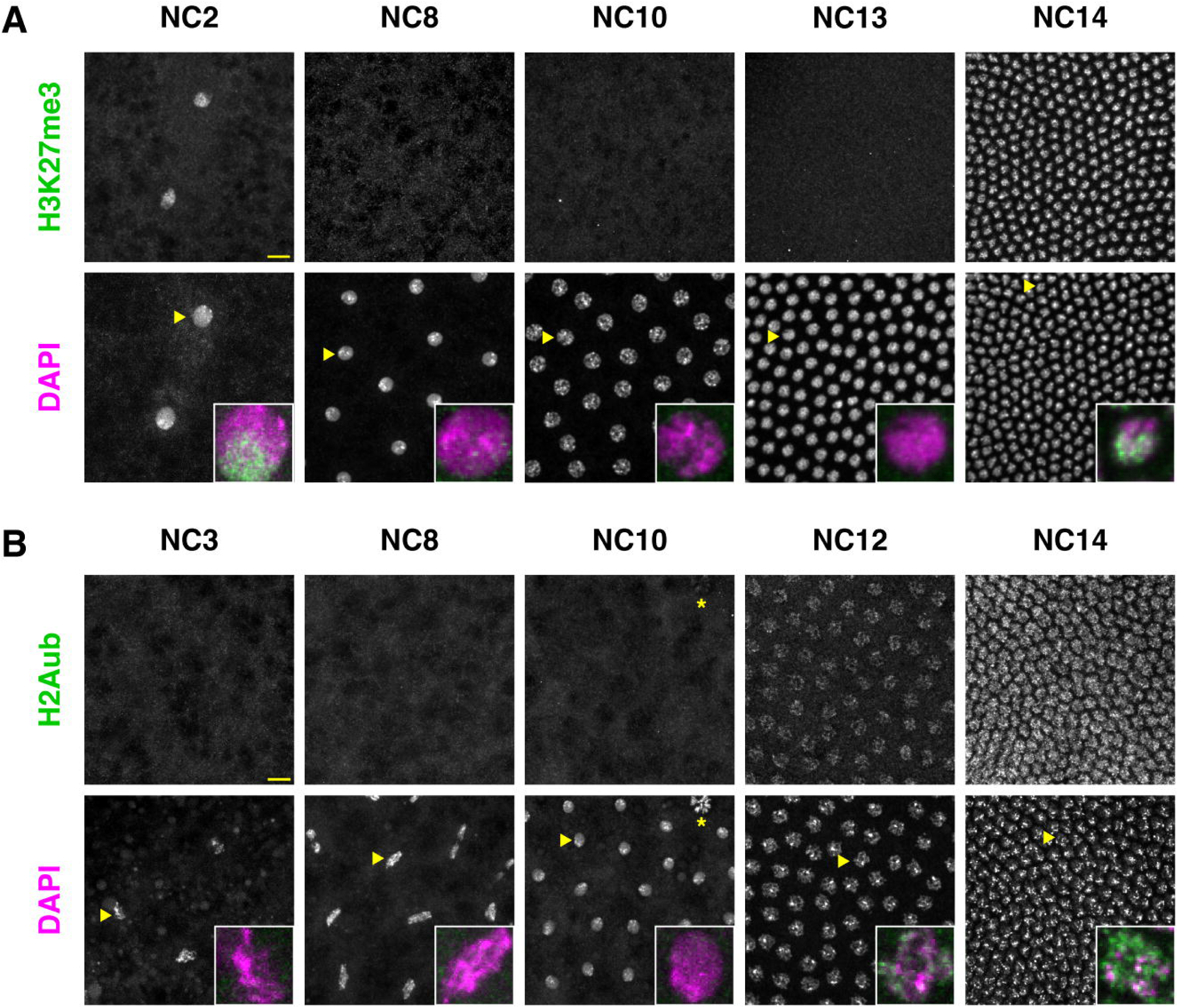
Immunostaining of H3K27me3 and H2Aub during cleavage stages. Embryos (*w^1118^*) were fixed and subjected to immunostaining for H3K27me3 (**A**) and H2Aub (**B**). Samples were co-stained with DAPI to highlight DNA/nuclei. Each panel shows maximum z-projected immunostaining for the indicated histone modification in the top row of images at the indicated nuclear cycle (top). The bottom row shows the corresponding DAPI image. Image intensities are normalized as described for Figure 3-figure supplement 1 to allow for semi-quantitative comparison of signal intensities across nuclear cycles. The inset shows a single enlarged nucleus (indicated by a yellow arrowhead) with a color merge of the modification (green) and DAPI image (magenta). The yellow asterisk in panel B, NC10 indicates the position of the polar body. H2Aub is not detectable in somatic nuclei above background until late cleavage divisions. The scale bar is shown in yellow and corresponds to 10 µm in all cases.

### Loss of GAGA-factor has limited impact on H3K27me3 establishment at ZGA

Based on the above observations, we next wanted to evaluate the impact of interfering with nucleating factors on the establishment of H3K27me3 patterns at ZGA. We attempted to perform maternal knockdowns of EGFP-Cg and Pho-sfGFP but consistent with prior reports, loss of function for either factor during oogenesis fails to yield significant quantities of viable embryos that develop to cellular blastoderm stages, thus limiting our ability to test this hypothesis with either factor (Beatty & Waddington, 1949; Breen & Duncan, 1986). Alternatively, we tested whether loss of GAGA-factor (GAF) would impact the re-establishment of H3K27me3 at ZGA. GAF, encoded by the *Trithorax-like* gene, has previously been implicated as a binding factor at PREs (Busturia et al., 2001; Mishra et al., 2001; Americo et al., 2002; Bejarano & Busturia, 2004). GAF is amongst a cohort of transcription factors including CLAMP and Pipsqueak that bind GA-rich (*(GA)_n_*) motifs that are widespread in the *Drosophila* genome and are frequently found within PRE sequences (Lehmann et al., 1998; Soruco et al., 2013). Mutation of *(GA)_n_* motifs within PRE reporter constructs demonstrates that such motifs can be necessary for PRE-dependent reporter silencing (Hagstrom et al., 1997; Horard et al., 2000; Busturia et al., 2001; Mishra et al., 2001; Americo et al., 2002; Brown et al., 2003; Bejarano & Busturia, 2004), and by ChIP-seq GAF is observed to bind 60% of pPREs (Tolhuis et al., 2006). Given prior reports of GAF’s role for pioneering chromatin accessibility at ZGA and its expected role for PRE-dependent gene silencing (Gaskill et al., 2021), we wished to test whether GAF function is required for the initial establishment at ZGA of H3K27me3 within PcG domains.

To knock down maternal GAF in early embryos, we turned to the recently developed Jabba-trap system (JT, (Seller et al., 2019)) to sequester an N-terminal EGFP-tagged GAF to cytoplasmic lipid vesicles (see Materials and Methods). Expression of JT using maternal alpha-tubulin 67c regulatory elements sequesters EGFP-GAF away from nuclei in cleavage stage embryos both during mitosis and interphase (Figure 5A, Figure 5-video 1 and Figure 5-video 2). By ChIP-seq, JT-mediated knockdown of EGFP-GAF (JT-GAF) results in a nearly complete depletion of GAF from blastoderm stage chromatin (Figure 5B), correlating well with the degree of cytoplasmic sequestration observed by microscopy. Given the extensive depletion in JT-GAF embryos, we next measured H3K27me3 to evaluate the dependency of this modification on GAF function.

**Figure 5:**
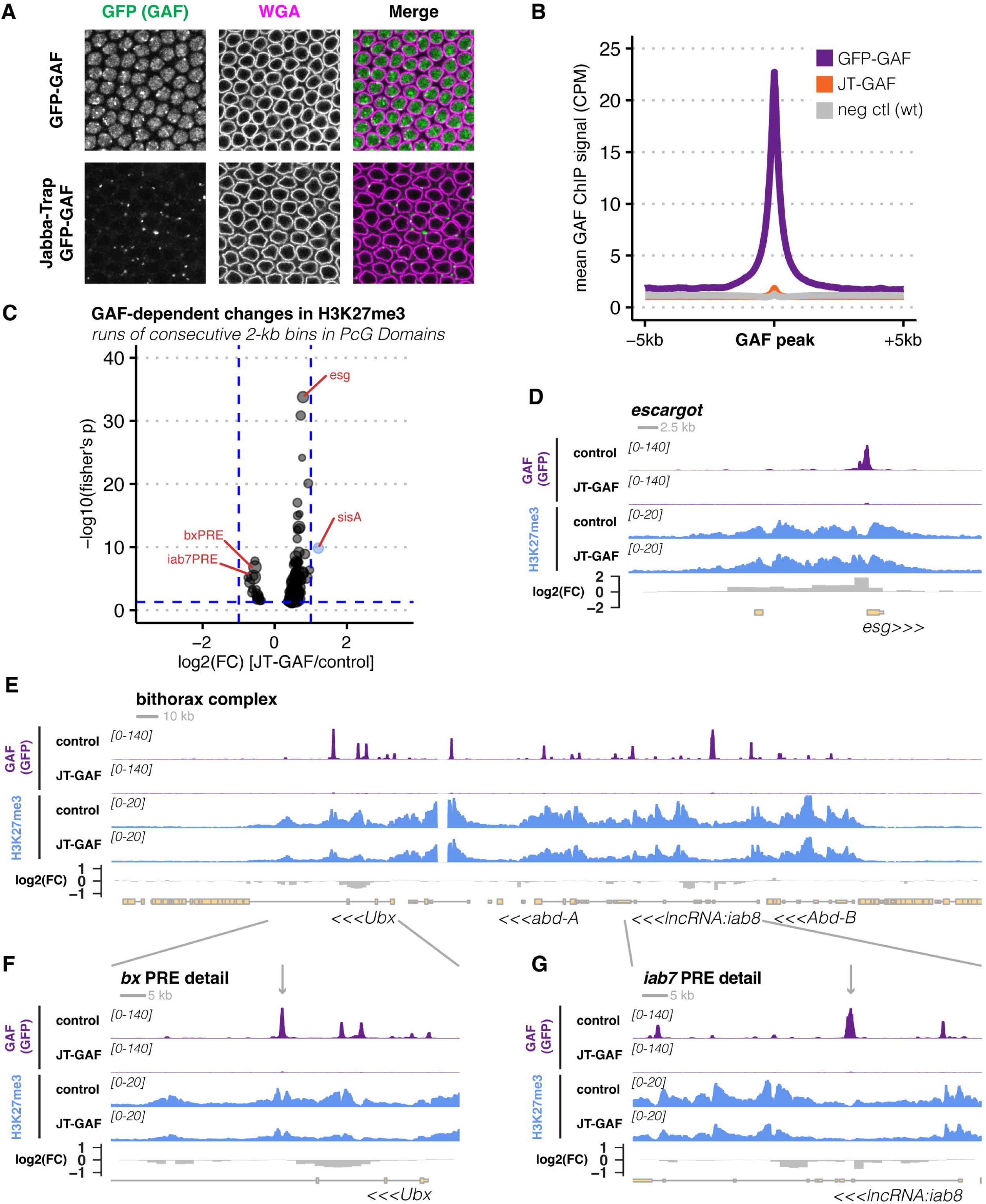
GAF knockdown has limited impact on accumulation of H3K27me3 at NC14. **A**) Control (EGFP-GAF) or GAF-knockdown (Jabba-Trap/+; EGFP-GAF) embryos were formaldehyde fixed and immunostained for GFP to detect GAF at NC14. Embryos were counterstained with wheat germ agglutinin (WGA) to highlight nuclear membranes. A single z-slice of individual channels (grey) and merged channels (GAF: green, WGA: magenta) shows that in the presence of Jabba-trap, EGFP-GAF is undetectable within NC14 nuclei. **B**) ChIP-seq was performed against EGFP-GAF in either positive control (EGFP-GAF), Jabba-trap GAF (JT-GAF), or negative control (*w^1118^*) samples with an anti-GFP antibody. Following peak calling, average GAF ChIP-seq signal in counts per million (CPM) over GAF peaks was measured and plotted, with ± 5 kb flanking sequence. The plot shows that whereas GAF (purple) is strongly enriched over peaks, Jabba-trap (orange) reduces GAF occupancy to near-background (grey) levels. **C**) A volcano plot is shown for the results of a DESeq2 differential enrichment analysis for H3K27me3 ChIP-seq between control (EGFP-GAF) and JT-GAF. Enrichment was measured over 2 kb bins and ‘runs’ of consecutive bins with p-values < 0.05 were merged. The volcano plot shows on the x-axis the average log2 fold change (FC) of the merged bins and the-log10 value of adjusted p-values merged using Fisher’s method. The size of a plotted point indicates the relative mean counts for the run of bins. Blue dotted lines indicate traditional cutoffs for significance testing (abs(log2(FC)) > 1 & p-adj < 0.05). Runs of bins exceeding these thresholds are colored blue. Regions of interest are labeled in red. While few regions show large-magnitude, GAF-dependent changes in K27me3, the general trend is for this modification to increase within PcG domains in the absence of GAF. **D**) Detail of ChIP-seq coverage over the *escargot* PcG domain. GAF ChIP seq for control or JT-GAF is shown at top (purple) and H3K27me3 for both genotypes is shown at bottom. The log2(FC) from the DESeq analysis over 2 kb bins is plotted in the bottom track. Scale bar is shown at top left, and y-axis range for coverage plots are indicated in brackets, with units of counts per million. **E-G**) Detail of ChIP-seq coverage over the bithorax complex (E) and zoomed-in views of the *bx* (F) and *iab7* (G) PREs are shown. Tracks are as described for panel D. The location of the PREs in F and G are indicated with a grey arrow. Scale bars are shown at top left for each respective plot. Y-axis range for coverage plots are indicated in brackets with units of counts per million.

Despite widespread binding of GAF to pPRE sequences and a prior expectation of GAF’s necessity for PRE function, we observe extensive establishment of H3K27me3 following JT-mediated knockdown of GAF and little major change in the modification status of pPREs (Figure 5C-G). A general challenge for quantitative evaluation of the impact of factor knockdown experiments on the establishment of H3K27me3 is the discrepancy in length scales at which transcription factors and PcG modifications interact with genomic loci. To quantify the effect of JT-GAF on H3K27me3, we counted recovered H3K27me3 ChIP-seq reads in 2 kb bins spanning the genome for replicates (n = 3) of each genotype and tested for differential enrichment between wild type (EGFP-GAF) and knockdown (JT-GAF) conditions using DESeq2 ((Love et al., 2014) see Materials and Methods and Source Code 1). This allowed us to score whether any of the 2 kb bins within PcG domains (n = 3883) showed significant changes in H3K27me3 distribution following GAF knockdown. By this approach, we observe only a limited number of 2 kb bins within PcG domains with modest decreases in H3K27me3 signal following GAF knockdown, and none of these pass significance testing (adjusted p-value < 0.05) at a standard magnitude of effect cutoff (log2(Fold Change) <-1) (Figure 5-figure supplement 1). A greater number of 2 kb bins within PcG domains show increases in H3K27me3 signal following GAF knockdown, and eight (0.2%) of these bins are scored as significant increases by the aforementioned criteria (Figure 5-figure supplement 1). Closer visual examination of H3K27me3 coverage identifies regions within PcG domains that may have functionally relevant, albeit partial, changes in H3K27me3 following GAF knockdown, where the magnitude of effect may not reach typical thresholds for significance calling. To identify such regions, we de-emphasized log2(Fold Change) as a criterion for significance calling and identified consecutive genomic runs of 2 kb bins within PcG domains with statistically significant differences in H3K27me3 signal (i.e., adjusted p-value < 0.05 with any log2(Fold Change)), yielding 361 runs of bins ranging in size from 2 to 24 kb in size (Figure 5-figure supplement 1, Materials and Methods). By this approach, 48% of PcG domains (n = 124) contain one or more runs of 2 kb bins with modest localized changes in H3K27me3 following GAF knockdown. Two of these domains have mixed changes, containing runs of bins that both show modest increases and decreases in H3K27me3 signal, the remainder show either increases or decreases. Of these, only 17 PcG domains (6.7%) are associated with one or more runs (n = 23) that undergo net reductions of this modification.

The bithorax cluster is the major representative PcG domain to undergo reduction in H3K27me3 following GAF knockdown, accounting for four of the 23 runs with net reductions averaging a log2(Fold Change) of-0.59 within significant 2 kb bins. At the bithorax cluster, we observe GAF to be necessary to attain maximal modification of the regions flanking the *bithorax* and *iab-7* PREs (Figure 5E-G, (Qian et al., 1991; Hagstrom et al., 1997; Mishra et al., 2001)). These effects of GAF on K27me3 deposition are likely to be direct, as 82% (n = 19) of these runs overlap with one or more GAF ChIP-seq peaks, including all four runs within the bithorax cluster. Effects like this are rare, however, and the major trend in these data is consistent with a role for GAF to counteract deposition of H3K27me3, since loss of GAF results in net gains of H3K27me3 at subregions within 109 PcG domains compared with only 17 domains with reductions. These GAF-dependent increases in H3K27me3 signal are also likely to be direct, as 109 (100%) of the runs contain one or more GAF ChIP peaks. As an example of this effect, the strongest gain in K27me3 signal following GAF knockdown is observed at the *escargot* locus, where an individual 2 kb bin corresponding to a GAF peak and the *escargot* transcriptional start site exceeds a log2(Fold Change) > 1 increase in K27me3, and runs of 2 kb bins within its associated PcG domain see an average log2(FoldChange) increase of 0.78 (Figure 5D and Figure 5-figure supplement 1). Taken together, these observations suggest that loss of GAF in blastoderm stage embryos results in limited effects on the establishment of H3K27me3, and that most of these moderate effects are consistent with a model where GAF operates at ZGA to counteract acquisition of high levels of this modification.

### Zelda is necessary for licensing a subset of PcG domains to deposit H3K27me3

The zinc-finger transcription factor Zelda is a major activator of the zygotic genome, operating through several known regulatory mechanisms (Liang et al., 2008). As a pioneer factor, Zelda is necessary for establishing and maintaining chromatin accessibility at a cohort of genomic loci required for the expression of a subset of early zygotic genes (Schulz et al., 2015; Sun et al., 2015; Hannon et al., 2017; McDaniel et al., 2019). As a transcription factor, Zelda also regulates gene expression, functioning mainly to activate target gene transcription (Harrison et al., 2011; Kvon et al., 2012; Stampfel et al., 2015; Hamm et al., 2017). Studies on *zelda* mutant embryos have also demonstrated additional phenotypes, including effects on genomic compartmentalization and formation of transcription factor hubs, that could either stem from pioneering or trans-regulatory activities or could represent additional biochemical functions (Hug et al., 2017; Dufourt et al., 2018; Mir et al., 2018; Yamada et al., 2019). Given Zelda’s role for activating expression of a cohort of zygotic genes, including those associated with PcG domains, we wished to measure the effect of Zelda on the establishment of PcG domains. We reasoned that Zelda could impact the modification state of these domains in at least two opposing ways. As an activator of transcription, Zelda could be required to promote an active state and suppress establishment of PcG modifications at target genes. Alternatively, as a pioneer, Zelda could be required to delineate cis-regulatory elements required for the establishment of PcG domains and modification of target genes. In the first hypothesis, loss of *zelda* would be predicted to result in the gain of PcG chromatin modifications at the expense of modifications associated with transcriptional activity. In contrast, with the second hypothesis, we would expect to observe in *zelda* mutants a loss of PcG modifications, independent of the transcriptional status of a locus. We therefore sought to distinguish these competing mechanisms.

To distinguish these hypotheses, we performed ChIP-seq for a battery of targets associated with transcriptional activity (RNA Pol2 (CTD pSer5)) and repression (E(z), H3K27me3, and H2Aub) comparing wild type embryos with those produced by *zld^294^* germline clones staged precisely at mid-NC14 (see Materials and Methods). Over the set of PcG domains in *zelda* mutants, we observe both increases and decreases in PcG histone modifications. As above, we quantified *zelda-*dependent changes in H3K27me3 ChIP-seq signal over 2 kb genomic bins. Unlike for GAF, loss of Zelda yields several genomic tracts with large-magnitude changes in H3K27me3 (Figure 6). Of the 3746 bins located within PcG domains for this analysis, 6% (n = 226) show statistically significant reductions in H3K27me3 at a log2(FoldChange) threshold of-1 (Figure 6-figure supplement 1). A smaller number (n = 62, 1.7% of bins in PcG domains) show increases in H3K27me3 signal following loss of *zelda* (Figure 6-figure supplement 1). Taking consecutive runs of bins with statistically significant adjusted p-values (independent of log2(Fold Change)), we recover 546 runs within PcG domains ranging in size from 2 to 46 kb in size. Of these, 55.1% (n = 301) show moderate to large increases in H3K27me3 signal, with 1.6% (n = 9) exceeding an average log2(FoldChange) threshold of 1 indicative of a large magnitude of effect. The remaining 44.9% (n = 245) of runs show decreases in H3K27me3 signal, with 5.3% (n = 29) exceeding an average log2(Fold Change) threshold of-1 indicating a large magnitude of effect (Figure 6A). We observe a high degree of correlation between Zelda-dependent changes in H3K27me3 and H2Aub (spearman’s rho = 0.78, Figure 6-figure supplement 1). The role of Zelda within PcG domains is therefore complex, with Zelda activity counteracting establishment of PcG modifications at a small number of domains, and Zelda activity being required for establishment at others.

**Figure 6:**
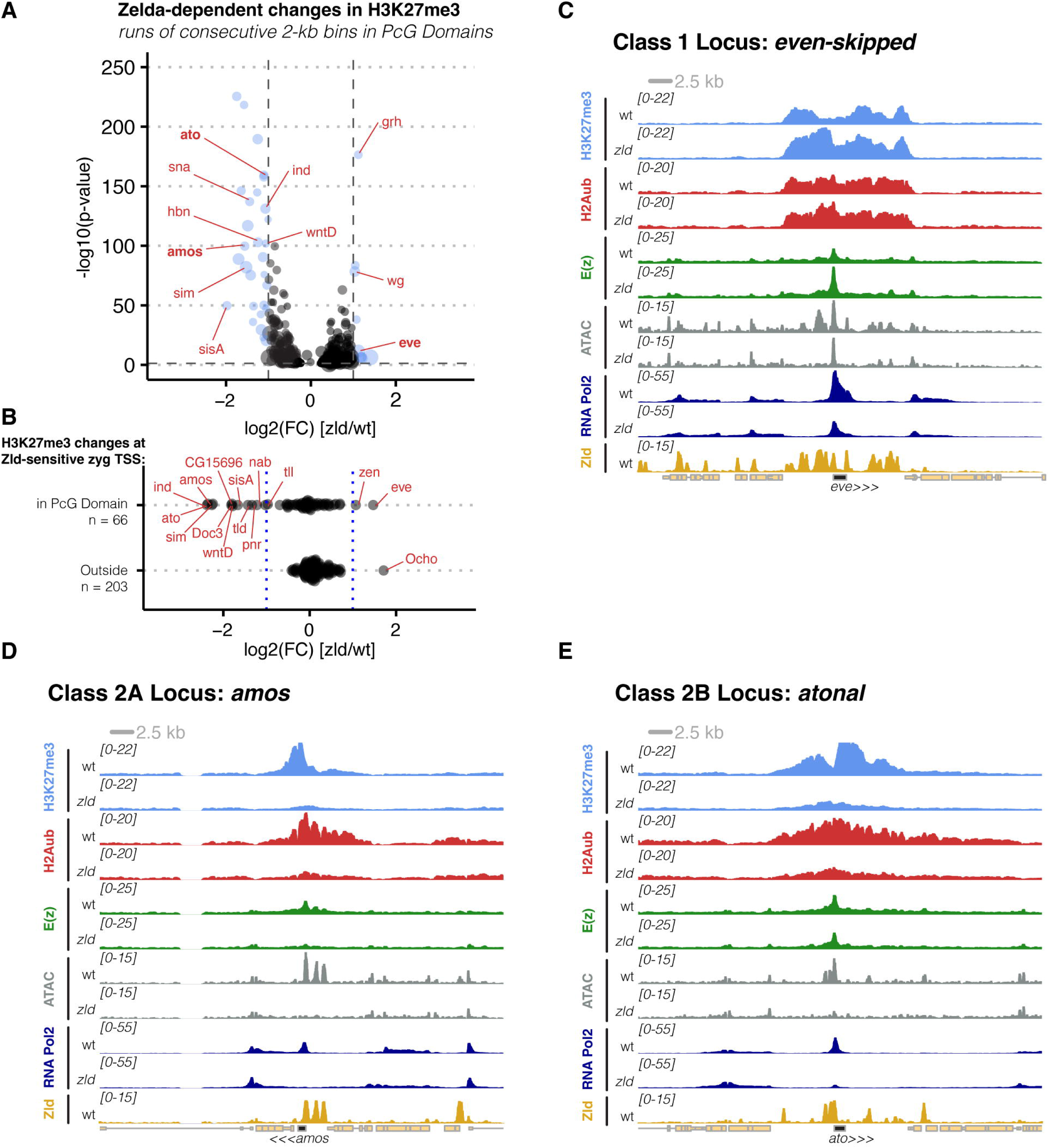
Zelda is necessary during ZGA for the establishment of H3K27me3 at a subset of PcG domains. **A**) A volcano plot is shown for DESeq2 analysis of H3K27me3 ChIP-seq comparing wild type and *zelda* germline clone embryos. Plotting methodology is as described in Figure 5C. On average, Zelda activity is required for establishment rather than antagonism of H3K27me3 at a subset of PcG domains. **B**) H3K27me3 is predominantly lost at sites requiring Zelda for RNA Pol2 recruitment. The log2(FC) of H3K27me3 ChIP-seq is plotted for 2 kb bins containing TSS that depend on Zelda for Pol2 recruitment. In the absence of Zelda, the TSS included in this analysis have major reductions in Pol2 occupancy. TSS were grouped into two categories: those within a PcG domain (top) and those outside a PcG domain (bottom). In the absence of Zelda, TSS within PcG domains that lose Pol2 (n = 66) also typically see losses in H3K27me3, with the exception of a few loci (*eve, zen*) that see increases. Loss of Pol2 at Zelda-sensitive TSS outside of PcG domains (n = 203) does not correspond to *de novo* accumulation of H3K27me3, except at the *ocho* locus. **C-E**) Coverage plots for loci representing classes of PcG domain responses to *zelda* loss of function. ChIP for factors indicated at left were performed for wild type (top track) and *zelda* germline clone (bottom track) embryos. ATAC data are from (Hannon et al., 2017) and Zelda ChIP-seq is from (Harrison et al., 2011). Scale bar is at top left and the y-axis range for each displayed track is indicated in brackets, with units of counts per million. Panel C shows a class 1 locus, *even skipped*. Class 1 are PcG domains where increases in K27me3 are seen in the absence of *zelda*. Panel D and E show Class 2 loci, which are PcG domains where decreases in K27me3 are seen in the absence of *zelda*. Class 2A loci (panel D) see reductions in K27me3 accompanied by loss of E(z) recruitment. Class 2B loci (Panel E) see reductions in K27me3 without significant loss of E(z) recruitment.

We next compared how changes in H3K27me3 status correlated with changes in RNA Pol2 recruitment following loss of Zelda. Depending on biological context, when early zygotic loci fail to be activated in *zelda* mutant embryos, a subset of these sites undergo changes in H3K27me3 status. As previously reported, we observe a significant loss of RNA Pol2 occupancy in *zld* mutant embryos at a subset of early zygotic target genes (Blythe & Wieschaus, 2015), and similar losses in histone modifications associated with active transcriptional states are also observed (Li et al., 2014). Through DESeq2 analysis, of 798 early, zygotic-only transcribed mRNA-encoding loci (Chen et al., 2013), 269 (33%) show statistically significant reductions of RNA Pol2 occupancy in *zelda* mutant embryos. Within this set of zygotic-only genes, 155 (19%) reside within a PcG domain, and 43% (n = 66) of these require Zelda for full RNA Pol2 occupancy. Comparing correlations between Zelda-dependent losses in RNA Pol2 occupancy and changes in H3K27me3 deposition, while there are 66 early zygotic-only genes within PcG domains that require Zelda for RNA Pol2 occupancy, only 14 (21%) of these show significant changes in H3K27me3, and of these, two account for increases, and 12 account for decreases in H3K27me3 deposition (Figure 6B). Therefore, while Zelda affects H3K27me3 establishment at a subset of targets within PcG domains, not all sites requiring Zelda for RNA Pol2 recruitment show changes in this modification. Of the subset of targets with Zelda-dependent changes in both H3K27me3 and RNA Pol2, most sites demonstrate losses in occupancy of both factors following loss of Zelda.

We also tested for evidence that Zelda could be required to counteract establishment of PcG modifications at genes not associated in wild type embryos with a PcG domain. As described above, E(z) binds near gene promoters located both within and outside of PcG domains. All 66 of the genes within PcG Domains that require Zelda for full RNA Pol2 binding also coincide with a promoter-proximal E(z) peak. The majority (134/ 203, 66%) of zygotic-only genes outside PcG domains that require Zelda for RNA Pol2 occupancy also coincide with a promoter-proximal E(z) peak. Despite this, loss of Zelda does not result in significant acquisition of H3K27me3 at genes located outside of PcG domains. Of the 203 genes requiring Zelda for RNA Pol2 recruitment, only one, *Ocho*, shows a significant increase of H3K27me3 over its transcription start site upon loss of Zelda (Figure 6B). Therefore, these Zelda-dependent effects on H3K27me3 deposition are almost entirely limited to the territories of PcG domains, and Zelda does not counteract in early embryos a latent competency for additional genomic loci to acquire PcG modifications. These results indicate that Zelda influences the establishment of chromatin modifications within a subset of PcG domains and allows us to categorize *zelda*-sensitive domains into two major classes. Overall, 16% (n = 38) of PcG domains are *zelda*-sensitive. Class 1 domains (n = 9) demonstrate increases and Class 2 domains (n = 29) demonstrate losses in both K27me3 and H2Aub following loss of Zelda. We focus on the characterization of these two classes of targets in the following analysis.

The increases in PcG modifications within Class 1 domains support a mechanism where Zelda’s trans-activation function is necessary to counteract the establishment of H3K27me3 and H2Aub. This class includes PcG domains associated with patterning genes such as *even-skipped*, *wingless* and *grainyhead*. We focus here on *even-skipped* as an exemplar of this class (Figure 6C). In *zelda* mutants, the *eve* transcription unit has significantly reduced RNA Pol2 occupancy, and several cis-regulatory elements within the *eve* PcG domain lose chromatin accessibility, but accessibility is retained in a promoter-proximal region that harbors the major peak of E(z) binding within this domain. E(z) occupancy is increased at the *eve* promoter pPRE in *zelda* mutants, and corresponding increases in H3K27me3 and H2Aub are observed within this locus. Notably, these increases are limited to the bounds of the wild type *eve* PcG domain, indicating that boundary integrity at this locus remains intact in the absence of *zelda*. These observations at the *eve* locus, and for Class 1 domains in general, are consistent with a model where loss of a trans-activating factor can result in gains in PcG activity at least at sites that are competent to do so.

In contrast to the Class 1 domains, Zelda is necessary for establishment of PcG modifications within 28 Class 2 domains, where loss of Zelda results in a failure to establish H3K27me3 and H2Aub during NC14. Comparison of chromatin modification states between wild type and *zelda* mutant embryos within the set of Class 2 domains identifies two major sub-classes which we will refer to as 2A and 2B, both of which show extensive loss of PcG modifications, but stemming from different effects of Zelda on chromatin accessibility and E(z) recruitment. In *zelda* mutants, Class 2A domains fail to establish PcG modifications, and this effect correlates with Zelda-dependent losses in chromatin accessibility and E(z) binding. Overall, we identify 6 total Class 2A domains, which contain *sisterless A, amos, cadherin-N, wntD, Chronophage,* and *Anion exchanger 2*. We focus here on *amos* as the exemplar of this class (Figure 6D). In *zelda* mutants, RNA Pol2 occupancy of *amos* is completely eliminated. Similarly, the three major accessible chromatin regions within this domain are *zelda*-dependent and fail to open in *zelda* mutants. E(z) binding within this domain is correspondingly lost in *zelda* mutants, and H3K27me3 and H2Aub are not established within this domain. In contrast, Class 2B loci (n = 17) demonstrate similar failures in establishment of PcG modifications but retain tracts of accessible chromatin and E(z) binding. Class 2B domains include developmental regulators such as *single-minded, snail, zfh-1, atonal, teashirt,* and *ind*. We focus here on *atonal* as the exemplar of this class. Similar to the *amos* domain, the *atonal* locus fails to fully establish H3K27me3, and to a lesser extent, H2Aub. While several prominent accessible chromatin regions within the *atonal* domain are Zelda-dependent, the main E(z) binding site within the *atonal* promoter shows a nearly complete loss of accessibility, but no quantitative difference in overall E(z) binding. Yet, despite this retention of E(z), very little H3K27me3 is deposited at this locus in *zelda* mutants. These observations for Class 2 domains are consistent with a mechanism where Zelda activity is necessary for designating a subset of genomic regions as PcG domains, albeit through different mechanisms. For Class 2A loci, Zelda operates as would be expected for a pioneer factor, namely, to establish accessibility at critical cis-regulatory elements, including presumptive PRE elements. Zelda’s role at Class 2B loci is less clear: although E(z), and presumably PRC2, is retained at pPRE sequences, in the absence of Zelda, these sites are not fully licensed to modify the surrounding domain.

## Discussion

In this study, we sought to understand the mechanism for re-establishment of the H3K27me3 modification during *Drosophila* ZGA. Prior studies have been inconsistent on whether this modification persists throughout the cleavage divisions, or whether it is largely established following large-scale ZGA (Chen et al., 2013; Li et al., 2014; Zenk et al., 2017; Reinig et al., 2020; Cardamone et al., 2025). In addition to repeating prior observations of limited H3K27me3 on late cleavage-stage chromatin and rapid accumulation during NC14, we extend these prior observations by showing that the DNA-binding factors thought to be necessary for nucleation and maintenance of PcG states demonstrate limited nuclear localization until the final few cleavage divisions. These observations are consistent with a model where the rapid cleavage divisions operate to reprogram the zygotic genome by diluting parental histone post-translational modifications which could otherwise influence gene expression states. Histone modifications associated with both active and inactive states are established upon cell cycle lengthening at NC14 (Chen et al., 2013; Li et al., 2014), and it has been argued that, in the absence of gene activation, sites will default to a repressed, PcG-modified state (Li et al., 2014; Ghotbi et al., 2021). Our results argue that establishment of the active-versus-repressed dichotomy requires prior regulatory input from transcription factors such as Zelda.

Here we observe that, at ZGA, E(z) and the nucleating factors Pho, GAF, and Cg bind broadly across the genome, but H3K27me3 deposition is only observed at a small subset of these sites. Prior genome-wide profiling of PcG components has observed a range of concordances between PcG factor localization and domains of H3K27me3 deposition. Early genome-wide studies in cell lines and broadly staged embryos found high degrees of concordance between PcG factors and H3K27me3 deposition (Schwartz et al., 2006; Tolhuis et al., 2006; Schuettengruber et al., 2009). However, more contemporary studies on tissues and different cell lines have found extensive binding of PcG components including nucleating factors outside of H3K27me3 domains (Schaaf et al., 2013; Loubière et al., 2016; Kang et al., 2017; Pherson et al., 2017; Brown et al., 2018), raising the possibility that PcG domains vary as a function of cell fate determination and that PcG binding outside of domains reflect a form of bivalency or poising for potential PcG regulation. Our results indicate that drastically reducing the fraction of active genes at ZGA does not lead to a corresponding increase in deposition of H3K27me3 outside of PcG domains and argues against a simple model where inactive genes default to a H3K27me3 modified state. Rather, this default state is specified. Zelda plays multiple roles within PcG domains to influence the histone modification status: depending on biological context, Zelda is necessary to counteract H3K27me3, pioneer accessibility of key pPRE sites, or license bound PRC1/2 to deposit their respective histone modifications. At a subset of targets, loss of Zelda activity reduces transcriptional activation, and results in increased H3K27me3, reflecting the expected dichotomy between H3K27me3 and gene on and off states (Bowman et al., 2014; Papp & Müller, 2006). At these sites, Zelda could function to counteract H3K27me3 modification status at certain sites by promoting the acetylation of H3K27 through recruitment of the CBP co-activator (Galouzis et al., 2025; Marsh et al., 2025). Zelda’s role as a pioneer factor is well appreciated, and PRE function is thought to require establishment of accessible chromatin states (Mito et al., 2007; Schulz et al., 2015; Sun et al., 2015). At present, the mechanism whereby Zelda could function to license PRC1/2 to deposit histone modifications is unclear, but these results point to a need to understand further the mechanisms whereby genomic loci are specified to partition into distinct classes of epigenetic regulatory compartments. While enrichment of E(z) at sites that require Zelda for licensing does not appear to depend on establishment of accessible chromatin, perhaps accessibility is required for additional cofactors to bind and signal to PRC1/2 to deposit modifications. As such, we emphasize that Zelda-dependent licensing could stem either directly or indirectly from Zelda function. Zelda has also been implicated in the formation of higher order chromatin structure (Hug et al., 2017). While loci like *eve* retain compartmentalization of PcG modified states in the absence of Zelda function, perhaps loci requiring Zelda for licensing fail to attain the correct higher-order chromatin structure necessary for efficient establishment of PcG states (Loubière et al., 2016; Ogiyama et al., 2018). Notably, Zelda’s effect on PcG states is limited to a subset of Zelda targets within PcG domains. Zelda-dependent PcG domains are enriched for loci involved in neurogenesis (e.g., *sna, amos, ind, ato, hbn, sim, N-cad*), where Zelda is also required in later developmental stages (Staudt et al., 2006; Larson et al., 2021). Within the context of embryonic patterning, Zelda could operate at these sites under the context of reprogramming not only to “put them in to play” but also to prime these sites for later silencing in non-neural cell lineages. Additionally, not all Zelda-bound pPREs depend on Zelda for licensing and not all pPREs contain Zelda binding sites. We speculate that additional factors (or combinations of factors) will function like Zelda to license pPREs to establish the active-versus-repressed regulatory dichotomy at developmental patterning genes.

We sought to test directly the requirement for GAF in the establishment of PcG modifications at ZGA and observe that loss of GAF function has limited effect on the initial distribution of H3K27me3. GAF has been associated with PRE function due to the frequent occurrence of *(GA)_n_*sites in PREs, and observations of impaired silencing function of PRE reporters when such sites are mutated (Busturia et al., 2001; Mishra et al., 2001; Americo et al., 2002; Bejarano & Busturia, 2004). Additionally, GAF has been shown to be required to establish long-range interchromosomal contacts between homologous copies of endogenous and transgenic PREs (Fitz-James et al., 2025), whose pairing is necessary for silencing function. Given the strong loss of silencing observed at PRE reporters with mutated *(GA)_n_* motifs, we speculate that loss of maternal GAF function is compensated for by other *(GA)_n_*binding factors such as CLAMP and Pipsqueak. Alternatively, it is possible that GAF is required during the maintenance but not the establishment phases of PcG domains, however prior work has shown no requirement for GAF to silence Hox genes in imaginal disc tissues (Brown et al., 2003). Loss of function studies for other PRE-binding factors in later developmental contexts have shown that loss of Pho can strongly reduce not only H3K27me3 maintenance but also binding of PRC1/2 components (Brown et al., 2018). To address these questions in the context of establishment, robust methods for depleting maternally supplied proteins in the embryo will be required. Future studies will address this issue.

Maintenance of long-term transcriptional repression in proliferative cells by the PcG system depends on PRE function to nucleate the replacement of modified histones following DNA replication (Coleman & Struhl, 2017; Laprell et al., 2017). Here, we repeat prior measurements demonstrating that the H3K27me3 modification fails to persist on chromatin through the cleavage divisions, and that key nucleating factors demonstrate limited nuclear localization until shortly before ZGA. Prior kinetic modeling of E(z) catalysis on cleavage stage chromatin supports this conclusion (Lundkvist et al., 2023; Degen et al., 2025). *Drosophila* and other organisms (Ciabrelli et al., 2017; Zenk et al., 2017; Tabuchi et al., 2018; Mei et al., 2021; Fitz-James et al., 2025) demonstrate the capacity to maintain PcG-dependent epigenetic states across generations, in a phenomenon termed transgenerational epigenetic inheritance (TEI). In the case of *Drosophila*, TEI has been most clearly shown through the stable inheritance over dozens of generations of PRE-bearing transgenes entrained either to active or repressed states (Ciabrelli et al., 2017). In mice and zebrafish, the H2Aub modification state is propagated through the maternal germline and serves as the mechanism for re-establishment of patterns of PcG modifications in early development through allosteric stimulation of PRC2 activity (Mei et al., 2021; Hickey et al., 2022; Matsuwaka et al., 2025). PRC1 activity is limited in the *Drosophila* ovary (Iovino et al., 2013), and consistent with this, we do not observe significant H2Aub on cleavage stage chromatin until shortly before ZGA, thus limiting its potential role in the context of *Drosophila* TEI. If mechanisms for TEI follow the same rules as maintenance of PcG states in late embryos, larvae, and adult tissues, then the observed lack of key nucleating factors until late cleavage divisions should limit the extent to which maternal H3K27me3 states are directly maintained (Coleman & Struhl, 2017; Laprell et al., 2017). Further investigation is necessary to determine the mechanism whereby parental PcG states can be inherited across the maternal to zygotic transition.

### Limitations of the Study

We acknowledge the following limitations to this study. Given the nature of antisera and immunodetection, we cannot completely rule out the action of low level (undetectable) but biologically significant quantities of Pho, GAF, Cg, or even H3K27me3 operating at early cleavage divisions. However, whatever level of these factors may be present in early cleavages, their nuclear abundances increase many-fold by NC14. The strength of this quantitative conclusion is driven by our use of GFP chimeras for the measurements of PcG effector proteins, which allows us to use a single antibody source and staining/imaging conditions, thus limiting the expected variability between antisera raised against distinct targets. We also acknowledge that while GAF knockdown by Jabba-trap eliminates nearly all detectable GAF on chromatin, it remains possible, however unlikely, that the very little GAF that remains associated with PREs is sufficient to promote H3K27me3 establishment at ZGA and can account for the reported limited effect of GAF knockdown on this modification. Finally, we acknowledge that further loss-of-function analysis will be necessary to determine the key nucleating factors responsible for the initial establishment of the zygotic H2Aub and H3K27me3 landscape.

## Data Availability

ChIP-seq datasets generated for this study have been deposited in Gene Expression Omnibus (accession number: GSE299311). Files for reproducing the differential enrichment analysis and code can be accessed at https://github.com/sblythe/Gonzaga_Saavedra_2025_Code_Supplement. The source code for the differential enrichment analysis is available as Source Code 1.

## Additional Datasets Used

The Zelda ChIP-seq track and peaks list are from the 3-hour timepoint from (Harrison et al., 2011) (GSE30757).

ATAC-seq tracks comparing NC14 wild type and *zelda* germline clone chromatin accessibility are from (Hannon et al., 2017) (GSE86966)

## Materials and Methods

### Drosophila Stocks and Gene Editing

Drosophila stocks were maintained on an enriched high-agar cornmeal media as reported previously (Cline, 1978). Embryos were collected from small population cages on apple-juice agarose plates smeared with a small amount of active yeast paste.

The wild type genotype for this study is *w[1118]* and serves as the ultimate genetic background for gene edited stocks. CRISPR/Cas9 was used to generate fluorescent protein chimeras of E(z), Pho, and Combgap. The Cas9 transgenic stock used for genome editing for *E(z)* and *pho* was *y w; nos>Cas9/CyO*. *cg* genome editing was performed in the *y vas>Cas9 ZH2A w* background. Injections were performed in-house as described previously (Soluri et al., 2020).

### Combgap

Of the four GFP chimeras used in this study, *cg* has the most complex splice-variant structure and current gene models identify no constant N-or C-terminal coding sequence. The N-terminus, however, only has two predicted splice isoforms, both originating from a common transcription start site, with one minor splice variant (isoform C) omitting the otherwise canonical translation start site. We decided to generate a *cg* allele that would produce *cg* mRNA variants with a common start codon by eliminating alternative start codon usage of isoform C variants. To do this, we mutated the splice acceptor that distinguishes the isoform C splice variant (chr2R:14174461-14174461 (G>A)) and we introduced a Drosophila Codon Optimized EGFP sequence plus an 11 amino acid linker via homology directed repair to replace the canonical start codon. Transcripts produced from this *EGFP-cg* locus would either contain the entire *Start-EGFP-linker-cg* sequence, or isoform C splice variants would now splice-in to the EGFP cassette at a downstream cryptic splice acceptor site upstream of EGFP glycine 4, and fail to initiate translation at the intended methionine 1. We therefore subsequently replaced glycine 4 with a methionine residue to allow for isoform C variants to initiate translation at the N-terminus of the EGFP cassette. We sequenced *EGFP-cg* cDNAs via Sanger sequencing to verify that all spliced isoforms contained an N-terminal EGFP cassette with a functional start codon. Genome-edited *EGFP-cg* flies are homozygous-viable.

*cg* gRNA target: chr2R:14174457-14174476 (-) 5’-GAAGGGCGGCGGATTCTGGG (CGG)-3

The underlined bases in the *cg* gRNA target indicate sites where synonymous mutations were introduced to eliminate cutting of the homology directed repair template (G>A in the PAM sequence), and where the splice acceptor for isoform C was mutated as described above.

The homology repair template contained 500 bp of left-and right-homology regions upstream of the EGFP-linker cassette and downstream of the mutation to the isoform C splice acceptor site. The homology repair template was constructed in a modified pBluescript vector with a 3xP3>dsRed cassette in the vector backbone that could be used to screen out unwanted whole-plasmid integrants produced by homologous recombination. The gRNA was expressed using the pU6 BbsI vector. Likely transformants were identified by measuring the frequency of mutations induced by *ebony* co-CRISPR as described previously (Soluri et al., 2020). Individual edited chromosomes were identified using PCR and correct site integration was verified by Sanger sequencing.

### E(z)

E(z) is expressed with a single common start codon. Prior work (Steffen et al., 2013) determined that E(z) could tolerate an N-terminal EGFP fusion. We therefore targeted the N-terminus with a multifunctional cassette, *start-EGFP-AID-3xHA-linker*, that would facilitate visualization, conditional knockdown, and immunodetection/affinity purification through the EGFP, auxin inducible degron (AID), and 3xHA epitope tag components. Genome-edited *EGFP-E(z)* flies are homozygous-viable. In experiments not presented in the current manuscript, we verified that EGFP-E(z) could be targeted by Jabba-Trap to yield maternal-zygotic E(z) loss of function phenotypes at high frequency and penetrance. Additionally, we determined that the auxin inducible degron tag inserted at this position was not sufficient to yield knockdown of E(z).

*E(z)* gRNA target: chr3L:10637753-10637772 (+) 5’-GCTATTCATAATGCCTTCGA (GGG)-3’

The underlined bases indicate the position of the *E(z)* start codon on the minus strand. Because the gRNA straddles the insertion site at the start codon, no additional mutations were introduced to prevent targeting of the homology directed repair template.

The homology repair template contained 500 bp of left-and right-homology regions upstream and downstream of the *start-EGFP-AID-3xHA-linker* cassette. The homology repair template was constructed in a modified pBluescript vector with a 3xP3>dsRed cassette in the vector backbone that could be used to screen out unwanted whole plasmid integrants produced by homologous recombination. The gRNA was expressed using the pU6 BbsI vector. Likely transformants were identified by measuring the frequency of mutations induced by *ebony* co-CRISPR as described previously (Soluri et al., 2020). Individual edited chromosomes were identified using PCR and correct site integration was verified by Sanger sequencing.

### Pho

Because a prior report indicated a lack of complementation for N-terminal GFP-Pho chimeras (Steffen et al., 2013), we chose to target the single Pho C-terminus with a *linker-3xHA-AID-sfGFP-stop* cassette. Genome-edited *pho-sfGFP* are homozygous-viable. In experiments not presented in the current manuscript, Jabba-Trap knockdown of *pho-sfGFP* yields reduced fecundity of adult females and the majority of eggs fail to develop to the cellular blastoderm stages. The efficacy of Pho knockdown via the AID tag has not been evaluated.

*Pho* gRNA target: chr4:1172891-1172910 (+) 5’-CATATACCACAAACGGTACA (TGG)-3’

The gRNA cuts 22 bp upstream of the annotated *pho* stop codon.

The homology repair template contained 273 bp of left-and 282 bp of right-homology sequence flanking the gRNA cut site, and a silent G>A mutation at position 3 of the PAM sequence to prevent cutting of the repair construct. The homology repair template was constructed in the pJAT7 vector (Stern et al., 2024) which also drove expression of the gRNA. pJAT7 inserts were synthesized by Twist Biosciences and introduced with an LR Clonase reaction as described previously (Stern et al., 2024). To facilitate selection of positive insertions, the *sfGFP* sequence was split at a TTAA sequence and a PiggyBac cassette containing a 3xP3>dsRed cassette was inserted, allowing for scoring of genome-edited individuals by red fluorescence in the eye. Whole plasmid integrants typically expressed substantially higher levels of 3xP3>dsRed fluorescence. Individual dsRed+ transformants without whole plasmid integration were verified using TagMap (Stern, 2017) followed by Nanopore sequencing (Plasmidsaurus, Kentucky, USA). Individuals homozygous for the C-terminal PiggyBac 3xP3>dsRed cassette died as pharate adults. The PiggyBac-dsRed cassette was excised by crossing single adult dsRed+ males to *alpha-tubulin>PB-transposase; Gat[eya]/Ci[D]* (product of crossing Bloomington Drosophila Stock Center stocks 32070 and 90852) and then by crossing the resulting dysgenic male progeny to *yw; Gat[eya]/Ci[D]* virgins. F2 males from the dysgenic cross were screened for loss of the PiggyBac cassette and individual stocks were verified by Nanopore sequencing (Plasmidsaurus) of PCR products spanning the sfGFP insertion site.

### GAF

An *EGFP-3xHA-linker* sequence was added to the N-terminus of the *Trl* locus (encoding GAF) by phi-C31 mediated insertion of a *attB-EGFP-3xHA-Trl_exon1-intron-LoxP-vector* sequence into a CRISPR modified stock, *attP-3xMyc-Trl_exon1-intron-LoxP*, kindly provided by Tsutomu Aoki and Paul Schedl (Princeton University). Insertion of the EGFP cassette into the target locus results in a duplication of the *Trl* N-terminus where the outer duplication contains the engineered EGFP cassette, and the inner duplication is the original N-terminus. The insertion provides a second LoxP site that allows for the subsequent excision of the original *Trl* N-terminus leaving only the *EGFP-3xHA-Trl* coding sequence with an upstream *attL* and a single residual intronic *LoxP* site. Excision of the duplicated *Trl* N-terminal sequence was performed by crossing transformants to *y w; MKRS/TM6B p{mini-white = Crew}DH2* (Bloomington Drosophila Stock Center 1501). Genome-edited *EGFP-Trl* individuals are homozygous-viable. Sequences were verified by Sanger sequencing of PCR products flanking the modified locus.

### Zelda Germline Clones

As described previously (Blythe & Wieschaus, 2015), *zelda* germline clones were made by crossing *w zld[294] FRT19A/FM7h; RpA-70-EGFP* females to *ovoD hs>flp FRT19A/Y* males and performing 3x 1-hour 37°C heat shocks on days 3-5 of development.

### Jabba-Trap

Based on the original reported design for Jabba-Trap (Seller et al., 2019), we designed a transgene that would yield maternal expression of Jabba-Trap directly driven by the maternal *alpha-Tubulin 67C* regulatory region. To do this, we synthesized the cDNA encoding the RG isoform of *jabba* chimerically fused to an anti-GFP nanobody (Kirchhofer et al., 2010) on both the N-and C-terminus, introducing 30 bp of codon optimization to the 5’ end of the N-terminal and 3’ end of the C-terminal anti-GFP nanobodies to allow for PCR amplification. Synthesized DNA was amplified by primers that appended NotI and NheI sites to the construct, and this product was cloned into a transgenic vector (pBABR) containing the 5’ regulatory sequences for *alpha-tubulin 67C* through the first intron and the *spaghetti squash* 3’ UTR. pBABR contains an attB site suitable for phi-C31 transgenesis. Transgenic lines were established at the attP40 and VK000033 attP landing sites by microinjection of the transgenic vector into phiC-31 expressing embryos.

The JT-GAF knockdown experiment was performed in embryos derived from the cross between *tub67C>JabbaTrap{attP40}/+; EGFP-Trl* mothers and *EGFP-Trl* fathers yielding embryos where both maternal and zygotic EGFP-GAF protein is sequestered by maternally expressed Jabba-Trap. Phenotypic details of this knockdown beyond effects on H3K27me3 will be reported elsewhere.

### Additional stocks used

- *y w; {mini-white = RpA-70-EGFP}attP2* (Blythe & Wieschaus, 2015)

- *w p{mini-white = His2Av-RFP}1* (Blythe & Wieschaus, 2015)

- *w zld[294] FRT19A/FM7h* (Liang et al., 2008)

- *w p{mini-white = His2Av-RFP}1 zld[294] FRT19A/FM7h* (recombinant of the two prior stocks)

- *C(1)DX y w f/ p{mini-white = ovoD1-18}P4.1, p{ry = hs>Flp}12, y w sn FRT19A* (Blythe & Wieschaus, 2015)

- *E(z)[61]/TM3, Sb* (Jones & Gelbart, 1990)

### Plasmids

Vectors for performing CRISPR mutagenesis are as follows:

- pBS-3xP3>RFP (Soluri et al., 2020). Cloning of HDR constructs for classic CRISPR approaches.

- pU6-2-BbsI-gRNA (gift from Kate O’Connor-Giles, Drosophila Genomics Resource Center). Cloning of gRNA constructs for classic CRISPR approaches.

- pJAT7 (gift from David Stern (Addgene plasmid # 204292)). Cloning of gRNA and HDR sequences for new CRISPR approaches.

### ChIP Sample Preparation

Timed collections of the indicated genotypes were performed, and embryos were dechorionated for 1-2 minutes in 4% sodium hypochlorite followed by extensive washing in deionized water. Dechorionated embryos were crosslinked by incubation for 15 minutes in a biphasic solution of 1x Phosphate Buffered Saline with 0.5% Triton X-100: Heptane:: 2 ml: 6 ml to which 180 µl of 20% paraformaldehyde was added. Crosslinked embryos were collected by centrifugation and the crosslinking reaction was stopped by a 1-minute incubation in 1x PBS + 0.5% Triton X-100 + 125 mM Glycine followed by three washes in 1x PBS + 0.5% Triton X-100. Crosslinked embryos were sorted by transferring the sample to a 35 mm petri dish lined with 1% agarose overlaid with a large volume of 1x PBS + 0.5% Triton X-100. For samples sorted by cell cycle (NC13, NC14E/M/L), the maternal genotype included RpA-70-EGFP which produces bright nuclear fluorescence scorable under an epifluorescent stereoscope at 100x magnification. NC14 embryos were subdivided into Early/Mid/Late groups by measuring the degree of nuclear elongation and cellularization as described previously (Blythe & Wieschaus, 2015). For measurement of E(z) binding comparing wild type and *zelda* germline clones, embryos were sorted using His2Av-RFP expression and the maternal genotypes for this experiment were: wild type = *w p{His2Av-RFP}1; EGFP-E(z)/+* and *zelda (clones) = w p{His2Av-RFP}1 zld[294] FRT19A; EGFP-E(z)/+.* To stage *zelda* mutant collections, we relied on the following criteria. Embryos were sorted first based on nuclear density, selecting NC14 embryos with regularly spaced nuclei (i.e., prior to significant nuclear clustering and fallout typical of late NC14 *zelda* mutants). To distinguish early and mid NC14 embryos, we relied on the severity of the *halo* mutant phenotype typical of embryos with inhibited zygotic genome activation. Sorted embryos were collected in 1.5 ml tubes, excess media was removed, and samples were frozen at - 80°C until needed.

For a particular experiment, a single biological replicate consisted of a 10-embryo equivalent of sheared chromatin produced from a batch of staged crosslinked embryos. Compared with prior ChIP-seq approaches (e.g., (Blythe & Wieschaus, 2015)), we amended as follows our sample preparation protocol to significantly increase the sensitivity of the approach by preparing ChIPped DNA ends by exposure to Tn5 transposase (tagmentation) prior to the typical wash steps of an immunoprecipitation. Aliquots of staged crosslinked embryos were resuspended in 700 µl of RIPA buffer (R0278 Sigma-Aldrich, Missouri, USA) supplemented with 1x protease inhibitor cocktail (Sigma P8340) and 1 mM Dithiothreitol. Chromatin was partly sheared by performing four 15-second rounds of sonication with a Branson 450 sonicator equipped with a 1/4-inch microtip horn (20% output, full duty cycle). 10-embryo equivalents of sheared chromatin were distributed to 1.5 ml DNA Lo-Bind tubes (Eppendorf, Hamburg, Germany) and brought to 650 µl final volume with RIPA buffer supplemented with protease inhibitors. Primary antibodies were added to the sheared chromatin preparations and incubated overnight at 4°C on a nutator. For each planned sequencing run, several biological replicates of a negative control ChIP were performed. Two negative control strategies were employed for this study. In most cases, an aliquot of sheared chromatin was immunoprecipitated with a rabbit polyclonal antibody to the c-myc epitope tag, which is absent from any of the genotypes in this study. Alternatively, for experiments measuring enrichment of GFP-tagged chimeric proteins, a rabbit polyclonal antibody to GFP was used as a negative control on wild type (*w[1118]*) chromatin. Protein G Dynabeads (15 µl per ChIP, Thermo-Fisher, Massachusetts, USA) were blocked on a nutator overnight in a solution of 1x PBS, 0.1% Triton X-100, 3% Bovine Serum Albumin (Sigma), and 0.2 mg/ml Yeast tRNA (Roche, Basel, Switzerland). The following morning, 15 µl of blocked Dynabeads were added to the sheared chromatin samples and incubated for one hour. Dynabeads were then immobilized on a magnetic stand and the captured material was exchanged into Tagmentation Wash Buffer (10 mM Tris pH 7.5, 5 mM MgCl_2_) by performing two 1 ml washes. Samples were then subjected to tagmentation by resuspending the Dynabead complexes in 20 µl of 1x Tn5 reaction buffer containing 1 µl of Tn5 transposase (Nextera, Illumina, California, USA) and incubating for 40 minutes in an Eppendorf Thermomixer set at 37°C with 1000 RPM shaking. Next, Dynabead complexes were subjected to five-minute washes, twice in “Wash buffer I” (20 mM Tris pH8.0, 0.1% SDS, 1% Triton X-100, 2mM EDTA, 150 mM NaCl), twice in “Wash buffer II” (Wash buffer I with 500 mM NaCl), and once in TE buffer (10 mM Tris pH 8.0, 1 mM EDTA). Tagmented, immunoprecipitated DNA fragments were eluted from the Dynabeads and were crosslink-reversed by incubation in “Elution Buffer” (50 mM Tris pH8.0, 10 mM EDTA, 1% SDS) containing 100 µg of Proteinase K (New England Biolabs, Massachusetts, USA) for four hours in an Eppendorf Thermomixer set at 65°C with 1500 RPM shaking. The eluted DNA fragments were purified using a Qiagen MinElute Enzyme Reaction Cleanup kit, performing one extra PE wash for three minutes and eluting after a three-minute incubation in 22.5 µl Qiagen Buffer EB.

### ChIP-seq Library Preparation and Sequencing

Following purification of tagmented ChIP samples, library preparation was performed by setting up PCRs containing 22.5 µl eluted ChIP DNA, 25 µl NEBNext Q5 2x PCR Mastermix (New England Biolabs), and 2.5 µl of a unique-dual-indexed primer mix (25 µM each primer). Primer designs were as previously reported (Soluri et al., 2020). Before performing any thermal cycling, reactions were mixed well and centrifuged, and a 5 µl aliquot of the primary PCR solution (pre-amplification) was added to a second, “side-reaction” that consisted of 5 µl primary PCR solution, 5 µl NEBNext Q5 2x PCR Mastermix, and 5 µl of a solution of 1.25 µM unique-dual-indexed primer mix in 1.8x SYBR-Green I (Thermo-Fisher). The primary PCR samples were stored at 4°C while the side-reaction was performed.

The side-reaction was subjected to QPCR to determine the total number of cycles necessary to amplify the primary PCR using the following cycling parameters: 1x 72°C, 5 minutes; 1x 98°C, 30 seconds; 20x (98°C, 10 seconds; 63°C, 30 seconds; 72°C 30 seconds). The total number of cycles to apply to the primary PCR was determined by finding the side-reaction cycle number (*N*) necessary to yield QPCR fluorescence intensity equal to 1/3 of the plateau fluorescence intensity. Based on these measurements, the primary PCR samples were then amplified using the following cycling parameters: 1x 72°C, 5 minutes; 1x 98°C, 30 seconds; *N*x (98°C, 10 seconds; 63°C, 30 seconds; 72°C 30 seconds); 1x 72°C, 5 minutes.

Barcoded DNA from the primary PCRs was purified by 1.8x Ampure, following the manufacturer’s instructions. DNA concentrations were determined by Qubit fluorometry using 1x HS DNA reagents (Thermo Fisher). The size-distribution of libraries was determined by TapeStation analysis using D5000 screen tapes (Agilent Technologies, California, USA).

Up to 48 libraries were pooled together in equimolar concentrations and submitted to Admera Health (South Plainfield, New Jersey, USA) for paired-end (2x 150 bp) sequencing using an Illumina NovaSeqX platform. One early dataset, RNA Pol2 CTD-pSer5 ChIP in wild type versus *zelda* was sequenced on a NextSeq500 platform at Northwestern University.

### ChIP-seq data processing and read mapping

ChIP-seq data were adaptor-trimmed and mapped on Northwestern University’s QUEST high-performance computing cluster. Demultiplexed fastq files were subjected to adaptor-trimming using TrimGalore v.0.6.10 (https://github.com/FelixKrueger/TrimGalore) and cutadapt v4.4 (Martin, 2011) and mapped to the *Drosophila melanogaster* dm6 reference genome assembly using bowtie2 v2.4.1 (Langmead & Salzberg, 2012) with option-X 2000. Suspected optical and PCR duplicates were marked by Picard MarkDuplicates version 2.21.4 (https://broadinstitute.github.io/picard/) using default parameters. Trimmed, mapped, and duplicate-marked reads were imported into R using the GenomicAlignments (version 1.38.2, (Lawrence et al., 2013)) and Rsamtools (version 2.18.0, (Morgan et al., 2020)) packages. Properly paired reads with primary mappings with map-quality scores greater than or equal to 10 were imported. Only reads corresponding to chromosome-level assemblies (chr2L, 2R, 3L, 3R, X) were considered further. Coverage plots (wiggle plots) were generated using the GenomicRanges package (version 1.42.0, (Lawrence et al., 2013)).

### Peak calling

Regions of enrichment for transcription factor ChIP targets were called using MACS2 v2.2.7.1 (Zhang et al., 2008) on merged replicate samples, using a GFP ChIP in wild type embryos as a negative control. In practice, peak calling results from initial adjusted p-value thresholds of q = 1e-5 were visually inspected against coverage plots for the ChIP target and negative-control ChIP to determine whether the selected adjusted p-value threshold adequately captured specific enrichment of the target factor. If q = 1e-5 was found to produce less than optimal results, the threshold was changed and the process repeated until the peak calling results were deemed by the user to accurately reflect biologically meaningful enrichments. The following are the final chosen q-value thresholds: E(z), 1e-7; Pho, 1e-40; Cg, 1e-30; GAF, 1e-5. Zelda peaks were generated by converting reported peaks (Harrison et al., 2011) from the dm3 to the dm6 reference genome using UCSC LiftOver in the rtracklayer package (version 1.50.0, (Lawrence et al., 2009)).

Broad regions of enrichment for H3K27me3 ChIP experiments were determined as described previously (Bonnet et al., 2019) using the STAN package in R (version 2.18.0, (Zacher et al., 2017)). H3K27me3 ChIP-seq coverage and negative control ChIP-seq coverage from late NC14 wild type samples were used for classifying genomic regions into three states (low me3/low background, high me3/low background, and high background). The high me3/low background state was processed further. To merge contiguou tracts of high H3K27me3 signal, genomic regions in the high me3/low background state were widened b 20 kbp, overlapping regions were merged, and 20 kb was subtracted from the resulting merged regions. Merged regions less than 4 kb in width were dropped from the list. This yielded a mapping of provisional Polycomb Domains. Following visual examination of the provisional mappings, a small number of regions corresponding to low-magnitude enrichment of H3K27me3 ChIP-seq coverage were dropped to simplify downstream analysis.

### Differential Enrichment Analysis

The procedure for performing differential enrichment analysis for H3K27me3 between wild type, Jabba-trap GAF and *zelda* germline clone embryos is presented in detail in the Source Code 1. Briefly, the dm6 reference genome assembly was subdivided into 2 kb bins spanning the canonical chromosome assemblies and the number of reads in each mapped ChIP sample overlapping with each bin wa tabulated. Bins with low read counts were filtered out and differential enrichment between genotypes wa calculated using DESeq2 (version 1.30.1, (Love et al., 2014)) using formulae with the form *∼ batch + genotype*, where “*batch*” reflected any batch effects identified through a preliminary principal component analysis. To identify consecutive runs of bins with statistically significant changes in ChIP signal, the DESeq results including adjusted p-values and log2fold changes were appended to a GenomicRange object representing the genomic coordinates of each associated bin, and bins with adjusted p-value greater than 0.05 were filtered out. The remaining bins with adjusted p-values less than 0.05 wer widened on each end by one bp to force overlaps with adjoining significant bins. Overlapping widened bins were then merged and one bp was removed from the ends of the remaining ranges after merging, yielding a mixture of new runs of consecutive significant bins and singleton bins. To propagate log2 fold-change values, the average log2 fold change of the bins comprising a run was calculated and appended to the run’s metadata. To combine p-value information, we applied Fisher’s method by calculating for *n* p-values (*p*) associated with bins in a run the right-tail χ-squared p-value for

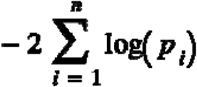

with 2*n* degrees of freedom (Yoon et al., 2021).

### Live Imaging

Embryos expressing an EGFP chimera (E(z), Pho, Cg, or GAF) and His2Av-RFP were collected from 0-2 hour plates and dechorionated in 4% sodium hypochlorite for 1 minute followed by extensive washing in deionized water. Dechorionated embryos were mounted in a small drop of Halocarbon 27 oil (Sigma-Aldrich) between a size 1.5 coverslip and a gas-permeable membrane (Biofoil, Heraeus, Hanau, Germany) stretched over an acrylic holder.

Mounted samples were imaged on a Leica SP8 laser scanning confocal microscope equipped with a white light laser using a 63x/1.3 NA glycerol-immersion objective. 512 x 256 pixel images were captured at 2x zoom with 400 Hz scan rate. Z-stacks were collected over the entire nuclear volume with a step size of 1 µm. Frames were collected for approximately two hours imaging every 30 seconds from metaphase of nuclear cycle 10 through one hour past metaphase 13 (approximately the start of gastrulation). Laser power was chosen to maximize intensity and minimize photobleaching in the GFP channel, as estimated by taking a zoomed-out overview image of the embryo following completion of the imaging session and verifying that bleaching was minimal. It was permissible for the His2Av-RFP imaging conditions to yield bleaching since this channel was only used for nuclear segmentation and not quantified.

To quantify GFP nuclear intensity, each time frame of a 3-dimensional image stack was segmented in the Histone RFP channel by first thresholding the image using Otsu’s method, filling in border pixels by dilation followed by erosion, and then breaking conjoined nuclei by using a watershed transformation. This yielded a nuclear mask from which the average pixel intensity in the GFP channel was calculated for each individual nucleus. This procedure was repeated over all frames of the movie. The average nuclear intensity for all nuclei in a frame was calculated, yielding the raw signal intensity for the movie. The raw signal intensity was next normalized to the mean intensity of the entire movie. To account for biological differences in nuclear cycle length between replicates, frames within individual nuclear cycles were binned to yield standard nuclear cycle durations of NC11 = 10 minutes, NC12 = 12.5 minutes, NC13 = 20 minutes, and NC14 = 60 minutes. Following binning for nuclear cycle length, average normalized intensities across replicates was calculated and plotted.

### Immunostaining

Immunostaining was performed on embryos fixed either by heat-methanol (H3K27me3) or formaldehyde (H2Aub, E(z), Pho, Cg, GAF) approaches. For either approach, embryos from 0-4 hour collections were first dechorionated in 4% Sodium Hypochlorite for 1-2 minutes followed by extensive washing in deionized water. Heat-Methanol fixation was performed by plunging dechorionated embryos into a 20 ml scintillation vial containing 5 ml of boiling 0.3% Triton X-100 + 0.4% (w/v) NaCl for ten seconds before placing on ice to cool rapidly. After cooling, the solution was removed and replaced with 5 ml heptane and 5 ml methanol to ‘pop’ embryos from their vitelline membranes. Formaldehyde fixation was performed by adding dechorionated embryos to a biphasic solution of 5 ml Heptane: 4 ml 1x phosphate buffered saline containing 1 ml of 20% paraformaldehyde solution in a 20 ml scintillation vial. The solution was placed on an orbital shaker for 20 minutes with 250 RPM shaking. The lower aqueous phase was carefully removed and replaced with 5 ml of methanol to “pop” embryos from their vitelline membranes. For both fixation methods, popped embryos were collected from the bottom of the scintillation vial and washed 3x in methanol prior to freezing for ≥ 24 hours in methanol. In preliminary trials for a particular antigen target, both fixation methods were performed, and a particular approach was chosen based on how well an antigen target would be stained under each fixation condition. For reasons that are not clear, formaldehyde fixation yielded poor signal even in late-stage embryos for H3K27me3 stainings, with excessive cytoplasmic background. Heat-methanol fixation greatly reduced Cg signals and caused excessive background with the H2Aub staining. Formaldehyde fixation was chosen for E(z), Pho, Cg, and GAF to maintain similar fixation conditions for these four factors which were to be compared directly.

The immunostaining procedure involved rehydration of fixed embryos in PTx (1x phosphate buffered saline + 0.1% Triton X-100), followed by one hour of blocking in PTx + 5% Bovine Serum Albumin. The primary antibody incubation was performed in blocking buffer overnight at 4°C on a nutator. Anti GFP, H2Aub, and H3K27me3 were used at a 1:500 dilution. Following the primary incubation, samples were washed for 40 minutes in four changes of PTx + 1% Bovine Serum Albumin. Secondary antibody incubations were performed in blocking buffer for two hours at room temperature on a nutator. The secondary antibody (goat anti rabbit coupled to alexa 546) was used at a 1:500 final dilution after pre-adsorption against fixed embryos at a 1:10 dilution. DAPI was added to 0.1 mg/ml final concentration during the final hour of secondary antibody incubation to stain DNA. If Alexa 647-coupled WGA (Invitrogen W32466) was included in a staining, it was added at a 1:100 final dilution at the same time as the DAPI treatment. Following the secondary incubation, samples were washed for 40 minutes in four changes of PTw (1x phosphate buffered saline + 0.1% Tween-20) prior to mounting embryos on slides in ProLong Gold (Thermo Fisher) with a size 1.5 coverslip, sealing prior to curing with clear nail polish.

Mounted samples were imaged on a Leica SP8 laser scanning confocal microscope equipped with a white light laser using a 63x/1.3 NA glycerol-immersion objective. 512×512 pixel images were captured at 2x zoom with 100 Hz scan rate. Z-stacks were collected over the entire nuclear volume with a step size of 1 µm. Alexa 546 conjugated secondary antibodies were excited at 557 nm. WGA coupled to Alexa 647, if used, was excited at 653 nm. DAPI was excited using a 405 nm laser.

### Antibodies

Antibodies were purchased from commercial sources and used as listed below.

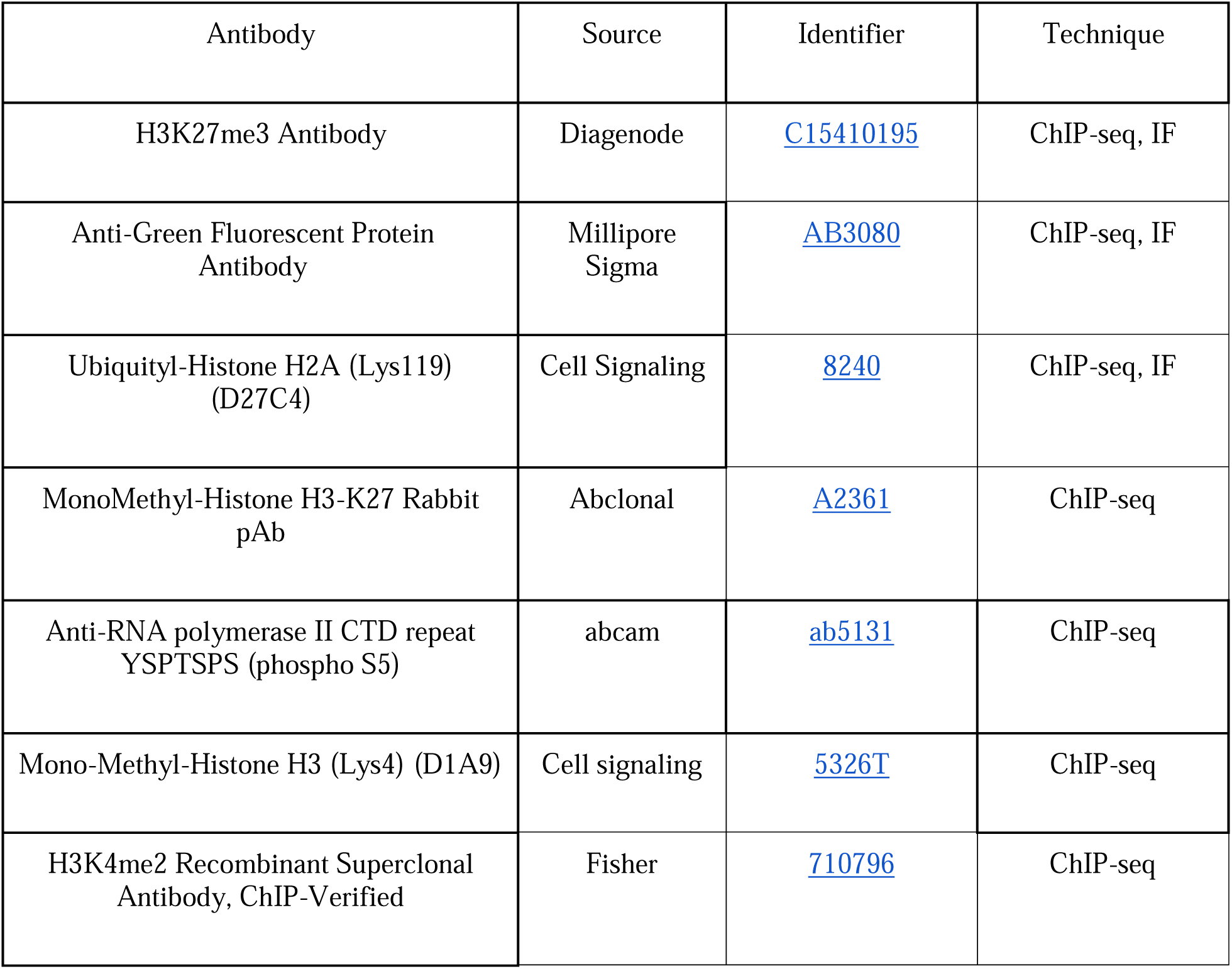
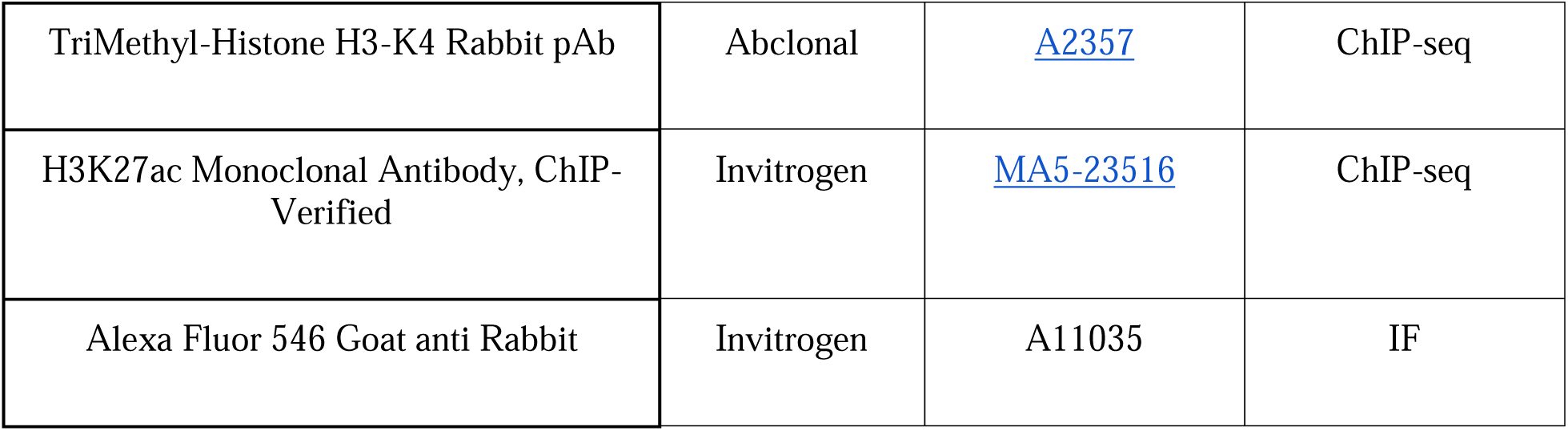

| Antibody | Source | Identifier | Technique |
| --- | --- | --- | --- |
| H3K27me3 Antibody | Diagenode | C15410195 | ChIP-seq, IF |
| Anti-Green Fluorescent Protein Antibody | Millipore Sigma | AB3080 | ChIP-seq, IF |
| Ubiquityl-Histone H2A (Lys119) (D27C4) | Cell Signaling | 8240 | ChIP-seq, IF |
| MonoMethyl-Histone H3-K27 Rabbit pAb | Abclonal | A2361 | ChIP-seq |
| Anti-RNA polymerase II CTD repeat YSPTSPS (phospho S5) | abcam | ab5131 | ChIP-seq |
| Mono-Methyl-Histone H3 (Lys4) (D1A9) | Cell signaling | 5326T | ChIP-seq |
| H3K4me2 Recombinant Superclonal Antibody, ChIP-Verified | Fisher | 710796 | ChIP-seq |
| TriMethyl-Histone H3-K4 Rabbit pAb | Abclonal | A2357 | ChIP-seq |
| H3K27ac Monoclonal Antibody, ChIP-Verified | Invitrogen | MA5-23516 | ChIP-seq |
| Alexa Fluor 546 Goat anti Rabbit | Invitrogen | A11035 | IF |

The anti H3K27me1 and me3 antisera chosen for this study have previously been subjected to specificity testing using the methodology reported here (Shah et al., 2018). Specificity data were recovered from https://www.chromatinantibodies.com.

## Acknowledgments

We thank Eric Wieschaus for comments and critiques on an earlier version of this manuscript. We thank Paul Schedl and Tsutomu Aoki for generously providing the *attP-3xMyc-Trl-loxP* stock, and Yue Yang and Jason Brickner for sharing access to instrumentation for sequencing and library preparation. We thank the Bloomington Drosophila Stock Center for providing stocks and Flybase for providing an essential resource to the *Drosophila* community. N.G-S., E.A.D., and C.C. were supported by the Cellular and Molecular Basis of Disease training program (T32 GM008061). E.A.D. is supported by an NSF GRFP fellowship (DGE-2234667) and is a Data Science Fellow at the Northwestern Institute on Complex Systems. Experiments were supported by the National Institutes of Health grant R01 HD101563 to S.A.B.. S.A.B. is a Pew Scholar in the Biomedical Sciences, supported by the Pew Charitable Trusts.

---

**Table 1:** Enrichment of Selected Transcription Factors with E(z) peaks that accumulate H3K27me3 by late NC14. Peaks of transcription factors (column 1) overlapping with E(z) peaks were counted and scored for whether overlapping E(z) peaks acquire H3K27me3 by late NC14. 2×2 contingency tables (factor-by-K27me3 domain membership, for the set of all E(z) peaks) were calculated and subjected to Fisher’s Exact Test. The number of factor peaks overlapping any E(z) peak (n = 4576) is shown in column 2. The number of factor peaks overlapping E(z) peaks within PcG domains (n = 1264) is shown in column 3. Log2(Odds Ratios) are reported in column 4, and p-values are reported in column 5. Negative log2(odds ratios) indicate depletion (red) and positive log2(odds ratios) indicate enrichment (blue) of a factor associating with E(z) peaks that accumulate H3K27me3. Additional pairwise comparisons that did not yield significant p-values are not shown.

| Factor | Number of Factor Peaks<br>overlapping any E(z)<br>Peak (%) | Number of Factor Peaks<br>overlapping an E(z) peak in<br>a PcG domain (%) | Enrichment in E(z)<br>PcG domains:<br>(log2[Odds Ratio]) | Enrichment in E(z)<br>PcG domains: (p-value) |
| --- | --- | --- | --- | --- |
| Pho | 1895 (41.4%) | 575 (45.5%) | 0.33 | 6e-4 |
| Combgap | 1691 (37.0%) | 321 (25.4%) | -1.05 | 2e-24 |
| GAGA-Factor | 3318 (72.5%) | 746 (59.0%) | -1.27 | 6e-35 |
| Combgap + GAGA-Factor | 1548 (33.8%) | 281 (22.2%) | -1.12 | 1e-25 |
| Zelda | 2348 (51.3%) | 828 (65.5%) | 1.16 | 8e-33 |
| Pho + Zelda | 1041 (22.7%) | 455 (36.0%) | 1.39 | 1e-37 |
| GAGA-Factor + Zelda | 2002 (43.8%) | 617 (48.8%) | 0.41 | 2e-5 |

**Figure 2-figure supplement 1: Transcription start site association of classes of E(z) peaks.** The number of E(z) peaks inside and outside of PcG Domains are shown in the bar graph, subset by whether a peak directly overlaps with an annotated transcription start site. We note that in comparison to the heatmaps this overlap measurement likely under-estimates the degree of overlap outside of PcG domains. This could be due to under-counting of active TSS, and/or a peak not strictly overlapping a TSS, but instead being near-by. Nevertheless, E(z) peaks outside of domains are significantly enriched for annotated active TSS regions with a log2 odds ratio of 2.5 and a Fisher’s exact p-value of 5.31e-105.

**Figure 3-figure supplement 1: Immunostaining of E(z) and three PRE-binding factors.** Embryos expressing EGFP-E(z) (**A**), Pho-sfGFP (**B**), EGFP-Cg (**C**), and EGFP-GAF (**D**) chimeras were formaldehyde-fixed and subjected to immunostaining with an anti-GFP antibody to facilitate measurement of nuclear concentration prior to nuclear migration. All images are maximum z-projections. For each chimera, the top row of images show GFP immunofluorescence signal from representative embryos staged at the nuclear cycle indicated at top. Samples were co-stained with DAPI to visualize DNA/nuclei. The insets show an enlarged single nucleus indicated by the yellow arrowhead (green: GFP chimera, magenta: DAPI). Displayed image intensities are consistent for post-migration nuclear cycles (NC10-14) and are linearly adjusted based on DAPI intensity for pre-migration nuclear cycles, when nuclei are found deep within the embryo. For EGFP-GAF (D), one example of a pre-migration mitosis is shown (NC7 anaphase) to demonstrate the ready detection of GAF on mitotic chromatin in contrast to much lower signal during pre-migration interphases. E(z) localizes to interphase nuclei from the beginning of the cleavage divisions and is readily detected. Pho, Cg, and GAF do not localize to interphase nuclei until after nuclear migration. The scale bar is shown in yellow in the bottom right of the top-left image in each panel, size = 10 µm.

**Figure 5-figure supplement 1: DESeq on 2 kb bins for the GAF knockdown experiment and making runs out of bins. A**) A volcano plot is shown for DESeq2 analysis of H3K27me3 ChIP-seq for EGFP-GAF versus JT-GAF. Overall, very few 2 kb bins show effects that surpass the significance thresholds for p-value and log2(fold change). The bin corresponding to the escargot transcription start site is labeled. **B**) Reproduced from Figure 5 with additional annotations indicating the location of the highlighted point in panel A and a broader region surrounding the bin with a more moderate effect on H3K27me3 following JT-sequestration of GAF. **C**) Strategy for generating consecutive 2 kb runs of bins with p-values < 0.05, and how fold-change and significance values are propagated. **D**) Reproduced from Figure 5 to show how making consecutive runs transforms the data.

**Figure 6-figure supplement 1: DESeq on 2 kb bins for the *zelda* mutant experiment and correlation between H3K27me3 and H2Aub. A**) A volcano plot is shown for DESeq analysis performed on H3K27me3 ChIP seq between *zelda* mutants and wild type embryos, staged mid NC14. 226 bins have significant reductions in H3K27me3, and 62 bins have significant increases. Significant bins containing a zygotic transcription start site are labeled with the gene symbol (red). **B**) Log2(fold change) values from DESeq analysis of H3K27me3 (x-axis) and H2Aub (y-axis) ChIP-seq between *zelda* and wild type embryos are plotted. The correlation coefficient is 0.78. Linear regression was performed, and the fit model is plotted in red (slope = 0.71). There is a strong positive correlation between changes in both modifications following loss of *zelda*, although the trend is for H2Aub to lag slightly in its magnitude of response compared with H3K27me3.

## Supplementary Information

Supplementary File 1: Annotated Polycomb Domains.

Supplementary File 2: Annotated E(z) Peaks List.

Supplementary File 3: Annotated Pho Peaks List.

Supplementary File 4: Annotated Cg Peaks List.

Supplementary File 5: Annotated GAF Peaks List.

Supplementary File 6: Annotated Zld Peaks List.

Source Code 1: Genomic Analysis Supplement

## Supplementary Movie Legends

Figure 3-video 1: His2Av-RFP; EGFP-E(z) NC10 to 60 minutes into NC14. A representative movie of (top) EGFP-E(z) and (bottom) a merged image with His2Av-RFP (magenta) and EGFP-E(z) (yellow) is shown. E(z) localizes to interphase nuclei throughout the period of NC10-NC14.

Figure 3-video 2: His2Av-RFP; Pho-sfGFP NC10 to 60 minutes into NC14. A representative movie of (top) Pho-sfGFP and (bottom) a merged image with His2Av-RFP (magenta) and Pho-sfGFP (yellow) is shown. Pho is initially excluded from nuclei and increases in concentration between NC11 and NC14.

Figure 3-video 3: His2Av-RFP; EGFP-Cg NC10 to 60 minutes into NC14. A representative movie of (top) EGFP-Cg and (bottom) a merged image with His2Av-RFP (magenta) and EGFP-Cg (yellow) is shown. Cg is initially excluded from nuclei and increases in concentration between NC12 and NC14.

Figure 3-video 4: His2Av-RFP; EGFP-GAF NC10 to 60 minutes into NC14. A representative movie of (top) EGFP-GAF and (bottom) a merged image with His2Av-RFP (magenta) and EGFP-GAF (yellow) is shown. GAF initially only localizes to pericentric GA-rich regions before increasing nucleoplasmic concentration between NC11 and 14.

Figure 5-video 1: His2Av-RFP; EGFP-GAF NC10 through NC14, control for Jabba-Trap knockdown experiment. This is a control movie to be compared with Figure 5-video 6, imaged under identical conditions. The movie shows a single z-slice of an His2Av-RFP; EGFP-GAF embryo from NC10 to NC14, with single-channel EGFP-GAF at left (grey), and the merged image at right (green = EGFP-GAF).

Figure 5-video 2: His2Av-RFP; Jabba-Trap/+; EGFP-GAF NC10 through NC14. This movie shows the effect of Jabba-Trap sequestration on EGFP-GAF in a single z-slice of an His2Av-RFP; Jabba-Trap/+; EGFP-GAF embryo, with single-channel EGFP-GAF at left (grey), and the merged image at right (green = EGFP-GAF).

## Supplementary File Legends

Supplementary File 1: Annotated Polycomb Domains. The list of PcG Domains used in the text is presented in tab-delimited format with additional annotations indicating co-occurrence with genomic features of interest. The columns are: Row number (numeric), Seqnames (Chromosome, character), Start (numeric), End (numeric), Width (numeric), Strand (character), Domain Name (character), with E(z) (logical), with Cg (logical), with GAF (logical), with Zld (logical), with a Maternal/Zygotic transcription start site (logical), Maternal/Zygotic transcription start site gene name (character), with Zygotic-only transcription start site (logical), Zygotic-only transcription start site gene name (character).

Supplementary File 2: Annotated E(z) Peaks. The list of E(z) peaks used in the text is presented in tab-delimited format with additional annotations indicating co-occurrence with genomic features of interest. The columns are: Row number (numeric), Seqnames (Chromosome, character), Start (numeric), End (numeric), Width (numeric), Strand (character), MACS Signal Value (numeric), MACS q-value (numeric), peak summit (numeric), within a PcG Domain (logical), PcG Domain Name (character), Maximal E(z) peak in Domain (logical), with Pho (logical), with Cg (logical), with GAF (logical), with Zld (logical), with a Maternal/Zygotic transcription start site (logical), Maternal/Zygotic transcription start site gene name (character), with a Zygotic-only transcription start site (logical), Zygotic-only transcription start site name (character).

Supplementary File 3: Annotated Pho Peaks. The list of Pho peaks used in the text is presented in tab-delimited format with additional annotations indicating co-occurrence with genomic features of interest. The columns are: Row number (numeric), Seqnames (Chromosome, character), Start (numeric), End (numeric), Width (numeric), Strand (character), MACS Signal Value (numeric), MACS q-value (numeric), peak summit (numeric), within a PcG domain (logical), PcG Domain Name (logical), with E(z) (logical), with Cg (logical), with GAF (logical), with Zld (logical), with a Maternal/Zygotic transcription start site (logical), Maternal/Zygotic transcription start site gene name (character), with a Zygotic-only transcription start site (logical), Zygotic-only transcription start site name (character).

Supplementary File 4: Annotated Cg Peaks. The list of Cg peaks used in the text is presented in tab-delimited format with additional annotations indicating co-occurrence with genomic features of interest. The columns are: Row number (numeric), Seqnames (Chromosome, character), Start (numeric), End (numeric), Width (numeric), Strand (character), MACS Signal Value (numeric), MACS q-value (numeric), peak summit (numeric), within a PcG domain (logical), PcG Domain Name (logical), with E(z) (logical), with Pho (logical), with GAF (logical), with Zld (logical), with a Maternal/Zygotic transcription start site (logical), Maternal/Zygotic transcription start site gene name (character), with a Zygotic-only transcription start site (logical), Zygotic-only transcription start site name (character).

Supplementary File 5: Annotated GAF Peaks. The list of GAF peaks used in the text is presented in tab-delimited format with additional annotations indicating co-occurrence with genomic features of interest. The columns are: Row number (numeric), Seqnames (Chromosome, character), Start (numeric), End (numeric), Width (numeric), Strand (character), MACS Signal Value (numeric), MACS q-value (numeric), peak summit (numeric), within a PcG domain (logical), PcG Domain Name (logical), with E(z) (logical), with Pho (logical), with Cg (logical), with Zld (logical), with a Maternal/Zygotic transcription start site (logical), Maternal/Zygotic transcription start site gene name (character), with a Zygotic-only transcription start site (logical), Zygotic-only transcription start site name (character).

Supplementary File 6: Annotated Zelda Peaks. The list of 3h after egg laying Zelda peaks from (Harrison et al., 2011) was lifted-over to dm6 coordinates. Here we report this list in tab-delimited format with additional annotations indicating co-occurrence with genomic features of interest. The columns are: Row number (numeric), Seqnames (Chromosome, character), Start (numeric), End (numeric), Width (numeric), Strand (character), Name (character), Peak Score (numeric), within a PcG domain (logical), PcG Domain Name (logical), with E(z) (logical), with Pho (logical), with Cg (logical), with GAF (logical), with a Maternal/Zygotic transcription start site (logical), Maternal/Zygotic transcription start site gene name (character), with a Zygotic-only transcription start site (logical), Zygotic-only transcription start site name (character).

