## Supplementary figures and images for "Nucleation-dependent propagation of Polycomb modifications emerges during the *Drosophila* maternal to zygotic transition"

### Figure 2-figure supplement 1

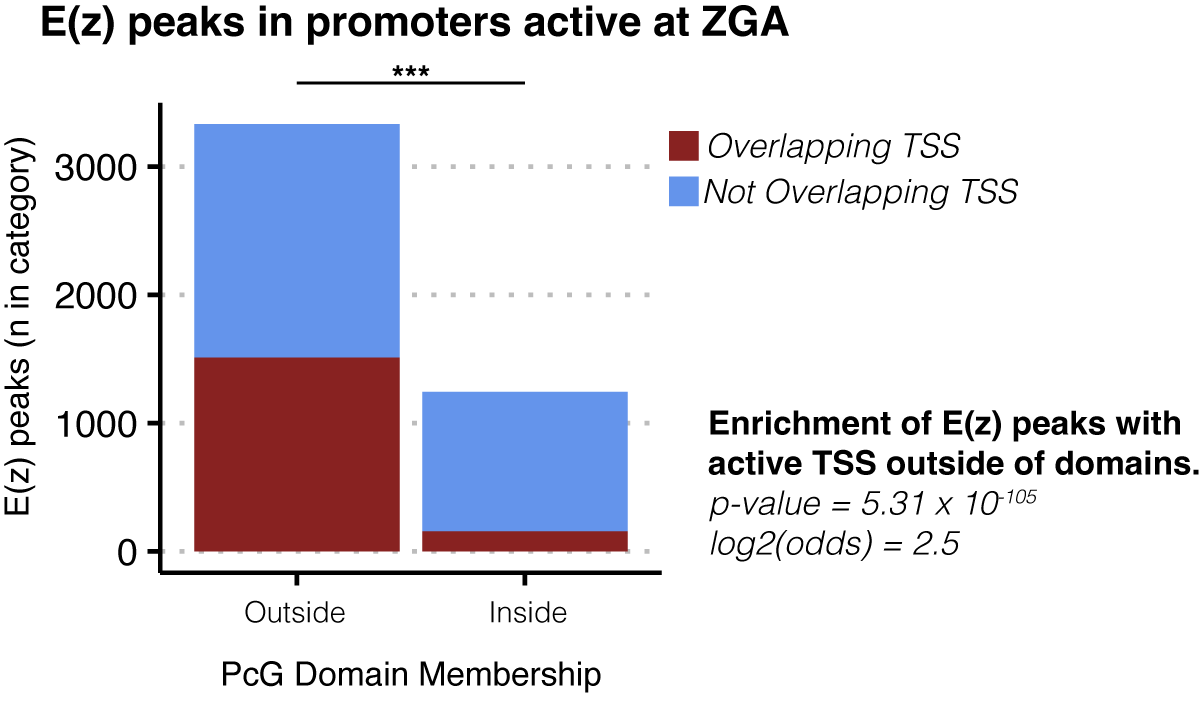

### Figure 3-figure supplement 1

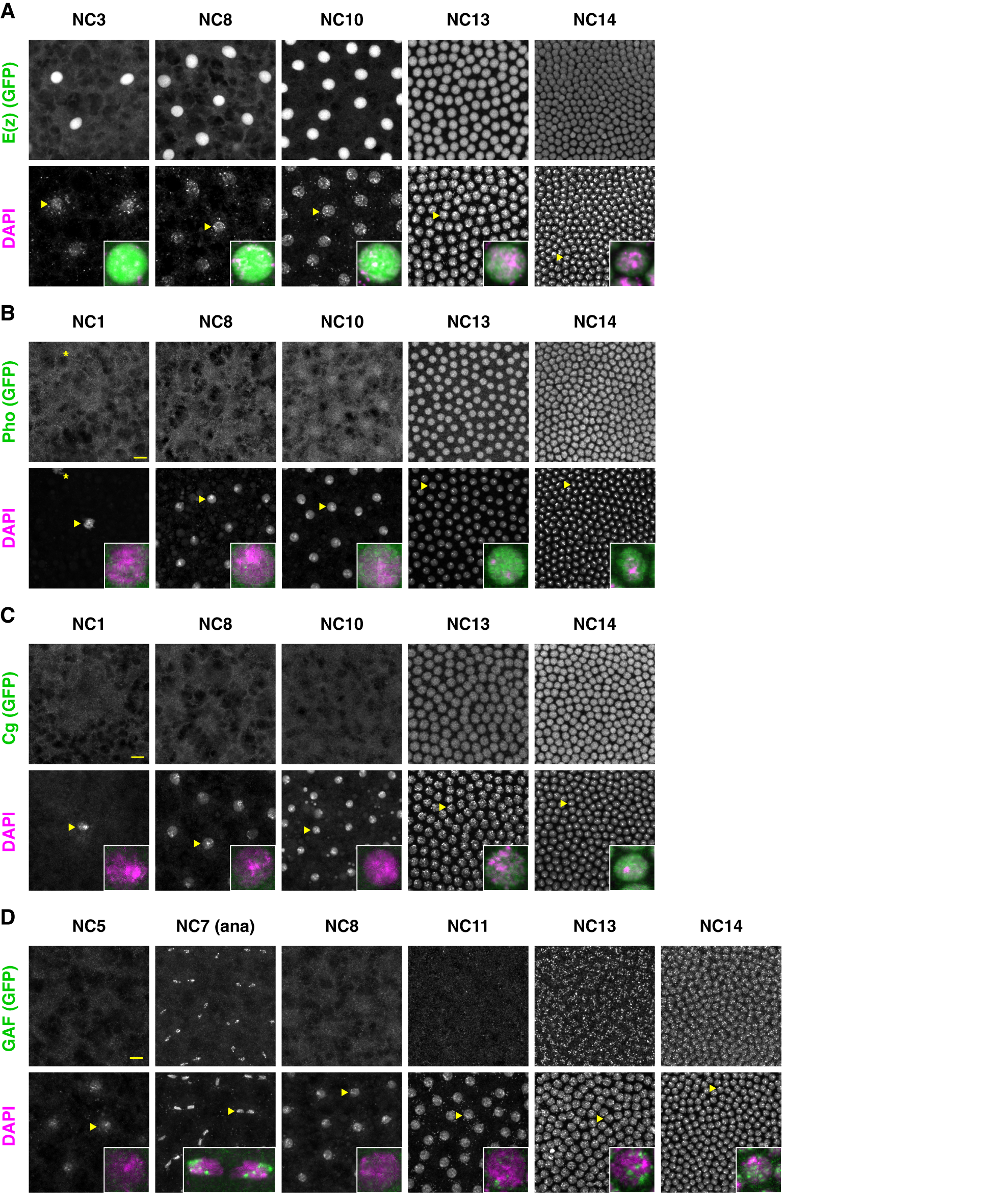

### Figure 5-figure supplement 1

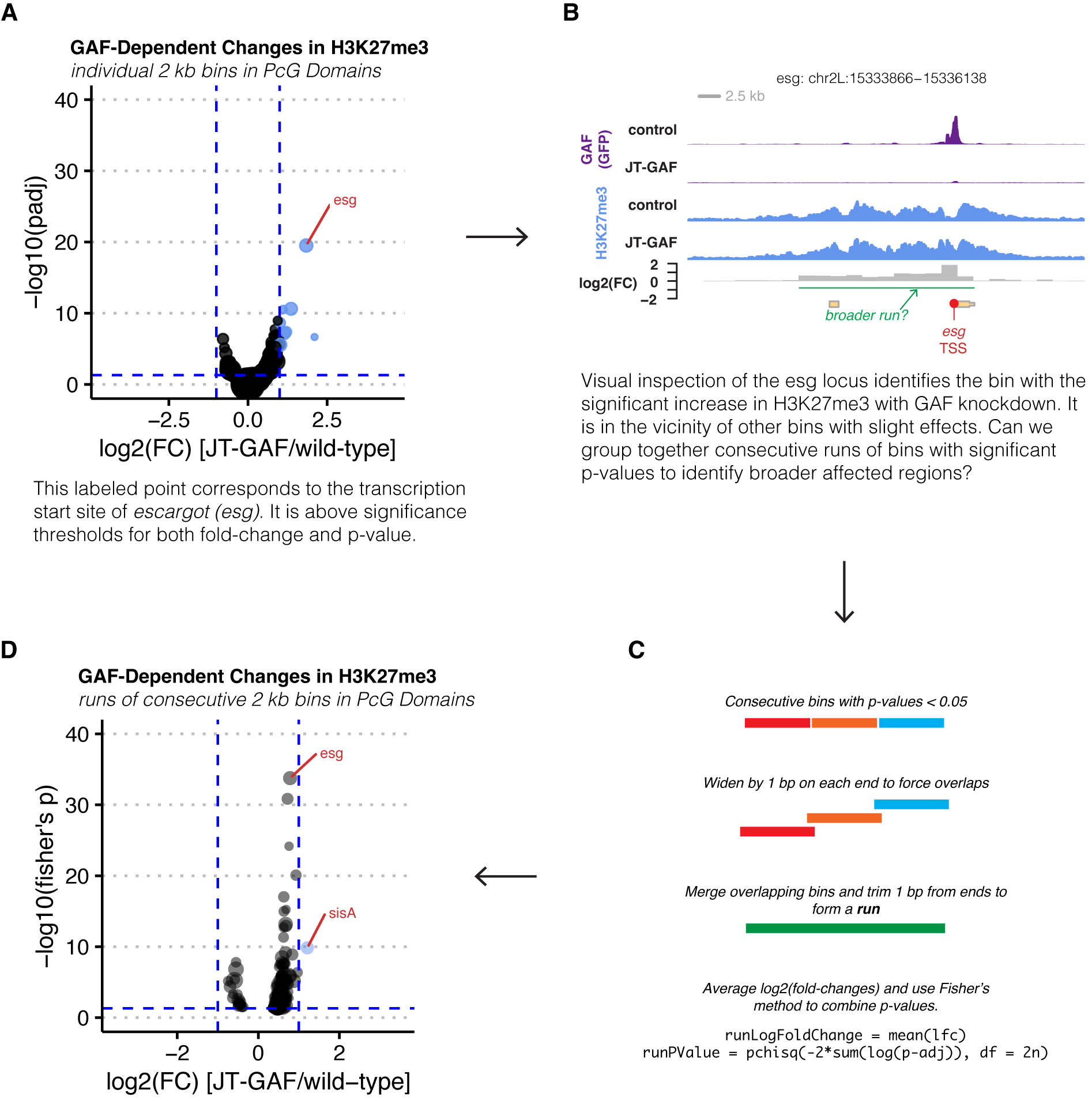

### Figure 6-figure supplement 1

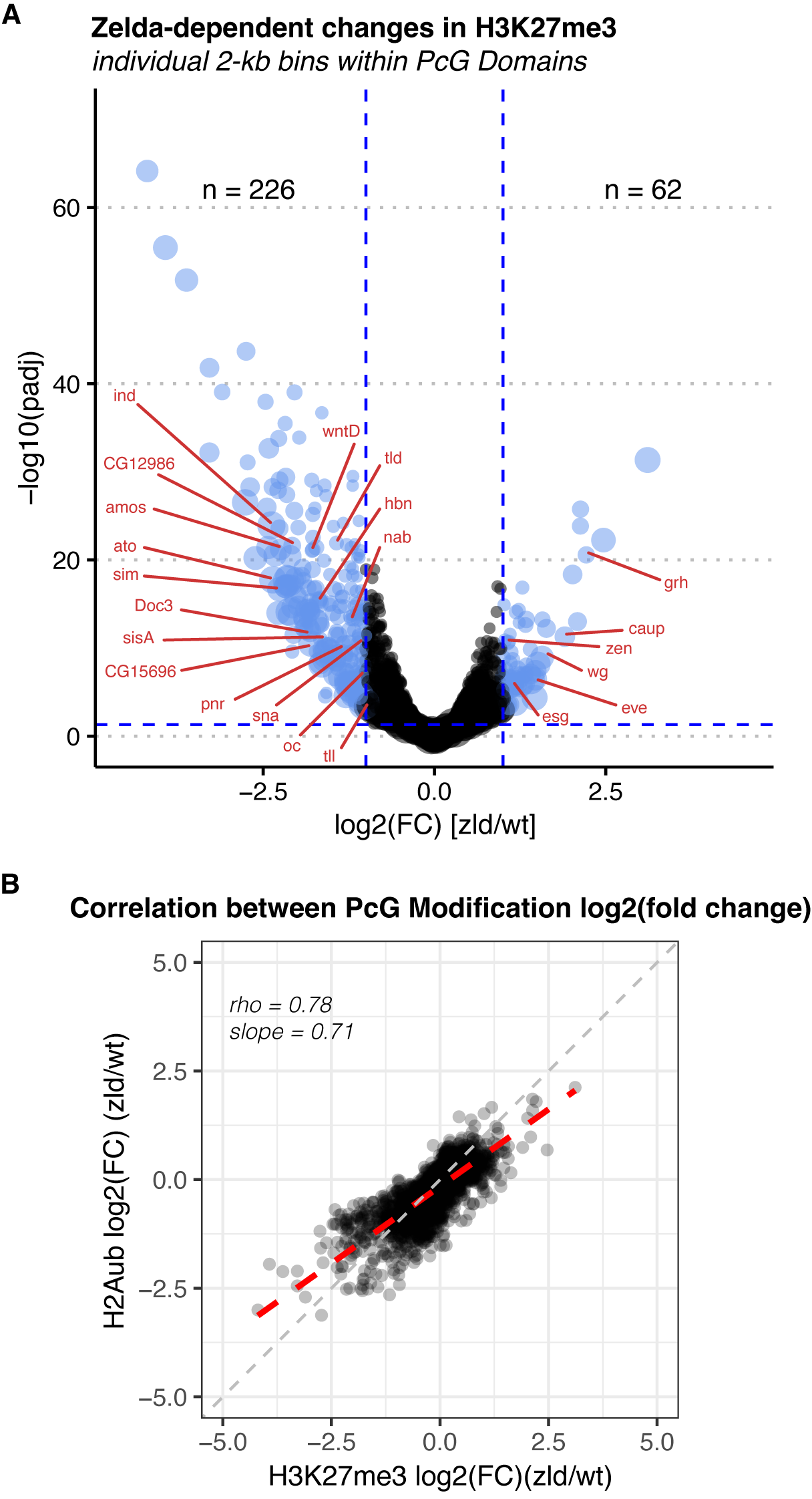
