## Supplementary material for "Nucleation-dependent propagation of Polycomb modifications emerges during the *Drosophila* maternal to zygotic transition": Source Code 1: Source_Code_1.html

Gonzaga-Saavedra 2025 Code Supplement


### Gonzaga-Saavedra 2025 Code Supplement

###### Shelby Blythe

#### 2025-06-07

This code supplement will reproduce the analyses reported in
Gonzaga-Saavedra 2025.

### Load Data

The necessary datasets for reproducing the data are loaded here. To
make this most portable, we have generated RDS formatted versions of
peaks lists that have been submitted to GEO and have provided counts
tables for the GAF and Zelda mutant/knockdown experiments to avoid
needing to repeat counting from raw .bam files.

The first filepath below should be changed to the relevant filepath
on your system where this repository has been cloned. Provided the
filenames of the data themselves do not change, the outcome of this
first chunk should be the error-free loading of the necessary
datasets.

```
rm(list = ls()); invisible(gc());

# load libraries and packages needed:

suppressPackageStartupMessages({
  library(GenomicRanges)
  library(tidyverse)
  library(DESeq2)
  library(BSgenome.Dmelanogaster.UCSC.dm6)
  library(Rsamtools)
  library(GenomicAlignments)
})

# Filepath to parental data directory:

d = "~/Dropbox/Blythe Lab/Manuscripts/Gonzaga-Saavedra/Github_Repo/Gonzaga_Saavedra_2025_Code_Supplement/Data/"

# E(z) peaks:

ez = readRDS(paste0(d, "Ez_peaks.RDS"))

# Pho peaks:

pho = readRDS(paste0(d, "Pho_peaks.RDS"))

# Cg peaks:

cg = readRDS(paste0(d, "Cg_peaks.RDS"))

# GAF peaks:

gaf = readRDS(paste0(d, "GAF_peaks.RDS"))

# Zld peaks (Harrison et al, lifted-over to dm6):

zld = readRDS(paste0(d, "Eisen_3h_Zld_peaks.RDS"))

# PcG Domains:

domains = readRDS(paste0(d, "NC14_PcG_Domains.RDS"))

# Pol2 engagement with zygotic only genes (Chen/Zeitlinger 2013)

zygtss = readRDS(paste0(d, "Chen_eLife_2013_zygotic_only_TSS.RDS"))

# Pol2 engagement with maternal-zygotic and zygotic-only genes (Chen/Zeitlinger 2013)

matzyg = readRDS(paste0(d, "Chen_eLife_2013_maternal_and_zygotic_TSS.RDS"))

# Sampling of ATAC data from NC11-NC13 to estimate timing of peak accessibility.

atac = readRDS(paste0(d, "NC11-13_ATAC.RDS"))

# count matrices for DESeq analysis

gafcts = readRDS(paste0(d, "count_matrices_gafKD.RDS"))
zldcts = readRDS(paste0(d, "count_matrices_zld.RDS"))

# the supplied gaf counts are just for the H3K27me3 experiment. The Zelda experiment on the other hand contains counts for H3K27me3, H2Aub, RNA Pol 2, H3K27ac, and H3K4me2. The following indexing variables identify the columns in the zelda experiment for each respective ChIP. 

k27I = c(1:8)
h2aI = c(9:12)
ezI = c(13:18)
polI = c(19:22)
acI = c(23:28)
k4I = c(29:32)

# Next we have some helper functions:

# the function below appends a UCSC-friendly name to each range

namer = function(cts){
    names(cts) = paste0(seqnames(cts), ":", start(cts), "-", end(cts))
    return(cts)
}

# the function below cleans up a GRanges object to be limited to canonical
# chromosomes, to all be the same width, and fixes the reference genome info
# appended to the object. 

cleanup = function(b, sizer = TRUE){
    b = b[seqnames(b) %in% paste0('chr', c('2L','2R','3L','3R','4','X'))]
    if(sizer){b = b[width(b) == median(width(b))]}
    seqlevels(b) = seqlevels(Dmelanogaster)
    seqinfo(b) = seqinfo(Dmelanogaster)
    seqlevels(b) = seqlevelsInUse(b)
    genome(b) = 'dm6'
    return(b)
}
```

Now that the data are loaded, we can perform calculations:

### Calculations

#### E(z) occurence in PcG Domains and co-occurrence with TF binding events

For the first section of the paper, we report numbers related to E(z)
occupancy in the genome and co-occurrence with selected transcription
factors.

The total number of E(z) peaks:

```
length(ez)
```

```
## [1] 4576
```

To calculate the fraction of these peaks that overlap with either
Pho, Cg, GAF, or Zld, we can append a logical vector reporting these
overlaps to the E(z) peaks list.

```
ez$withPho = as.logical(countOverlaps(ez, pho))
ez$withCg = as.logical(countOverlaps(ez, cg))
ez$withGAF = as.logical(countOverlaps(ez, gaf))
ez$withZld = as.logical(countOverlaps(ez, zld))
```

This method of counting overlaps just reports “any” overlap. One peak
for one factor can overlap with one or more of the other, and it is
scored as an overlap.

In the manuscript, we note the overall overlap of an E(z) peak with
one or more of any of these four factors. To calculate this:

```
table(ez$withPho | ez$withCg | ez$withGAF | ez$withZld)[2]
```

```
## TRUE 
## 3966
```

```
table(ez$withPho | ez$withCg | ez$withGAF | ez$withZld)[2]/length(ez)
```

```
##      TRUE 
## 0.8666958
```

There are 3966 peaks that overlap with one or more of these factors,
and this accounts for 87% of the E(z) peaks overall.

#### PcG domain features

We reckon “PcG Domains” using a HMM tool to identify tracts of high
H3K27me3 signal in NC14L chromatin. How many do we have?

```
length(domains)
```

```
## [1] 255
```

What is their median length?

```
median(width(domains))
```

```
## [1] 19201
```

Size range?

```
range(width(domains))
```

```
## [1]   4201 350001
```

Mean width

```
mean(width(domains))
```

```
## [1] 29349.24
```

How many domains contain one (or more) E(z) peaks?

```
table(as.logical(countOverlaps(domains, ez)))[2]
```

```
## TRUE 
##  237
```

```
table(as.logical(countOverlaps(domains, ez)))[2]/length(domains)
```

```
##      TRUE 
## 0.9294118
```

How many E(z) peaks are found within Domains?

```
table(as.logical(countOverlaps(ez, domains)))[2]
```

```
## TRUE 
## 1264
```

```
table(as.logical(countOverlaps(ez, domains)))[2]/length(ez)
```

```
##      TRUE 
## 0.2762238
```

We report the fraction not within domains:

```
table(as.logical(countOverlaps(ez, domains)))[1]
```

```
## FALSE 
##  3312
```

```
table(as.logical(countOverlaps(ez, domains)))[1]/length(ez)
```

```
##     FALSE 
## 0.7237762
```

#### E(z) peak engagement with TSS

We will now append “domain” and “tss” membership with each E(z)
peak.

```
ez$domain = as.logical(countOverlaps(ez, domains))
ez$tss = as.logical(countOverlaps(ez, matzyg))
```

This list of TSS is curated from Chen et al 2013, and contains both
the maternal-zygotic and zygotic-only TSS.

```
length(matzyg)
```

```
## [1] 3917
```

This accounts for 3917 total promoters.

What fraction of E(z) peaks associate with an active TSS?

```
table(ez$tss)
```

```
## 
## FALSE  TRUE 
##  2908  1668
```

There are 1668 E(z) peaks that engage with one or more active TSS at
NC14. Does it make a difference if the E(z) peak is in a domain or
not?

```
table(ez$tss, ez$domain)[c(2:1), c(2:1)]
```

```
##        
##         TRUE FALSE
##   TRUE   161  1507
##   FALSE 1103  1805
```

```
fisher.test(table(ez$tss, ez$domain)[c(2:1), c(2:1)])
```

```
## 
##  Fisher's Exact Test for Count Data
## 
## data:  table(ez$tss, ez$domain)[c(2:1), c(2:1)]
## p-value < 2.2e-16
## alternative hypothesis: true odds ratio is not equal to 1
## 95 percent confidence interval:
##  0.1452892 0.2095185
## sample estimates:
## odds ratio 
##  0.1748891
```

There is a substantial enrichment for E(z) binding outside of domains
at TSS sites. Here’s the exact p-value and log2 transformed odds
ratio:

```
fisher.test(table(ez$tss, ez$domain)[c(2:1), c(2:1)])$p.value
```

```
## [1] 5.314684e-105
```

```
log2(fisher.test(table(ez$tss, ez$domain)[c(2:1), c(2:1)])$estimate)
```

```
## odds ratio 
##  -2.515488
```

Because of the way that I phrased the contingency table for this
test, the resulting odds ratio reports “enrichment of *in domain*
and *is tss*” so we see a negative odds ratio indicating that
there is a strong enrichment for E(z) binding not in PcG domains at
TSS.

A stacked barplot with this information:

```
s = as.data.frame(ez) %>% 
    select(inDomain, tss) %>% 
    ggplot(aes(x = inDomain, fill = tss)) +
    geom_bar(position = 'stack', show.legend = FALSE) +
    scale_fill_manual(values = c("cornflowerblue","brown4")) +
    labs(x = "domain membership", y = "n E(z) peaks", title = "E(z) peaks in promoters active at ZGA") +
    ggthemes::theme_clean()
s
```

#### Table 1:

For table 1, we ask whether co-binding of E(z) with other
transcription factors disproportionately correlates with an E(z) peak
acquiring H3K27me3. We report the overlap of a factor with any E(z)
peak, with an E(z) peak within a domain, and a fisher’s exact test
result.

```
cat("Pho\n")
```

```
## Pho
```

```
test = ez$withPho

sum(test)
```

```
## [1] 1895
```

```
sum(test)/length(ez)
```

```
## [1] 0.4141171
```

```
sum(test[ez$domain])
```

```
## [1] 575
```

```
sum(test[ez$domain])/sum(ez$domain)
```

```
## [1] 0.4549051
```

```
fisher.test(table(test, ez$domain)[c(2:1), c(2:1)])$p.value
```

```
## [1] 0.0006157442
```

```
log2(fisher.test(table(test, ez$domain)[c(2:1), c(2:1)])$estimate)
```

```
## odds ratio 
##  0.3326473
```

```
cat("Cg\n")
```

```
## Cg
```

```
test = ez$withCg

sum(test)
```

```
## [1] 1691
```

```
sum(test)/length(ez)
```

```
## [1] 0.3695367
```

```
sum(test[ez$domain])
```

```
## [1] 321
```

```
sum(test[ez$domain])/sum(ez$domain)
```

```
## [1] 0.2539557
```

```
fisher.test(table(test, ez$domain)[c(2:1), c(2:1)])$p.value
```

```
## [1] 2.471405e-24
```

```
log2(fisher.test(table(test, ez$domain)[c(2:1), c(2:1)])$estimate)
```

```
## odds ratio 
##  -1.051104
```

```
cat("GAF\n")
```

```
## GAF
```

```
test = ez$withGAF

sum(test)
```

```
## [1] 3318
```

```
sum(test)/length(ez)
```

```
## [1] 0.7250874
```

```
sum(test[ez$domain])
```

```
## [1] 746
```

```
sum(test[ez$domain])/sum(ez$domain)
```

```
## [1] 0.5901899
```

```
fisher.test(table(test, ez$domain)[c(2:1), c(2:1)])$p.value
```

```
## [1] 6.340563e-35
```

```
log2(fisher.test(table(test, ez$domain)[c(2:1), c(2:1)])$estimate)
```

```
## odds ratio 
##  -1.270851
```

```
cat("Zelda\n")
```

```
## Zelda
```

```
test = ez$withZld

sum(test)
```

```
## [1] 2348
```

```
sum(test)/length(ez)
```

```
## [1] 0.5131119
```

```
sum(test[ez$domain])
```

```
## [1] 828
```

```
sum(test[ez$domain])/sum(ez$domain)
```

```
## [1] 0.6550633
```

```
fisher.test(table(test, ez$domain)[c(2:1), c(2:1)])$p.value
```

```
## [1] 7.927968e-33
```

```
log2(fisher.test(table(test, ez$domain)[c(2:1), c(2:1)])$estimate)
```

```
## odds ratio 
##   1.162554
```

```
cat("Pho and Zelda\n")
```

```
## Pho and Zelda
```

```
test = ez$withPho & ez$withZld

sum(test)
```

```
## [1] 1041
```

```
sum(test)/length(ez)
```

```
## [1] 0.2274913
```

```
sum(test[ez$domain])
```

```
## [1] 455
```

```
sum(test[ez$domain])/sum(ez$domain)
```

```
## [1] 0.3599684
```

```
fisher.test(table(test, ez$domain)[c(2:1), c(2:1)])$p.value
```

```
## [1] 1.442608e-37
```

```
log2(fisher.test(table(test, ez$domain)[c(2:1), c(2:1)])$estimate)
```

```
## odds ratio 
##   1.387137
```

```
cat("Pho and GAF\n")
```

```
## Pho and GAF
```

```
test = ez$withPho & ez$withGAF

sum(test)
```

```
## [1] 1573
```

```
sum(test)/length(ez)
```

```
## [1] 0.34375
```

```
sum(test[ez$domain])
```

```
## [1] 461
```

```
sum(test[ez$domain])/sum(ez$domain)
```

```
## [1] 0.3647152
```

```
fisher.test(table(test, ez$domain)[c(2:1), c(2:1)])$p.value
```

```
## [1] 0.07028793
```

```
log2(fisher.test(table(test, ez$domain)[c(2:1), c(2:1)])$estimate)
```

```
## odds ratio 
##  0.1836663
```

```
cat("Pho and Cg\n")
```

```
## Pho and Cg
```

```
test = ez$withPho & ez$withCg

sum(test)
```

```
## [1] 937
```

```
sum(test)/length(ez)
```

```
## [1] 0.204764
```

```
sum(test[ez$domain])
```

```
## [1] 249
```

```
sum(test[ez$domain])/sum(ez$domain)
```

```
## [1] 0.1969937
```

```
fisher.test(table(test, ez$domain)[c(2:1), c(2:1)])$p.value
```

```
## [1] 0.4364827
```

```
log2(fisher.test(table(test, ez$domain)[c(2:1), c(2:1)])$estimate)
```

```
##  odds ratio 
## -0.09596462
```

```
cat("Zld and GAF\n")
```

```
## Zld and GAF
```

```
test = ez$withZld & ez$withGAF

sum(test)
```

```
## [1] 2002
```

```
sum(test)/length(ez)
```

```
## [1] 0.4375
```

```
sum(test[ez$domain])
```

```
## [1] 617
```

```
sum(test[ez$domain])/sum(ez$domain)
```

```
## [1] 0.4881329
```

```
fisher.test(table(test, ez$domain)[c(2:1), c(2:1)])$p.value
```

```
## [1] 2.29002e-05
```

```
log2(fisher.test(table(test, ez$domain)[c(2:1), c(2:1)])$estimate)
```

```
## odds ratio 
##  0.4078636
```

```
cat("Zld and Cg\n")
```

```
## Zld and Cg
```

```
test = ez$withZld & ez$withCg

sum(test)
```

```
## [1] 1053
```

```
sum(test)/length(ez)
```

```
## [1] 0.2301136
```

```
sum(test[ez$domain])
```

```
## [1] 260
```

```
sum(test[ez$domain])/sum(ez$domain)
```

```
## [1] 0.2056962
```

```
fisher.test(table(test, ez$domain)[c(2:1), c(2:1)])$p.value
```

```
## [1] 0.01654655
```

```
log2(fisher.test(table(test, ez$domain)[c(2:1), c(2:1)])$estimate)
```

```
## odds ratio 
## -0.2816318
```

```
cat("Cg and GAF\n")
```

```
## Cg and GAF
```

```
test = ez$withCg & ez$withGAF
sum(test)
```

```
## [1] 1548
```

```
sum(test)/length(ez)
```

```
## [1] 0.3382867
```

```
sum(test[ez$domain])
```

```
## [1] 281
```

```
sum(test[ez$domain])/sum(ez$domain)
```

```
## [1] 0.2223101
```

```
fisher.test(table(test, ez$domain)[c(2:1), c(2:1)])$p.value
```

```
## [1] 1.271649e-25
```

```
log2(fisher.test(table(test, ez$domain)[c(2:1), c(2:1)])$estimate)
```

```
## odds ratio 
##  -1.115712
```

Overall, the individual and dual factor enrichment demonstrates that
Pho is slightly enriched at E(z) peaks that contribute to domains, but
the surprise here is that Zelda shows the strongest of enrichments at
H3K27me3 domains.

In the reporting of Table 1, we include only individual and pair-wise
comparisons that yield a significant fisher’s p-value. Cg+Zld, Pho+GAF,
and Pho+Cg do not yield a significant p-value.

#### Chromatin Accessibility over prospective PREs. (Table 2)

To estimate when an E(z) peak likely first acquires chromatin
accessibility, we take a set of ATAC-seq measurements spanning NC11
through NC13 and estimate when an E(z) peak first gains accessibility.
To do this, we determine the 95th percentile ATAC score for each E(z)
peak at each timepoint (rather than averaging or taking the maximum
value) and we perform clustering. The clusters identify ‘time groups’
that correlate with timing of chromatin accessibility.

```
# above, we loaded the atac data (bigwigs) as a GRangesList. Here we make a counts table.

tab = data.frame(
  NC11 = score(atac[[1]]),
  NC12 = score(atac[[2]]),
  NC13 = score(atac[[3]]),
  overEz = rep(NA, length(atac[[1]]))
)

overEz = findOverlaps(atac[[1]], ez[ez$domain])
tab$overEz[queryHits(overEz)] = subjectHits(overEz)

# now we group by peak and report the 95th percentile score at each timepoint to a table:

zez = tab %>% 
  filter(!is.na(overEz)) %>% 
  group_by(overEz) %>% 
  summarize(z11 = quantile(NC11, 0.95), z12 = quantile(NC12, 0.95), z13 = quantile(NC13, 0.95))

# calculate the distance metric, dropping the column with the peak ID:

dd = dist(zez[,-1])

# perform clustering, using "ward.D" method

set.seed(123)
h = hclust(dd, method = "ward.D")

# we are looking for four categories, so we use cutree to make four groups

k4 = cutree(h, k = 4)

# now we add the cluster identities to the table of peaks:

zez$clust = as.factor(k4)

# tidy the levels

levels(zez$clust) = c("nc12", "nc13", "nc14", "nc11")

zez$clust = relevel(zez$clust, ref = "nc11")

# view membership:

table(zez$clust)
```

```
## 
## nc11 nc12 nc13 nc14 
##  134  201  333  596
```

```
table(zez$clust)/nrow(zez)
```

```
## 
##      nc11      nc12      nc13      nc14 
## 0.1060127 0.1590190 0.2634494 0.4715190
```

We can visualize the changes in accessibility by timepoint and by
group with a simple graph:

```
zez %>% 
  group_by(clust) %>% 
  summarize(k11 = mean(z11), k12 = mean(z12), k13 = mean(z13)) %>% 
  pivot_longer(cols = c(k11, k12, k13), values_to = "score", names_to = 'stage') %>% 
  ggplot(aes(x = stage, y = score, color = clust, group = clust)) +
  geom_line(linewidth = 2) +
  geom_point(size = 5) +
  scale_color_manual(values = rainbow(length(unique(k4))))
```

We can now transfer these clustering information to the
`domains` object.

```
pPRE = as.data.frame(ez[ez$domain])
pPRE$clust = as.factor(zez$clust)

pPRE = GRanges(pPRE)

domover = findOverlaps(pPRE, domains)

pPRE$domainI[queryHits(domover)] = subjectHits(domover)

# pPRE
```

To calculate on a per-domain basis the earliest accessible E(z)
region:

```
as.data.frame(pPRE) %>% 
  dplyr::select(clust, domainI) %>% 
  group_by(domainI) %>% 
  mutate(earliest = case_when(
    any(clust == 'nc11') ~ "nc11",
    any(clust == 'nc12') ~ "nc12",
    any(clust == 'nc13') ~ "nc13", 
    any(clust == 'nc14') ~ "nc14"
  )) %>% 
  dplyr::summarize(e = unique(earliest)) %>%
  mutate(total = dplyr::n()) %>% 
  ungroup() %>% 
  group_by(e) %>% 
  summarize(n = dplyr::n(), pct = dplyr::n()/unique(total))
```

```
## # A tibble: 4 × 3
##   e         n   pct
##   <chr> <int> <dbl>
## 1 nc11     86 0.363
## 2 nc12     70 0.295
## 3 nc13     55 0.232
## 4 nc14     26 0.110
```

This indicates that at the domain-level, there appears to be an
enrichment of domains having at least one early-opening pPRE. This is
currently reported in the manuscript as a table. The barplot is sort of
a striking visual indicator of the magnitude of this effect.

```
# I can't be bothered to code this from the raw tables:

x = rbind(c(10.6, 15.9, 26.3, 47.1), c(36.3, 29.5, 23.2, 10.9))
colnames(x) = paste0("nc", c("11", "12", "13", "14"))
barplot(x, beside = TRUE, col = c("cornflowerblue", "darkblue"), las = 1, ylab = "percent newly open", main = "timing of pPRE accessibility", ylim = c(0,50))
legend("topleft", legend = c("individual pPRE", "earliest in domain"), fill = c("cornflowerblue", "darkblue"))
```

We can formally demonstrate the statistical significance of this
observation using a chi-squared test for given probabilities:

```
chisq.test(x = c(86, 70, 55, 26), p = c(134/1264, 201/1264, 333/1264, 596/1264))$p.value
```

```
## [1] 3.727598e-52
```

```
chisq.test(x = c(86, 70, 55, 26), p = c(134/1264, 201/1264, 333/1264, 596/1264))$method
```

```
## [1] "Chi-squared test for given probabilities"
```

### GAF and Zelda

#### Method Description

There are two main experiments where DESeq2 is used to perform a
comparison. Experiment 1 measures changes in H3K27me3 following
knockdown of GAGA-Factor (GAF). Experiment 2 measures changes in
H3K27me3 in zelda mutants.

For both of these analyses, we rely on counting ChIP signal from each
experimental condition/replicate over 2-kb bins covering the whole
genome. These bins are annotated with membership/overlap with genomic
annotations, including ChIP-seq peaks for transcription factors,
transcriptional start sites, membership in a Polycomb Group (PcG)
Domain, et cetera. The bins are defined using GenomicRanges::tileGenome
and these are filtered to remove low-count bins. Because the bins are
filtered for low-count regions, certain features that are present in the
whole genome annotations may be missing in the final DESeq dataset.
Principal component analysis (PCA) is next performed to evaluate for the
presence of batch effects in the data. In this implementation, a ‘batch
effect’ is defined as a non-biological or unknown contribution to a
significant fraction of the sample variance, as determined by PCA.
Principal components capturing a significant unexplained source of
variance are added to the sample metadata for inclusion in the DESeq
analysis model. The method for model fitting used by DESeq is optimized
by estimating dispersions using the three model fit types (parametric,
local, or mean), calculating mean absolute residual values for the
fittings. The fit method with the smallest mean absolute residual value
is then chosen for DESeq analysis. In this study, the “local” fitting
method provided slightly lower mean absolute residual values than the
“parametric” method. DESeq2 is then performed on the filtered counts
table, using a model of ~ batch + genotype if a batch effect is
observed, or simply ~ genotype if no batch effect is observed, and a
Wald test is performed to identify bins with differential enrichment
between genotypes.

The DESeq results are recovered for the by-genotype comparison
(mutant over wild-type). These raw DESeq results identify 2-kb bins that
can be scored as significant by typical methods, i.e., thresholding for
an absolute log2 fold-change (LFC) greater than one, and an adjusted
p-value less than 0.05. However, because changes in H3K27me3 signal can
potentially span a sequential series or run of 2-kb bins, it is
convenient to identify such runs of adjacent bins passing a significance
test threshold. To identify runs of bins, the DESeq results table was
filtered for bins with a adjusted p-value less than 0.05 and converted
to a GenomicRanges object. Note, the magnitude-of-effect estimator, LFC,
was not used as a thresholding parameter for identifying runs of
significant bins. Adjacent bins were merged through an implementation of
dialysis, reduction, and erosion: filtered bin starts and ends were
extended outward by one bp to create one bp overlaps between adjacent
bins. Overlapping bins were merged using the GenomicRanges::reduce
function, and then one bp was subtracted from the ends of the resulting
table. DESeq measurements (baseMean, LFC, and adjusted p-value) were
propagated to the merged bins as follows. 2-kb bins overlapping the runs
of bins were identified, and baseMean and LFC values for the overlapping
2-kb bins were averaged and assigned to the run. To propagate the
adjusted p-value measurements, we used Fisher’s method, calculating the
chi-squared test statistic by summing the natural logarithms of the
adjusted p-values and multiplying by negative two.

Significant 2-kb bins or runs of bins were scored for membership in
PcG domains by measuring any overlap between a bin or run and an
annotated PcG Domain.

#### Implementation

```
# make a set of 2k bins to use for DESeq analysis

bins2k = namer(cleanup(tileGenome(seqlengths(Dmelanogaster), tilewidth = 2000, cut.last.tile.in.chrom = TRUE)))

# find which bins overlap with PcG domains

bod = findOverlaps(bins2k, domains)

# associate bins with additional peak sets of interest.

bins2k$withZld = as.logical(countOverlaps(bins2k, zld))
bins2k$withGAF = as.logical(countOverlaps(bins2k, gaf))
bins2k$withPho = as.logical(countOverlaps(bins2k, pho))
bins2k$withEz = as.logical(countOverlaps(bins2k, ez))
bins2k$withTSS = as.logical(countOverlaps(bins2k, zygtss))

# assign overlapping zygotic TSS IDs and PcG domains with bins.

bot = findOverlaps(bins2k, zygtss)
concordance = tapply(zygtss$fb_symbol[subjectHits(bot)], queryHits(bot), function(x){
    paste0(x, collapse = ", ")
})
bins2k$tssName = rep(NA, length(bins2k))
bins2k$tssName[as.numeric(names(concordance))] = concordance

bins2k$inDomain = rep(NA, length(bins2k))
bins2k$inDomain[queryHits(bod)] = names(domains)[subjectHits(bod)]

head(bins2k)
```

```
## GRanges object with 6 ranges and 7 metadata columns:
##                     seqnames      ranges strand |   withZld   withGAF   withPho
##                        <Rle>   <IRanges>  <Rle> | <logical> <logical> <logical>
##        chr2L:1-2000    chr2L      1-2000      * |     FALSE     FALSE     FALSE
##     chr2L:2001-4000    chr2L   2001-4000      * |     FALSE     FALSE     FALSE
##     chr2L:4001-6000    chr2L   4001-6000      * |      TRUE      TRUE     FALSE
##     chr2L:6001-8000    chr2L   6001-8000      * |     FALSE      TRUE     FALSE
##    chr2L:8001-10000    chr2L  8001-10000      * |     FALSE     FALSE     FALSE
##   chr2L:10001-12000    chr2L 10001-12000      * |     FALSE     FALSE     FALSE
##                        withEz   withTSS     tssName              inDomain
##                     <logical> <logical> <character>           <character>
##        chr2L:1-2000     FALSE     FALSE        <NA>                  <NA>
##     chr2L:2001-4000     FALSE     FALSE        <NA>                  <NA>
##     chr2L:4001-6000      TRUE     FALSE        <NA> PcGD:chr2L:5000-10000
##     chr2L:6001-8000      TRUE     FALSE        <NA> PcGD:chr2L:5000-10000
##    chr2L:8001-10000     FALSE     FALSE        <NA> PcGD:chr2L:5000-10000
##   chr2L:10001-12000     FALSE     FALSE        <NA>                  <NA>
##   -------
##   seqinfo: 6 sequences from dm6 genome
```

With these bins and count tables, we can now perform an analysis for
both the GAF and the Zelda experiments.

##### Filtering Low Count Bins

For each experiment, we will calculate the sum of reads in each bin
and filter out bins with low reads. To do this, we calculate the rowSums
for bins in PcG domains and bins outside of PcG Domains, log10
transform, and plot a histogram. This should allow us to eyeball a
cutoff where we just nick off the lower tail of the bins within
domains.

Rationale: the signal should be concentrated within Domains.
Therefore, we use the distribution of bins within domains as our guide
for thresholding.

###### GAF

```
hist(log10(rowSums(gafcts[is.na(bins2k$inDomain),] + 1)), breaks = 1000, main = "GAF: distribution of 2k bin row sums H3K27me3", freq = TRUE, xlim = c(0, 5), xlab = "log10(summed bin counts + 1)")
hist(log10(rowSums(gafcts[!is.na(bins2k$inDomain),] + 1)), breaks = 200, add = TRUE, col = 'red', border = 'red', freq = TRUE)
abline(v = 3.12, col = 'magenta', lty = 'dashed')
```

```
gafthresh = 10^3.12
```

The eyeballed threshold is 1e+3.12, so we apply this to our counts
table and filter out low-count bins.

```
gafcts = gafcts[rowSums(gafcts) > gafthresh,]
```

###### zld

For the Zelda comparisons, we are interested in H3K27me3 as well as
other modifications/ChIP results.

###### zld H3K27me3

First, H3K27me3

```
hist(log10(rowSums(zldcts[is.na(bins2k$inDomain), k27I] + 1)), breaks = 1000, main = "Zld: distribution of 2k bin row sums K27me3", xlim = c(0,5))
hist(log10(rowSums(zldcts[!is.na(bins2k$inDomain),k27I] + 1)), breaks = 200, col = 'red', border = 'red', add = TRUE)
abline(v = 3.33, col = 'magenta', lty = 'dashed')
```

```
zldthresh = 10^3.33
```

Here we show the distribution for the K27me3 data, but below, we
filter the zelda counts table by the set of bins that are in common to
the K27me3, H2Aub, and E(z) counts tables at a threshold of 1e+3.33.

```
zldorig = zldcts

keepers = apply(
    cbind(bad1 = rowSums(zldcts[,k27I]) > zldthresh,
          bad2 = rowSums(zldcts[,h2aI]) > zldthresh,
          bad3 = rowSums(zldcts[,ezI]) > zldthresh
    ), 1, all)

zld2a = zldcts[keepers, h2aI]
zldcts = zldcts[keepers, k27I]
```

###### zld Pol2

We use the zld Pol2 ChIP experiment mainly to explore the
distribution of zelda-dependent zygotic TSS across the PcG domains. It
is less important that this counts table corresponds exactly with the
modifications above.

```
hist(log10(rowSums(zldorig[is.na(bins2k$inDomain), polI] + 1)), breaks = 1000, main = "Zld: distribution of 2k bin row sums RNA Pol2", xlim = c(0,5))
hist(log10(rowSums(zldorig[!is.na(bins2k$inDomain), polI] + 1)), breaks = 200, col = 'red', border = 'red', add = TRUE)
abline(v = 2.7, col = 'magenta', lty = 'dashed')
```

```
zldPthresh = 10^2.7
```

```
zldpol = zldorig[, polI]
zldpol = zldpol[rowSums(zldpol) > zldPthresh,]
```

###### zld H2Aub

We do not do much with the H2Aub data in this manuscript but it is
useful to compare between H3K27me3 and H2Aub to ask whether there are
correlations in effects at sites that have reduced/increased
modifications following loss of Zelda.

```
hist(log10(rowSums(zldorig[is.na(bins2k$inDomain), h2aI] + 1)), breaks = 1000, main = "Zld: distribution of 2k bin row sums H2Aub", xlim = c(0,5))
hist(log10(rowSums(zldorig[!is.na(bins2k$inDomain), h2aI] + 1)), breaks = 200, col = 'red', border = 'red', add = TRUE)
abline(v = 3.33, col = 'magenta', lty = 'dashed')
```

The distributions are close between the H3K27me3 and H2Aub datasets,
so to ensure maximal overlap between the analyses, and because we just
want to compare this data to the H3K27me3 data, we will count H2Aub over
the same set of bins available for the H3K27me3 analysis. This object
`zld2a` was defined above containing the H2Aub datasets.

We are now set to evaluate batch effects.

##### Evaluating batch effects

To do this, we initiate the DESeq object for each analysis and first
perform a PCA.

###### GAF

```
coldataG = data.frame(sample = colnames(gafcts), 
                       genotype = factor(rep(c("wt", "gaf"), 3),levels = c('wt','gaf'))
                       )

rownames(coldataG) = coldataG$sample

ddsG = DESeqDataSetFromMatrix(countData = gafcts, colData = coldataG, design = ~ genotype)

rld= rlog(ddsG, blind=FALSE, fitType = "local")

pca= plotPCA(rld, intgroup = c("genotype"), returnData = TRUE)
percentVar= round(100* attr(pca, "percentVar"))

ggplot(pca, aes(PC1,PC2, color= genotype))+
  geom_point(size=4, alpha=0.7)+
  xlab(paste0("PC1: ", percentVar[1], "% variance")) +
  ylab(paste0("PC2: ", percentVar[2], "% variance")) +
  scale_color_manual(values= c("azure3", "cornflowerblue"))+ theme(axis.line = element_line(color = "black"))+
  theme_bw()+
  coord_equal() +
   xlim(-10, 10) +
   ylim(-10, 10) +
  labs(title = "PCA 2k bins, GAF")
```

Here we see that the first principal component identifies some kind
of batch effect, and that the second PC picks up genotype-specific
differences. Normally this would be a concern, but as we see in the
reported results, the GAF KD does not have a strong effect on the
K27me3.

We can account for this batch effect by appending the numeric value
of PC1 to the coldataG object and re-specifying the DESeq object with
the updated design.

```
coldataG$pc1 = pca$PC1

ddsG = DESeqDataSetFromMatrix(countData = gafcts, colData = coldataG, design = ~ pc1 + genotype)
```

###### zld

##### H3K27me3

We repeat the same approach as above for the Zld K27me3 samples.
Note, we over-write variables below.

```
coldataZ = data.frame(sample = colnames(zldcts), 
                       genotype = factor(rep(c("wt", "zld"), each = 4),levels = c('wt','zld'))
                       )

rownames(coldataZ) = coldataZ$sample

ddsZ = DESeqDataSetFromMatrix(countData = zldcts, colData = coldataZ, design = ~ genotype)

rld= rlog(ddsZ, blind=FALSE, fitType = "local")

pca= plotPCA(rld, intgroup = c("genotype"), returnData = TRUE)
percentVar= round(100* attr(pca, "percentVar"))

ggplot(pca, aes(PC1,PC2, color= genotype))+
  geom_point(size=4, alpha=0.7)+
  xlab(paste0("PC1: ", percentVar[1], "% variance")) +
  ylab(paste0("PC2: ", percentVar[2], "% variance")) +
  scale_color_manual(values= c("azure3", "cornflowerblue"))+ theme(axis.line = element_line(color = "black"))+
  theme_bw()+
  coord_equal() +
   xlim(-10, 10) +
   ylim(-10, 10) +
  labs(title = "PCA 2k bins, zld")
```

This is better, and reflects a stronger influence of Zld on the
K27me3 deposition, as we will soon see. Nevertheless, PC2 accounts for
13% of the variance, and likely represents some kind of a batch effect
in these data. We will account for it as we did above.

```
coldataZ$pc2 = pca$PC2

ddsZ = DESeqDataSetFromMatrix(countData = zldcts, colData = coldataZ, design = ~ pc2 + genotype)
```

###### RNA Pol2

```
coldataZp = data.frame(sample = colnames(zldpol), 
                       genotype = factor(rep(c("wt", "zld"), each = 2),levels = c('wt','zld'))
                       )

rownames(coldataZp) = coldataZp$sample

ddsZp = DESeqDataSetFromMatrix(countData = zldpol, colData = coldataZp, design = ~ genotype)

rld= rlog(ddsZp, blind=FALSE, fitType = "local")

pca= plotPCA(rld, intgroup = c("genotype"), returnData = TRUE)
percentVar= round(100* attr(pca, "percentVar"))

ggplot(pca, aes(PC1,PC2, color= genotype))+
  geom_point(size=4, alpha=0.7)+
  xlab(paste0("PC1: ", percentVar[1], "% variance")) +
  ylab(paste0("PC2: ", percentVar[2], "% variance")) +
  scale_color_manual(values= c("azure3", "cornflowerblue"))+ theme(axis.line = element_line(color = "black"))+
  theme_bw()+
  coord_equal() +
   xlim(-23, 23) +
   ylim(-23, 23) +
  labs(title = "PCA 2k bins, zld, Pol2")
```

There does not seem to be a significant batch effect for the Pol2
data.

###### H2Aub

```
coldataZ2a = data.frame(sample = colnames(zld2a), 
                       genotype = factor(rep(c("wt", "zld"), each = 2),levels = c('wt','zld'))
                       )

rownames(coldataZ2a) = coldataZ2a$sample

ddsZ2a = DESeqDataSetFromMatrix(countData = zld2a, colData = coldataZ2a, design = ~ genotype)

rld= rlog(ddsZ2a, blind=FALSE, fitType = "local")

pca= plotPCA(rld, intgroup = c("genotype"), returnData = TRUE)
percentVar= round(100* attr(pca, "percentVar"))

ggplot(pca, aes(PC1,PC2, color= genotype))+
  geom_point(size=4, alpha=0.7)+
  xlab(paste0("PC1: ", percentVar[1], "% variance")) +
  ylab(paste0("PC2: ", percentVar[2], "% variance")) +
  scale_color_manual(values= c("azure3", "cornflowerblue"))+ theme(axis.line = element_line(color = "black"))+
  theme_bw()+
  coord_equal() +
  xlim(-15, 15) +
  ylim(-15, 15) +
  labs(title = "PCA 2k bins, zld")
```

There does not appear to be a substantial batch effect for the H2Aub
data. We now need to evaluate the optimal fit for the DESeq model.

##### Estimating an optimal DESeq fit type

We want to estimate dispersions using the three available fit types
and determine the minimal mean absolute residual derived from these
fittings. In practice, the ‘parametric’ model sometimes does not fit
well, and it defaults to “local”.

###### GAF

```
ddsG = estimateSizeFactors(ddsG)
parG = estimateDispersions(ddsG, fitType = "parametric", maxit = 500, quiet = TRUE)
locG = estimateDispersions(ddsG, fitType = "local", quiet = TRUE)
meaG = estimateDispersions(ddsG, fitType = "mean", quiet = TRUE)
```

Now we calculate the mean absolute residual and choose the model with
the minimum value.

```
c(parametric = median(abs(mcols(parG)$dispGeneEst - mcols(parG)$dispFit)),
  local = median(abs(mcols(locG)$dispGeneEst - mcols(locG)$dispFit)), 
  mean = median(abs(mcols(meaG)$dispGeneEst - mcols(meaG)$dispFit)))
```

```
##  parametric       local        mean 
## 0.009892165 0.008413572 0.009882398
```

In this case the local fit minimizes the mean absolute residual.
These differences don’t seem huge though…

```
plotDispEsts(locG, main = "GAF Local Fit Dispersion Estimates")
```

###### zld

```
ddsZ = estimateSizeFactors(ddsZ)
parZ = estimateDispersions(ddsZ, fitType = "parametric", maxit = 500, quiet = TRUE)
```

```
## -- note: fitType='parametric', but the dispersion trend was not well captured by the
##    function: y = a/x + b, and a local regression fit was automatically substituted.
##    specify fitType='local' or 'mean' to avoid this message next time.
```

```
locZ = estimateDispersions(ddsZ, fitType = "local", quiet = TRUE)
meaZ = estimateDispersions(ddsZ, fitType = "mean", quiet = TRUE)
```

The warning tells us that parametric isn’t going to cut it, so in
this case we’d probably only be considering either local or mean
fitting.

```
c(parametric = median(abs(mcols(parZ)$dispGeneEst - mcols(parZ)$dispFit)),
  local = median(abs(mcols(locZ)$dispGeneEst - mcols(locZ)$dispFit)), 
  mean = median(abs(mcols(meaZ)$dispGeneEst - mcols(meaZ)$dispFit)))
```

```
##  parametric       local        mean 
## 0.005200454 0.005229282 0.009058573
```

And again local fitting has the lowest mean absolute residuals.

```
plotDispEsts(locZ, main = "Zelda Local Fit Dispersion Estimates")
```

We will do a local fit type for the zld Pol2 data.

Now we are ready to do DESeq2.

##### DESeq2

Above, we determined that both experiments should use the local fit
type for the DESeq step. We will perform the DESeq using a Wald test and
a local fit. We will calculate a results table and apply a
log2(FoldChange) shrink to minimize effects from low mean count regions
on the Log2(FC) measurement. Then we will associate the results table
with the annotations in our 2-kb bins. To do this, we will need a
minimized data.frame of the bin metadata, plus the bin name
identifiers:

```
bindf = as.data.frame(bins2k)
bindf = bindf %>% 
    rownames_to_column(var = "bin_name") %>% 
    select(-seqnames, -start, -end, -width, -strand)
```

###### GAF

```
ddsG = DESeq(ddsG, test = "Wald", fitType = "local", quiet = TRUE)

resG = results(ddsG, contrast = c("genotype", "gaf", "wt"), independentFiltering = FALSE)
resG = lfcShrink(dds = ddsG, res = resG, type = 'apeglm', coef = 3, quiet = TRUE)

resG = as.data.frame(resG)

resG= resG %>%
    mutate(
        baseMean = round(baseMean, 2), 
        lfc = round(log2FoldChange, 4)
        ) %>% 
    rownames_to_column(var = "bin_name") %>% 
    select(-lfcSE, -pvalue, -log2FoldChange)

resG = left_join(resG, bindf)
```

```
## Joining, by = "bin_name"
```

###### zld

##### H3K27me3

```
ddsZ = DESeq(ddsZ, test = "Wald", fitType = "local", quiet = TRUE)
resZ = results(ddsZ, contrast = c("genotype", "zld", "wt"), independentFiltering = FALSE)
resZ = lfcShrink(dds = ddsZ, res = resZ, type = 'apeglm', coef = 3, quiet = TRUE)

resZ = as.data.frame(resZ)

resZ= resZ %>%
    mutate(
        baseMean = round(baseMean, 2), 
        lfc = round(log2FoldChange, 4)
        ) %>% 
    rownames_to_column(var = "bin_name") %>% 
    select(-lfcSE, -pvalue, -log2FoldChange)

resZ = left_join(resZ, bindf)
```

```
## Joining, by = "bin_name"
```

###### Pol2

```
ddsZp = DESeq(ddsZp, test = "Wald", fitType = "local", quiet = TRUE)
resZp = results(ddsZp, contrast = c("genotype", "zld", "wt"), independentFiltering = FALSE)
resZp = lfcShrink(dds = ddsZp, res = resZp, type = 'apeglm', coef = 2, quiet = TRUE)

resZp = as.data.frame(resZp)

resZp= resZp %>%
    mutate(
        baseMean = round(baseMean, 2), 
        lfc = round(log2FoldChange, 4)
        ) %>% 
    rownames_to_column(var = "bin_name") %>% 
    select(-lfcSE, -pvalue, -log2FoldChange)

resZp = left_join(resZp, bindf)
```

```
## Joining, by = "bin_name"
```

###### H2Aub

```
ddsZ2a = DESeq(ddsZ2a, test = "Wald", fitType = "local", quiet = TRUE)
resZ2a = results(ddsZ2a, contrast = c("genotype", "zld", "wt"), independentFiltering = FALSE)
resZ2a = lfcShrink(dds = ddsZ2a, res = resZ2a, type = 'apeglm', coef = 2, quiet = TRUE)

resZ2a = as.data.frame(resZ2a)

resZ2a= resZ2a %>%
    mutate(
        baseMean = round(baseMean, 2), 
        lfc = round(log2FoldChange, 4)
        ) %>% 
    rownames_to_column(var = "bin_name") %>% 
    select(-lfcSE, -pvalue, -log2FoldChange)

resZ2a = left_join(resZ2a, bindf)
```

```
## Joining, by = "bin_name"
```

##### Generating “Runs” of Bins

We have bins above with adjusted p-values less than 0.05, and they
may be adjacent to other bins with a similarly small adjusted p-value.
We want now to associate these adjacent bins into continuous ‘runs’ of
bins, and to reassociate with the runs the genomic features that any of
the constituent bins associate with.

We have below a general function that will perform this run-making
operation, given a set of DESeq results and a set of original starting
bins.

```
runner = function(res, bins){

    b = left_join(
        as.data.frame(bins) %>% 
            rownames_to_column(var = "bin_name"),
        res %>% 
            select(bin_name, baseMean, padj, lfc)
        , by = 'bin_name') %>% 
        mutate(
        padj = case_when(
            is.na(padj) ~ 1,
            !is.na(padj) ~ padj
        ),
        baseMean = case_when(
            is.na(baseMean) ~ 0,
            !is.na(baseMean) ~ baseMean
        ),
        lfc = case_when(
            is.na(lfc) ~ 0,
            !is.na(lfc) ~ lfc
        )
        ) %>% 
        GRanges()
    
    runs = b[b$padj <= 0.05]
    
    start(runs) = start(runs) - 1
    end(runs) = end(runs) + 1
    
    runs = GenomicRanges::reduce(runs)
    
    start(runs) = start(runs) + 1
    end(runs) = end(runs) - 1
    
    
    overruns = findOverlaps(query = b, subject = runs)

    mcols(runs) = data.frame(
        meanBaseMean = rep(NA, length(runs)),
        meanLFC = rep(NA, length(runs)) ,
        fishersp = rep(NA, length(runs))
    )

    query_idx <- queryHits(overruns)
    subject_idx <- subjectHits(overruns)

    grouped_data <- split(query_idx, subject_idx)

    runs$meanBaseMean <- sapply(grouped_data, function(idx) mean(b$baseMean[idx], na.rm = TRUE))
    runs$meanLFC <- sapply(grouped_data, function(idx) mean(b$lfc[idx], na.rm = TRUE))

    runs$fishersp <- sapply(grouped_data, function(idx) {
        pchisq(q = (-2 * sum(log(b$padj[idx]))), 
            df = 2 * length(idx), 
            lower.tail = FALSE)
    })

    runs$ranker = rank(abs(runs$meanLFC) * log10(runs$meanBaseMean))
    
    runs$inDomain = rep(NA, length(runs))
    overd = findOverlaps(runs, domains) # reference to outside data (domains)
    runs$inDomain[queryHits(overd)] = names(domains)[subjectHits(overd)]
    runs$withZld = as.logical(countOverlaps(runs, zld)) # reference to outside data (zld)
    runs$withGAF = as.logical(countOverlaps(runs, gaf)) # reference to outside data (gaf)
    runs$withPho = as.logical(countOverlaps(runs, pho)) # reference to outside data (pho)
    runs$withEz = as.logical(countOverlaps(runs, ez)) # reference to outside data (ez)
    
    runs$withTSS = as.character(rep(NA, length(runs)))
    overt = findOverlaps(runs, zygtss) # reference to outside data (zygtss)
    concattss = tapply(zygtss$fb_symbol[subjectHits(overt)], queryHits(overt), function(x) paste0(x, collapse = ", "))
    
    runs$withTSS[as.numeric(names(concattss))] = concattss 
    runs = namer(runs) # reference to outside data (namer)
    
    return(as.data.frame(runs))
}
```

###### GAF

```
runsG = runner(resG, bins2k)
```

###### zld

```
runsZ = runner(resZ, bins2k)
```

We have above calculated runs of bins with padj < 0.05, and
propagated the mean “baseMean”, “log2FoldChange” and calculated a
chi-square p-value using Fisher’s method for combining p-values. We have
associated the runs with domains, peaks of interest, as well as the
names of zygotic TSS regions that occur within the run.

##### Analysis

We can now visualize the distributions of calculated differential
enrichment. We favor looking at the effects that happen within domains,
since this is where we expect changes to be occurring, but we can also
evaluate the extent of changes outside of domains as well.

###### GAF

First, by bin:

```
resG %>% 
    filter(!is.na(inDomain)) %>% 
    mutate(sig = padj < 0.05 & abs(lfc) > 1) %>% 
    mutate(tn = case_when(
        withTSS & sig ~ tssName,
        !(withTSS & sig) ~ NA_character_
    )) %>% 
    ggplot(aes(x = lfc, y = -log10(padj), col = sig, size = baseMean, label = tn)) +
    geom_point(alpha = 0.8) + 
    scale_color_manual(values = c("black","cornflowerblue")) +
    xlim(-4.5, 4.5) +
    ylim(0, 40) +
    coord_equal(ratio = (9/40)) +
    geom_hline(yintercept = -log10(0.05), col = 'blue', linetype = 'dashed') +
    geom_vline(xintercept = c(-1, 1), col = 'blue', linetype = 'dashed') +
    scale_size_area() +
    ggrepel::geom_label_repel(show.legend = FALSE, 
                             box.padding = 0.8, 
                             max.overlaps = 1e+5, 
                             size = 2, 
                             color = "brown3",
                             na.rm = TRUE
                             )
```

and by run, first with all tss labels…

```
runsG %>% 
    filter(!is.na(inDomain)) %>% 
    mutate(sig = fishersp < 0.05 & abs(meanLFC) > 1) %>% 
    ggplot(aes(x = meanLFC, 
               y = -log10(fishersp), 
               col = sig, 
               size = meanBaseMean, 
               label = withTSS)
           ) +
    geom_point(alpha = 0.8) + 
    scale_color_manual(values = c("black","cornflowerblue")) +
    xlim(-4.5, 4.5) +
    ylim(0, 40) +
    coord_equal(ratio = (9/40)) +
    geom_hline(yintercept = -log10(0.05), col = 'blue', linetype = 'dashed') +
    geom_vline(xintercept = c(-1, 1), col = 'blue', linetype = 'dashed') +
    scale_size_area() +
    ggrepel::geom_label_repel(show.legend = FALSE, 
                             box.padding = 0.8, 
                             max.overlaps = 1e+5, 
                             size = 2, 
                             color = "brown3",
                             na.rm = TRUE
                             )
```

and next, by filtering the tss of interest:

```
favorites = c("esg", "sna", "en", "inv", "sisA")

runsG %>% 
    filter(!is.na(inDomain)) %>% 
    mutate(sig = fishersp < 0.05 & abs(meanLFC) > 1) %>% 
    mutate(wT = case_when(
        withTSS %in% favorites ~ withTSS,
        !withTSS %in% favorites ~ NA_character_
    )) %>%
    ggplot(aes(x = meanLFC, 
               y = -log10(fishersp), 
               col = sig, 
               size = meanBaseMean, 
               label = wT)
           ) +
    geom_point(alpha = 0.8) + 
    scale_color_manual(values = c("black","cornflowerblue")) +
    xlim(-4.5, 4.5) +
    ylim(0, 40) +
    coord_equal(ratio = (9/40)) +
    geom_hline(yintercept = -log10(0.05), col = 'blue', linetype = 'dashed') +
    geom_vline(xintercept = c(-1, 1), col = 'blue', linetype = 'dashed') +
    scale_size_area() +
    ggrepel::geom_label_repel(show.legend = FALSE, 
                             box.padding = 0.8, 
                             max.overlaps = 1e+5, 
                             size = 2, 
                             color = "brown3"
                             )
```

```
## Warning: Removed 229 rows containing missing values or values outside the scale range
## (`geom_label_repel()`).
```

Are there bin-wise or run-wise changes outside of domains?

```
resG %>% 
    filter(is.na(inDomain)) %>% 
    mutate(sig = padj < 0.05 & abs(lfc) > 1) %>% 
    mutate(tn = case_when(
        withTSS & sig ~ tssName,
        !(withTSS & sig) ~ NA_character_
    )) %>% 
    ggplot(aes(x = lfc, y = -log10(padj), col = sig, size = baseMean, label = tn)) +
    geom_point(alpha = 0.8) + 
    scale_color_manual(values = c("black","cornflowerblue")) +
    xlim(-4.5, 4.5) +
    ylim(0, 40) +
    coord_equal(ratio = (9/40)) +
    geom_hline(yintercept = -log10(0.05), col = 'blue', linetype = 'dashed') +
    geom_vline(xintercept = c(-1, 1), col = 'blue', linetype = 'dashed') +
    scale_size_area() +
    ggrepel::geom_label_repel(show.legend = FALSE, 
                             box.padding = 0.8, 
                             max.overlaps = 1e+5, 
                             size = 2, 
                             color = "brown3",
                             na.rm = TRUE
                             )
```

By run, outside of domains:

```
runsG %>% 
    filter(is.na(inDomain)) %>% 
    mutate(sig = fishersp < 0.05 & abs(meanLFC) > 1) %>% 
    mutate(wT = case_when(
        !is.na(withTSS) & sig ~ withTSS,
        !(!is.na(withTSS) & sig) ~ NA_character_
    )) %>% 
    ggplot(aes(x = meanLFC, 
               y = -log10(fishersp), 
               col = sig, 
               size = meanBaseMean, 
               label = wT)
           ) +
    geom_point(alpha = 0.8) + 
    scale_color_manual(values = c("black","cornflowerblue")) +
    xlim(-4.5, 4.5) +
    ylim(0, 40) +
    coord_equal(ratio = (9/40)) +
    geom_hline(yintercept = -log10(0.05), col = 'blue', linetype = 'dashed') +
    geom_vline(xintercept = c(-1, 1), col = 'blue', linetype = 'dashed') +
    scale_size_area() +
    ggrepel::geom_label_repel(show.legend = FALSE, 
                             box.padding = 0.8, 
                             max.overlaps = 1e+5, 
                             size = 2, 
                             color = "brown3",
                             na.rm = TRUE
                             )
```

###### zld

First, by bin:

```
resZ %>% 
    filter(!is.na(inDomain)) %>% 
    mutate(sig = padj < 0.05 & abs(lfc) > 1) %>% 
    mutate(tn = case_when(
        withTSS & sig ~ tssName,
        !(withTSS & sig) ~ NA_character_
    )) %>% 
    ggplot(aes(x = lfc, y = -log10(padj), col = sig, size = baseMean, label = tn)) +
    geom_point(alpha = 0.8) + 
    scale_color_manual(values = c("black","cornflowerblue")) +
    xlim(-4.5, 4.5) +
    ylim(0, 70) +
    coord_equal(ratio = (9/70)) +
    geom_hline(yintercept = -log10(0.05), col = 'blue', linetype = 'dashed') +
    geom_vline(xintercept = c(-1, 1), col = 'blue', linetype = 'dashed') +
    scale_size_area() +
    ggrepel::geom_label_repel(show.legend = FALSE, 
                             box.padding = 0.8, 
                             max.overlaps = 1e+5, 
                             size = 2, 
                             color = "brown3",
                             na.rm = TRUE
                             )
```

There are quite a few of these. What if we limit it to TSS-containing
bins within domains?

```
resZ %>% 
    filter(!is.na(inDomain) & withTSS) %>% 
    mutate(sig = padj < 0.05 & abs(lfc) > 1) %>% 
    mutate(tn = case_when(
        withTSS & sig ~ tssName,
        !(withTSS & sig) ~ NA_character_
    )) %>% 
    ggplot(aes(x = lfc, y = -log10(padj), col = sig, size = baseMean, label = tn)) +
    geom_point(alpha = 0.8) + 
    scale_color_manual(values = c("black","cornflowerblue")) +
    xlim(-4.5, 4.5) +
    ylim(0, 40) +
    coord_equal(ratio = (9/40)) +
    geom_hline(yintercept = -log10(0.05), col = 'blue', linetype = 'dashed') +
    geom_vline(xintercept = c(-1, 1), col = 'blue', linetype = 'dashed') +
    scale_size_area() +
    ggrepel::geom_label_repel(show.legend = FALSE, 
                             box.padding = 0.2, 
                             max.overlaps = 1e+5, 
                             size = 2, 
                             color = "brown3",
                             na.rm = TRUE,
                             min.segment.length = 0.1
                             )
```

Let’s see a table of how this breaks down:

```
ct = resZ %>% 
    filter(withTSS) %>% 
    mutate(sig = padj < 0.05 & abs(lfc) > 1, domain = !is.na(inDomain)) %>% 
    group_by(sig, domain) %>% 
    summarize(n = dplyr::n()) %>% 
    pivot_wider(values_from = n, names_from = domain, names_prefix = "domain.") %>% 
    ungroup() %>% 
    arrange(desc(sig)) %>% 
    mutate(sig = paste0("sig.", sig)) %>% 
    select(sig, "domain.TRUE", "domain.FALSE") %>% 
    column_to_rownames(var = "sig")
```

```
## `summarise()` has grouped output by 'sig'. You can override using the `.groups`
## argument.
```

```
ct
```

```
##           domain.TRUE domain.FALSE
## sig.TRUE           22            2
## sig.FALSE         133          311
```

Here, we have 155 early zygotic TSS regions within PcG domains (left
column in `ct`). 22 of these have significant changes in
H3K27me3 in zelda mutants. Outside of domains (right column in
`ct`), only 2 zygotic TSS have significant changes in
H3K27me3. There are 133 TSS within domains that do not have significant
changes in H3K27me3 following loss of Zelda.

Back to volcano plots. Now we plot by run, first with all tss
labels…

```
runsZ %>% 
    filter(!is.na(inDomain)) %>% 
    mutate(sig = fishersp < 0.05 & abs(meanLFC) > 1) %>% 
    mutate(wT = case_when(
        !is.na(withTSS) & sig ~ withTSS,
        is.na(withTSS) & sig ~ NA_character_
    )) %>%
    ggplot(aes(x = meanLFC, 
               y = -log10(fishersp), 
               col = sig, 
               size = meanBaseMean, 
               label = wT)
           ) +
    geom_point(alpha = 0.8) + 
    scale_color_manual(values = c("black","cornflowerblue")) +
    xlim(-4.5, 4.5) +
    ylim(0, 250) +
    coord_equal(ratio = (9/250)) +
    geom_hline(yintercept = -log10(0.05), col = 'blue', linetype = 'dashed') +
    geom_vline(xintercept = c(-1, 1), col = 'blue', linetype = 'dashed') +
    scale_size_area() +
    ggrepel::geom_label_repel(show.legend = FALSE, 
                             box.padding = 0.8, 
                             max.overlaps = 1e+5, 
                             size = 2, 
                             color = "brown3"
                             )
```

```
## Warning: Removed 530 rows containing missing values or values outside the scale range
## (`geom_label_repel()`).
```

and next by filtering the tss of interest:

```
favorites = c("esg", "sna", "sisA", "grh", "wg", "eve", "ind", "hbn", "sisA", "amos", "ato, CG9626", "sim", "wntD")

runsZ %>% 
    filter(!is.na(inDomain)) %>% 
    mutate(wT = case_when(
        withTSS %in% favorites ~ withTSS,
        !withTSS %in% favorites ~ NA_character_
    )) %>% 
    mutate(sig = fishersp < 0.05 & abs(meanLFC) > 1) %>% 
    ggplot(aes(x = meanLFC, 
               y = -log10(fishersp), 
               col = sig, 
               size = meanBaseMean, 
               label = wT)
           ) +
    geom_point(alpha = 0.8) + 
    scale_color_manual(values = c("black","cornflowerblue")) +
    xlim(-4.5, 4.5) +
    ylim(0, 250) +
    coord_equal(ratio = (9/250)) +
    geom_hline(yintercept = -log10(0.05), col = 'blue', linetype = 'dashed') +
    geom_vline(xintercept = c(-1, 1), col = 'blue', linetype = 'dashed') +
    scale_size_area() +
    ggrepel::geom_label_repel(show.legend = FALSE, 
                             box.padding = 1, 
                             max.overlaps = 1e+5, 
                             size = 2, 
                             color = "brown3"
                             )
```

```
## Warning: Removed 534 rows containing missing values or values outside the scale range
## (`geom_label_repel()`).
```

###### Zld with Pol2

###### Number of early zygotic TSS with zelda-sensitive RNA Pol2

How many total zygotic TSS survived the filtering for DESeq? There
are 813 in the Chen source list.

```
resZp %>% 
  filter(withTSS) %>% 
  nrow()
```

```
## [1] 798
```

798 make it through the RNA Pol2 DESeq2 analysis. How many have
significantly reduced RNA Pol2 signal in Zelda mutants?

```
resZp %>% 
  filter(withTSS) %>% 
  mutate(sigDown = lfc < -1 & padj < 0.05) %>% 
  group_by(sigDown) %>% 
  summarize(n = dplyr::n())
```

```
## # A tibble: 2 × 2
##   sigDown     n
##   <lgl>   <int>
## 1 FALSE     529
## 2 TRUE      269
```

There are 269 of 798 regions that show a statistically significant
reduction in Pol2 signal following loss of zelda (33.7%)

###### How many zygotic TSS overlap with E(z) peaks?

Overall, how many (significant or not) overlap with E(z) peaks?

```
resZp %>% 
  filter(withTSS) %>% 
  group_by(withEz) %>% 
  summarize(n = dplyr::n())
```

```
## # A tibble: 2 × 2
##   withEz     n
##   <lgl>  <int>
## 1 FALSE    326
## 2 TRUE     472
```

472 of 798 early zygotic TSS overlap with an E(z) peak. (59.1%)

###### How many zygotic TSS reside in PcG domains?

```
resZp %>% 
  filter(withTSS, !is.na(inDomain)) %>% 
  nrow()
```

```
## [1] 155
```

```
ct = resZp %>% 
    filter(withTSS) %>% 
    mutate(sigDown = padj < 0.05 & lfc < -1, domain = !is.na(inDomain)) %>% 
    group_by(sigDown, domain) %>% 
    summarize(n = dplyr::n()) %>% 
    pivot_wider(values_from = n, names_from = domain, names_prefix = "domain.") %>% 
    ungroup() %>% 
    arrange(desc(sigDown)) %>% 
    mutate(sigDown = paste0("sigDown.", sigDown)) %>% 
    select(sigDown, "domain.TRUE", "domain.FALSE") %>% 
    column_to_rownames(var = "sigDown")
```

```
## `summarise()` has grouped output by 'sigDown'. You can override using the
## `.groups` argument.
```

```
ct
```

```
##               domain.TRUE domain.FALSE
## sigDown.TRUE           66          203
## sigDown.FALSE          89          440
```

There are 155 early zygotic TSS within PcG domains, and there are 269
early zygotic TSS that depend on Zelda for Pol2 occupancy. The overlap
between these groups is 66 TSS (42% of the TSS within PcG domains).

```
fisher.test(ct)
```

```
## 
##  Fisher's Exact Test for Count Data
## 
## data:  ct
## p-value = 0.0106
## alternative hypothesis: true odds ratio is not equal to 1
## 95 percent confidence interval:
##  1.101731 2.335345
## sample estimates:
## odds ratio 
##   1.606403
```

There is a weak statistically significant enrichment for
Zelda-dependent Pol2 within PcG domains. But this is not a particularly
strong magnitude of effect (or statistical outcome).

The volcano plot for the Pol2 DESeq looks like this:

```
resZp %>% 
    filter(!is.na(inDomain) & withTSS) %>% 
    mutate(sig = padj < 0.05 & abs(lfc) > 1) %>% 
    ggplot(aes(x = lfc, y = -log10(padj), col = sig, size = baseMean)) +
    geom_point(alpha = 0.5) + 
    scale_color_manual(values = c("black","red")) +
    xlim(-7.5, 7.5) +
    ylim(0, 70) +
    coord_equal(ratio = (15/70)) +
    geom_hline(yintercept = -log10(0.05), col = 'blue', linetype = 'dashed') +
    geom_vline(xintercept = c(-1, 1), col = 'blue', linetype = 'dashed') +
    scale_size_area()
```

It is over-busy to plot all of the TSS labels on this graph, so here
are some ‘favorites’ labeled (based on genes that we saw above have
significant differences at the level of H3K27me3).

```
favorites = c("esg", "sna", "sisA", "grh", "wg", "eve", "ind", "hbn", "sisA", "amos", "ato", "sim", "wntD")

resZp %>% 
    filter(withTSS & !is.na(inDomain)) %>% 
    mutate(wT = case_when(
        tssName %in% favorites ~ tssName,
        !tssName %in% favorites ~ NA_character_
    )) %>% 
    mutate(sig = padj < 0.05 & abs(lfc) > 1) %>% 
    ggplot(aes(x = lfc, 
               y = -log10(padj), 
               col = sig, 
               size = baseMean, 
               label = wT)
           ) +
    geom_point(alpha = 0.5) + 
    scale_color_manual(values = c("black","red")) +
    xlim(-7.5, 7.5) +
    ylim(0, 70) +
    coord_equal(ratio = (15/70)) +
    geom_hline(yintercept = -log10(0.05), col = 'blue', linetype = 'dashed') +
    geom_vline(xintercept = c(-1, 1), col = 'blue', linetype = 'dashed') +
    scale_size_area() +
    ggrepel::geom_label_repel(show.legend = FALSE, 
                             box.padding = 0.3, 
                             max.overlaps = 1e+5, 
                             size = 2
                             ) +
    labs(title = "DESeq: RNAPol2 over zyg TSS in PcG Domains", x = "LFC [zld/wt]" )
```

```
## Warning: Removed 143 rows containing missing values or values outside the scale range
## (`geom_label_repel()`).
```

Most of the regions mentioned in the text have significant changes in
RNA Pol2. Snail continues to be expressed, albeit at a slightly lower
level of expression, in Zelda mutants (Nien PLoS Genet. 2011). Wingless
hasn’t been measured by in situ, but wingless is paused, and this lack
of a change could reflect loss of a signal in the gene body that we
aren’t picking up here. Grainyhead is not expressed robustly until
gastrula stages, so ‘it isn’t on yet’. I believe this is also the case
for esg. Homeobrain is a Bicoid target, so it may not care whether zelda
is around for expression, like other gap genes like Hb and Kr.

###### What is the magnitude of change for H3K27me3 for bins containing TSS that are zelda-dependent for RNA Pol2 recruitment?

If we take the set of TSS that have significantly reduced RNA Pol2 in
a zelda mutant, what is the distribution of log2(FoldChanges) in
H3K27me3?

```
downTSS = resZp %>% 
    filter(withTSS & lfc < -1 & padj <0.05) %>% 
    select(tssName)
downTSS = downTSS[,1]

tt = resZ %>% 
    filter(tssName %in% downTSS) %>% 
    mutate(domain = !is.na(inDomain)) %>% 
    mutate(wT = case_when(
        abs(lfc) > 1 ~ tssName,
        abs(lfc) <=1 ~ NA_character_
    )) %>% 
    group_by(domain) %>% 
    ggplot(aes(x = lfc, y = domain, label = wT)) +
    ggbeeswarm::geom_beeswarm(groupOnX = FALSE, alpha = 0.5) +
    geom_vline(xintercept = c(-1, 1), linetype = 'dotted', col = 'blue') +
    ggrepel::geom_text_repel(show.legend = FALSE, 
                             box.padding = 0.5, 
                             max.overlaps = 1e+5, 
                             size = 2, 
                             na.rm = TRUE, 
                             min.segment.length = 0.1, 
                             color = 'brown3'
                             ) +
    xlim(-3.5, 3.5) +
    coord_equal(ratio = 3/2) +
    labs(y = "PcG Domain Membership", x = "H3K27me3 LFC") +
    ggthemes::theme_clean()
tt
```

The plot above shows a number of things. For the set of zld-dependent
TSS within PcG domains, these are TSS that have statistically
significant losses in Pol2 recruitment that happen to fall within PcG
domains, and therefore are associated with some degree of H3K27me3.
Because this mark is theoretically associated with transcriptional
repression, one might expect that reduced gene activity (here estimated
by the recruitment of Pol2) would result in increased H3K27me3. But we
don’t see that except at the *zen* and *eve* loci. Instead
we see that most of these sites that lose Pol2 also lose H3K27me3,
almost as if Zelda is responsible for putting these sites into play and
enabling them to either recruit Pol2 or establish H3K27me3.

Another thing that is going on is that for TSS that require zelda for
Pol2 recruitment that sit outside of PcG domains, as reckoned in a
wild-type embryo, we do not see a tremendous acquisition of H3K27me3.
Why might we have expected this? Many of these sites harbor E(z) peaks
right at the TSS. Does loss of Zelda uncover some kind of latent
competency for these sites to acquire H3K27me3? Apparently not.

###### How many Zelda-dependent TSS outside of PcG domains overlap with E(z) peaks?

```
resZp %>% 
  filter(tssName %in% downTSS) %>% 
  mutate(domain = !is.na(inDomain)) %>% 
  group_by(domain, withEz) %>% 
  summarize(n = dplyr::n())
```

```
## `summarise()` has grouped output by 'domain'. You can override using the
## `.groups` argument.
```

```
## # A tibble: 3 × 3
## # Groups:   domain [2]
##   domain withEz     n
##   <lgl>  <lgl>  <int>
## 1 FALSE  FALSE     69
## 2 FALSE  TRUE     134
## 3 TRUE   TRUE      66
```

OK, all 66 of the Zelda-sensitive TSS within domains coincide with an
E(z) peak. This is reassuring. Of the 203 zelda-sensitive TSS outside of
PcG domains, 134 of them coincide with an E(z) peak (66%).

#### Additional numbers quoted in the results section

##### GAF

###### Number of bins within PcG domains for the GAF analysis:

```
sum(!is.na(bins2k$inDomain))
```

```
## [1] 3997
```

Overall within the starting bin set there are 3997 bins within
domains, but we filtered for low counts.

```
sum(!is.na(resG$inDomain))
```

```
## [1] 3883
```

There are 3883 bins in domains for the GAF analysis.

###### Number of bins that pass significance testing in the GAF analysis

At the level of bins, we say that “we observe only a limited number
of 2 kb bins within PcG domains with modest decreases in H3K27me3 signal
following GAF knockdown, and none of these pass significance
testing.”

```
resG %>% 
  filter(!is.na(inDomain)) %>% 
  mutate(sigDown = lfc < -1 & padj < 0.05) %>% 
  group_by(sigDown) %>% 
  summarize(n = dplyr::n())
```

```
## # A tibble: 1 × 2
##   sigDown     n
##   <lgl>   <int>
## 1 FALSE    3883
```

All of the down regions are not significantly down. There are however
some regions that have a p-value less than 0.05 and an lfc < 0, buf
fail to reach the magnitude of effect cutoff.

```
resG %>% 
  filter(!is.na(inDomain)) %>% 
  mutate(pDown = lfc < 0 & padj < 0.05) %>% 
  group_by(pDown) %>% 
  summarize(n = dplyr::n())
```

```
## # A tibble: 2 × 2
##   pDown     n
##   <lgl> <int>
## 1 FALSE  3847
## 2 TRUE     36
```

Here we have 36 bins that fall into this category.

In terms of regions that go “up”, we have a few that pass both
p-value and magnitude of effect thresholds:

```
resG %>% 
  filter(!is.na(inDomain)) %>% 
  mutate(sigUp = lfc > 1 & padj < 0.05) %>% 
  group_by(sigUp) %>% 
  summarize(n = dplyr::n())
```

```
## # A tibble: 2 × 2
##   sigUp     n
##   <lgl> <int>
## 1 FALSE  3875
## 2 TRUE      8
```

There are 8 individual 2kb bins that fall into this category.

And at the level of “any significant”?

```
resG %>% 
  filter(!is.na(inDomain)) %>% 
  mutate(pUp = lfc > 0 & padj < 0.05) %>% 
  group_by(pUp) %>% 
  summarize(n = dplyr::n())
```

```
## # A tibble: 2 × 2
##   pUp       n
##   <lgl> <int>
## 1 FALSE  3506
## 2 TRUE    377
```

There are 377 regions that have adjusted p-values below the
threshold, just not necessarily with a significant magnitude of
effect.

###### Runs of bins in the GAF experiment

We report the number of “runs” that we get by considering consecutive
bins with adjusted p-values less than 0.05:

```
runsG %>% 
  nrow()
```

```
## [1] 361
```

Overall, whether within PcG domains or outside of domains, there are
361 runs.

What is their size distribution?

```
range(width(GRanges(runsG)))
```

```
## [1]  2000 24000
```

How many are within Domains?

```
runsG %>% 
  filter(!is.na(inDomain)) %>% 
  nrow
```

```
## [1] 234
```

234 have overlap with a PcG domain. How many different domains are
represented in this list?

```
runsG %>% 
  filter(!is.na(inDomain)) %>% 
  group_by(inDomain) %>% 
  summarize(n = dplyr::n()) %>% 
  nrow()
```

```
## [1] 124
```

124 different domains contain one or more runs of 2 kb bins with
modest localized changes in H3K27me3 following GAF knockdown. However,
we need to check whether any of these contain both increases and
decreases in H3K27me3 signal.

```
runsG %>% 
  filter(!is.na(inDomain)) %>% 
  mutate(ups = meanLFC > 0, downs = meanLFC < 0) %>% 
  group_by(inDomain) %>% 
  summarize(boths = any(ups) & any(downs)) %>% 
  filter(boths)
```

```
## # A tibble: 2 × 2
##   inDomain                     boths
##   <chr>                        <lgl>
## 1 PcGD:chr2L:16405200-16490000 TRUE 
## 2 PcGD:chr3L:10286200-10334200 TRUE
```

2 domains contain both increases and decreases in signal.

The first one corresponds to a domain that contains *dac*. The
second one corresponds to a domain that contains a bunch of lncRNAs next
to an olfactory receptor. Neither of these are of particular functional
interest to this investigator.

How many Domains have net GAF-dependent reductions in signal?

```
runsG %>% 
  filter(!is.na(inDomain)) %>% 
  filter(meanLFC < 0) %>% 
  group_by(inDomain) %>% 
  summarize(n = dplyr::n()) %>% 
  nrow()
```

```
## [1] 17
```

There are 17 Domains with GAF dependent reductions in signal. How
many distinct runs comprise this set?

```
runsG %>% 
  filter(!is.na(inDomain)) %>% 
  filter(meanLFC < 0) %>% 
  nrow()
```

```
## [1] 23
```

23 runs total split over 17 domains have GAF-dependent reductions in
signal. How many are associated with a GAF peak?

```
runsG %>% 
  filter(!is.na(inDomain)) %>% 
  filter(meanLFC < 0) %>% 
  summarize(n = sum(withGAF))
```

```
##    n
## 1 19
```

There are 19 of 23 runs that contain a GAF peak. “The effect is
likely to be direct.”

How many gains in K27me3 do we see in GAF knockdowns?

```
runsG %>% 
    filter(!is.na(inDomain)) %>% 
    filter(meanLFC > 0) %>% 
    group_by(inDomain) %>%
    summarize(n = dplyr::n()) %>% 
    nrow()
```

```
## [1] 109
```

And what is the likelihood of this being a direct effect of GAF?

```
runsG %>% 
    filter(!is.na(inDomain)) %>% 
    filter(meanLFC > 0) %>% 
    summarize(n = sum(withGAF))
```

```
##     n
## 1 109
```

All of these runs have a GAF peak, so this is very likely to be a
direct effect of GAF.

##### Zelda

###### Number of bins within domains for this analysis?

```
resZ %>% 
  filter(!is.na(inDomain)) %>% 
  nrow()
```

```
## [1] 3746
```

###### How many show statistically significant reductions in H3K27me3?

```
resZ %>% 
    filter(!is.na(inDomain) & lfc < -1 & padj < 0.05) %>% 
    nrow()
```

```
## [1] 226
```

```
226/3746
```

```
## [1] 0.06033102
```

six percent of bins within domains show statisticially significant
reductions in H3K27me3. Not a huge amount, overall.

###### Howman bins within domains show increases in H3K27me3?

```
resZ %>% 
    filter(!is.na(inDomain) & lfc >1  & padj < 0.05) %>% 
    nrow()
```

```
## [1] 62
```

```
62/3746
```

```
## [1] 0.01655099
```

###### How many runs of bins are there?

```
runsZ %>% 
    filter(!is.na(inDomain)) %>% 
    nrow()
```

```
## [1] 546
```

###### What is their size range?

```
range(width(GRanges(runsZ %>% filter(!is.na(inDomain)))))
```

```
## [1]  2000 46000
```

###### How many runs show moderate to large increases in H3K27me3?

```
runsZ %>% 
    filter(!is.na(inDomain) & meanLFC > 0) %>% 
    nrow()
```

```
## [1] 301
```

```
301/546
```

```
## [1] 0.5512821
```

###### How many runs have mean LFC > 1?

```
runsZ %>% 
    filter(!is.na(inDomain) & meanLFC > 1) %>% 
    nrow()
```

```
## [1] 9
```

```
9/546
```

```
## [1] 0.01648352
```

###### How many runs have moderate to large reductions in H3K27me3?

```
runsZ %>% 
    filter(!is.na(inDomain) & meanLFC < 0) %>% 
    nrow()
```

```
## [1] 245
```

```
245/546
```

```
## [1] 0.4487179
```

###### How many runs show large magnitude decreases in H3K27me3?

```
runsZ %>% 
    filter(!is.na(inDomain) & meanLFC < -1) %>% 
    nrow()
```

```
## [1] 29
```

```
29/546
```

```
## [1] 0.05311355
```

###### How many PcG domains can the runs with increases be associated with?

```
runsZ %>% 
    filter(!is.na(inDomain) & meanLFC > 1) %>% 
    group_by(inDomain) %>% 
    summarize(n = n()) %>% 
    nrow()
```

```
## [1] 9
```

Nine domains have runs that show increases.

```
range(width(GRanges(runsZ %>% filter(!is.na(inDomain) & meanLFC > 1))))
```

```
## [1]  2000 34000
```

Ranging in width from 2k to 34k bp in size.

```
runsZ %>% 
  filter(!is.na(inDomain) & meanLFC > 1) %>% 
  select(inDomain, withTSS)
```

```
##                                             inDomain withTSS
## chr2L:7290001-7316000     PcGD:chr2L:7277400-7315000      wg
## chr2L:15332001-15334000 PcGD:chr2L:15308600-15342600     esg
## chr2R:9976001-9982000     PcGD:chr2R:9972800-9988800     eve
## chr2R:17804001-17838000 PcGD:chr2R:17808000-17834200     grh
## chr2R:23044001-23046000 PcGD:chr2R:23042200-23056200    <NA>
## chr2R:23618001-23638000 PcGD:chr2R:23618600-23635000    <NA>
## chr3L:1174001-1184000     PcGD:chr3L:1177800-1182600    bab2
## chr3R:21458001-21460000 PcGD:chr3R:21376600-21517800    <NA>
## chrX:18310001-18312000   PcGD:chrX:18240000-18319800    <NA>
```

Five of the runs/domains that show increases contain early zygotic
TSS. These correspond to wg, esg, eve, grh, and bab2.

###### How many domains are represented by runs with Zelda dependent decreases?

```
runsZ %>% 
    filter(!is.na(inDomain) & meanLFC < -1) %>% 
    group_by(inDomain) %>% 
    summarize(n = n()) %>% 
    nrow()
```

```
## [1] 29
```

There are 29 domains associated with zelda dependent decreases.

```
range(width(GRanges(runsZ %>% filter(!is.na(inDomain) & meanLFC < -1))))
```

```
## [1]  6000 46000
```

Ranging in size from 6k to 46k bp in size.

```
runsZ %>% 
  filter(!is.na(inDomain) & meanLFC < -1) %>% 
  filter(!is.na(withTSS)) %>% 
  select(withTSS)
```

```
##                             withTSS
## chr2L:15472001-15492000         sna
## chr2L:18592001-18604000        amos
## chr2R:20944001-20970000         hbn
## chr3L:8998001-9016000          Doc3
## chr3L:15022001-15052000         ind
## chr3R:8266001-8302000   ato, CG9626
## chr3R:13060001-13078000         sim
## chr3R:13286001-13304000        wntD
## chr3R:20836001-20850000     CG15696
## chrX:11320001-11326000         sisA
## chrX:17812001-17820000      CG12986
```

Twelve of these significantly down domains contains an early zygotic
TSS.

###### How correlated are changes between H3K27me3 and H2Aub in Zelda mutants?

```
comp = resZ
comp$ub = resZ2a$lfc
comp %>% 
  filter(!is.na(inDomain)) %>% 
  ggplot(aes(x = lfc, y = ub)) +
  geom_point(alpha = 0.25) +
    geom_smooth(method = "lm", se = FALSE, color = "red", linetype = 'dashed') +
  geom_abline(slope = 1, intercept = 0, col = 'grey', linetype = 'dashed') +
  coord_equal() +
  xlim(-5,5) +
  ylim(-5,5) +
  labs(x = "H3K27me3 lfc (zld/wt)", y = "H2Aub lfc (zld/wt)", title = "correlation between PcG modifications") +
  theme_bw()
```

```
## `geom_smooth()` using formula = 'y ~ x'
```

We can estimate whether the slope is significantly different than 1
by performing a t-test on the output of the linear fitting.

```
fit = lm(ub ~ lfc, data = comp %>% 
             filter(!is.na(inDomain)) 
         )
summary(fit)
```

```
## 
## Call:
## lm(formula = ub ~ lfc, data = comp %>% filter(!is.na(inDomain)))
## 
## Residuals:
##      Min       1Q   Median       3Q      Max 
## -1.67292 -0.16566  0.03313  0.20406  1.28930 
## 
## Coefficients:
##              Estimate Std. Error t value Pr(>|t|)    
## (Intercept) -0.155281   0.005437  -28.56   <2e-16 ***
## lfc          0.709356   0.009550   74.28   <2e-16 ***
## ---
## Signif. codes:  0 '***' 0.001 '**' 0.01 '*' 0.05 '.' 0.1 ' ' 1
## 
## Residual standard error: 0.3311 on 3744 degrees of freedom
## Multiple R-squared:  0.5957, Adjusted R-squared:  0.5956 
## F-statistic:  5517 on 1 and 3744 DF,  p-value: < 2.2e-16
```

The slope of the regression is 0.68, which is pretty far off from 1,
which would be expected if H2Aub and H3K27me3 responded identically to
loss of zelda.

```
slope_est = coef(fit)[["lfc"]]
slope_se = summary(fit)$coefficients["lfc", "Std. Error"]
t_val = (slope_est - 1) / slope_se
df = df.residual(fit)
p_val = 2 * pt(-abs(t_val), df)

p_val
```

```
## [1] 5.609845e-182
```

The t-test p-value for the slope (null hypothesis is = 1) is pretty
convincingly small, so we can reject the null hypothesis.

```
confint(fit)
```

```
##                  2.5 %     97.5 %
## (Intercept) -0.1659404 -0.1446216
## lfc          0.6906325  0.7280797
```

Another gut-check is to check whether the 95% confidence interval
spans 0 for the intercept and 1 for the slope. Nope.

There is at least a clear positive correlation between these two
marks in terms of log fold change comparing zelda and wild-type embryos.
What is the correlation coefficient?

```
comp %>% 
  filter(!is.na(inDomain)) %>% 
  summarize(cor = cor(lfc, ub, method = 'spearman'))
```

```
##         cor
## 1 0.7801611
```

```
sessionInfo()
```

```
## R version 4.0.3 (2020-10-10)
## Platform: x86_64-apple-darwin17.0 (64-bit)
## Running under: macOS Big Sur 10.16
## 
## Matrix products: default
## BLAS:   /Library/Frameworks/R.framework/Versions/4.0/Resources/lib/libRblas.dylib
## LAPACK: /Library/Frameworks/R.framework/Versions/4.0/Resources/lib/libRlapack.dylib
## 
## locale:
## [1] en_US.UTF-8/en_US.UTF-8/en_US.UTF-8/C/en_US.UTF-8/en_US.UTF-8
## 
## attached base packages:
## [1] parallel  stats4    stats     graphics  grDevices utils     datasets 
## [8] methods   base     
## 
## other attached packages:
##  [1] GenomicAlignments_1.26.0              Rsamtools_2.6.0                      
##  [3] BSgenome.Dmelanogaster.UCSC.dm6_1.4.1 BSgenome_1.58.0                      
##  [5] rtracklayer_1.50.0                    Biostrings_2.58.0                    
##  [7] XVector_0.30.0                        DESeq2_1.30.1                        
##  [9] SummarizedExperiment_1.20.0           Biobase_2.50.0                       
## [11] MatrixGenerics_1.2.0                  matrixStats_0.57.0                   
## [13] forcats_0.5.1                         stringr_1.4.0                        
## [15] dplyr_1.0.7                           purrr_0.3.4                          
## [17] readr_2.0.1                           tidyr_1.1.4                          
## [19] tibble_3.1.5                          ggplot2_3.5.1                        
## [21] tidyverse_1.3.1                       GenomicRanges_1.42.0                 
## [23] GenomeInfoDb_1.26.2                   IRanges_2.24.1                       
## [25] S4Vectors_0.28.1                      BiocGenerics_0.36.0                  
## 
## loaded via a namespace (and not attached):
##  [1] ggbeeswarm_0.6.0       colorspace_2.0-0       ellipsis_0.3.2        
##  [4] fs_1.5.0               rstudioapi_0.13        farver_2.0.3          
##  [7] ggrepel_0.9.1          bit64_4.0.5            mvtnorm_1.1-3         
## [10] AnnotationDbi_1.52.0   fansi_0.4.1            apeglm_1.12.0         
## [13] lubridate_1.7.10       xml2_1.3.2             splines_4.0.3         
## [16] geneplotter_1.68.0     knitr_1.30             jsonlite_1.7.2        
## [19] broom_0.7.9            annotate_1.68.0        dbplyr_2.1.1          
## [22] compiler_4.0.3         httr_1.4.7             backports_1.2.1       
## [25] assertthat_0.2.1       Matrix_1.2-18          fastmap_1.1.0         
## [28] cli_3.6.4              htmltools_0.5.2        tools_4.0.3           
## [31] coda_0.19-4            gtable_0.3.0           glue_1.4.2            
## [34] GenomeInfoDbData_1.2.4 ggthemes_4.2.4         Rcpp_1.0.7            
## [37] bbmle_1.0.24           cellranger_1.1.0       jquerylib_0.1.4       
## [40] vctrs_0.6.5            nlme_3.1-149           xfun_0.31             
## [43] rvest_1.0.1            lifecycle_1.0.4        XML_3.99-0.5          
## [46] zlibbioc_1.36.0        MASS_7.3-53            scales_1.3.0          
## [49] hms_1.1.1              RColorBrewer_1.1-2     yaml_2.2.1            
## [52] memoise_1.1.0          emdbook_1.3.12         sass_0.4.0            
## [55] bdsmatrix_1.3-4        stringi_1.5.3          RSQLite_2.2.1         
## [58] genefilter_1.72.1      BiocParallel_1.24.1    rlang_1.1.5           
## [61] pkgconfig_2.0.3        bitops_1.0-6           evaluate_0.14         
## [64] lattice_0.20-41        labeling_0.4.2         bit_4.0.4             
## [67] tidyselect_1.1.0       plyr_1.8.6             magrittr_2.0.1        
## [70] R6_2.5.0               generics_0.1.0         DelayedArray_0.16.0   
## [73] DBI_1.2.2              mgcv_1.8-33            pillar_1.6.3          
## [76] haven_2.4.3            withr_3.0.2            survival_3.2-7        
## [79] RCurl_1.98-1.2         modelr_0.1.8           crayon_1.4.1          
## [82] utf8_1.1.4             tzdb_0.1.2             rmarkdown_2.14        
## [85] locfit_1.5-9.4         grid_4.0.3             readxl_1.3.1          
## [88] blob_1.2.1             reprex_2.0.1           digest_0.6.27         
## [91] xtable_1.8-4           numDeriv_2016.8-1.1    munsell_0.5.0         
## [94] beeswarm_0.2.3         vipor_0.4.5            bslib_0.3.0
```
